# Genetic basis of traits and performance in UK silver birch

**DOI:** 10.1101/2025.07.04.662749

**Authors:** Rômulo Carleial, Phoebe Swift, Michael D. Charters, Domen Finzgar, Eve Anthoney, Elina Sahlstedt, Katja Rinne-Garmston, Jo Clark, Richard A. Nichols, Joan Cottrell, Richard Whittet, Richard Buggs

## Abstract

Knowledge of the genomic basis of phenotypes in forest trees lags that of other economically important organisms. We sequenced the genomes of 2054 silver birch trees from 20-year-old field trials at three locations in Britain. Each trial site contains trees from the same 29 source populations, and one trial site (Drummond) is outside the climate envelope of the source sites. We discovered many single nucleotide polymorphisms (SNP), and a structural polymorphism, associated with several growth traits. Using models trained with genome-wide SNPs we estimate the genomic breeding values of 148 plus trees. We discover 117 SNPs significantly associated with source-site environmental variables, but genetic offset estimates using these SNPs provide poor predictions of tree growth rates (Drummond excluded). They are outperformed in this task by offset estimates using thousands of non-significant SNPs. Such estimates for Drummond in current climates are higher than for source sites under a future worst case climate scenario.

## Main

Trees play key roles in sequestering carbon^1,2^, slowing biodiversity loss^3,4^ and providing raw materials for economic use^5^. Our understanding of the genetic basis of tree traits and adaptation is limited because, unlike model plant species and annual crops, the longevity and large size of trees make them hard to manipulate^6,7^. Methods for genomic analysis of hard-to-manipulate organisms have been developed for humans and can be applied to trees. These methods typically rely on large scale population sequencing^8,9^, which has only recently become affordable for trees. For some tree species, long-term forestry provenance trials exist where a diverse range of genotypes are grown in common environments; these are ripe for large scale resequencing analyses unforeseen when the experiments were set up^7,10–15^.

Silver birch (*Betula pendula* Roth.) has a broad native distribution throughout Europe and northern Asia and its subspecies, *szechuanica* (often known as *B. platyphylla*), extends south into China^16^. It is a pioneer species and is widely grown for forestry^17^. Britain represents the extreme western edge of its range, where *B. pubescens* (downy birch) is also common. Together, they occupy 18% of Britain’s broadleaf woodland coverage (Forest Research, 2023). Of the two, *Betula pendula* is preferred as a productive timber species, and is more genetically tractable as a diploid (*B. pubescens* is tetraploid). Several provenance trials exist for *B. pendula* in Britain, designed to test for local performance and inform future planting choices. Early results from one series showed that provenances from the north of Britain grow more slowly than those from the south, in all trial sites ^18^. Here, we sequence the genomes of trees in this trial series and conduct genome-wide association studies on a range of phenotypic traits, and genome-environment association studies. We use these to build models to predict trait variation which we apply to a set of trees previously selected for phenotypically superior growth and form. We also test the ability of genomic forecasting of adaptation – using varying numbers of SNPs – to predict the growth performance of the provenances at the trial sites.

## Results

### Genomic diversity of samples

We generated approximately 20X whole genome coverage in 150 bp paired-end reads for 2,200 trees. Of these, 2,054 were from three trial sites planted in 2003, each containing the same 29 provenances (Fig. 1a). A further 148 were plus trees, selected phenotypically for superior growth and form in 1995-2005 from northern Britain as sources of potential future germplasm for planting (Fig. 1b). Allele balance of the 2,054 samples revealed 64 putative *B. pubescens* tetraploids and 89 putative triploid hybrids (Supplementary Figs. 1 & 2). The relatedness matrix constructed with KING^19^ correctly identified five technical replicates but also revealed 10 accidental double samples. It also showed many first and second (kinship coefficient >= 0.0886) degree relatives within certain provenances (Supplementary Fig. 3), and 73 trees which were more closely related to trees from a different provenance than that which they were believed to originate from. Only 11 of these could be traced back to an identification error during sampling, while the remaining 62 did not follow a pattern consistent with tissue sampling errors. This suggests two, non-mutually-exclusive, possibilities: (1) mislabelling of samples during seed collection, seed processing, the nursery stage or trial establishment, or (2) historic planting of seeds at the sites of origin from a common source. Given this uncertainty, these 73 trees were excluded from genome-environment associations. For less sensitive analyses such as population structure, these 73 trees were retained, with the 11 sampling errors reassigned to their correct provenance. Technical replicates, double samples and polyploids were excluded. The sample size now comprised 1,873 provenance trial samples and 144 plus trees (2,017 in total) and ∼5.9 million high quality SNPs (for a density plot by chromosome see Supplementary Fig. 5). For population structure we used a reduced set of ∼1.3 million SNPs in approximate linkage equilibrium.

**Figure 1:**
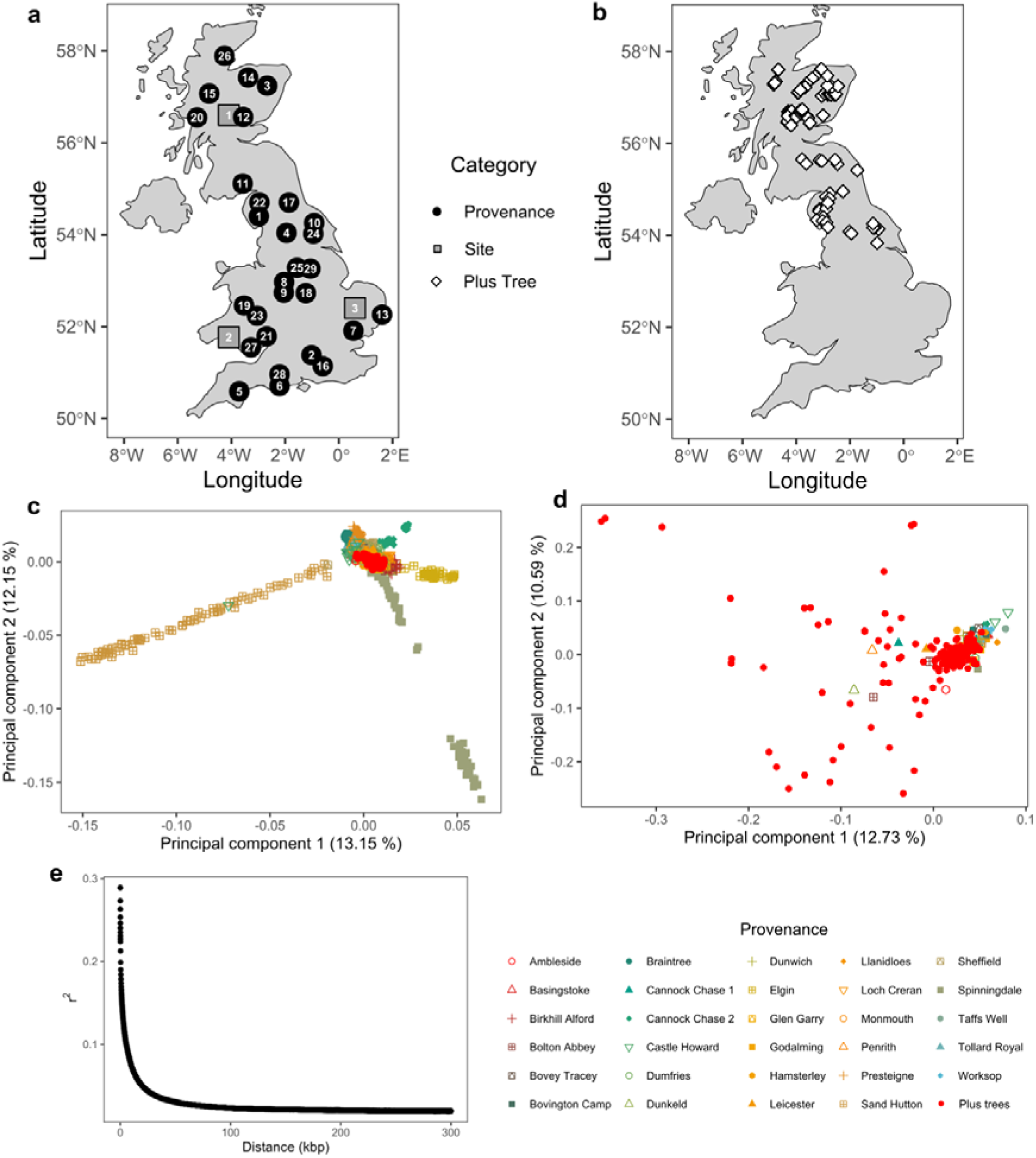
Location of trial sites, silver birch (*Betula pendula*) provenance source populations and selected plus trees. **>a,** Map of Britain showing locations of 20-year-old silver birch provenance trial sites (gray squares) at (1) Drummond Hill (Perthshire, Scotland), (2) Llandovery (Carmarthenshire, Wales), and (3) Thetford (Norfolk, England), and the 29 provenances (black points) common to each trial (provenance numbers match the alphabetical order in the bottom figure legend i.e. 1. Ambleside … 29. Worksop). **b**, Source locations of the Future Trees Trust silver birch plus trees (locations from https://ukfgr.org/research-trials/). **c,** Principal component analysis (PCA) of SNP data for 1,873 individuals across 29 British silver birch provenances and 144 plus trees. **d,** Genetic diversity of silver birch plus trees from the Glencorse clonal archive in Scotland with respect to the UK diversity. Only two randomly selected individuals for each of the 29 provenances were used for comparison. **e**, Whole-genome linkage disequilibrium decay in silver birch. Plus trees are highlighted in red in **c** and **d**, and provenances are colour and shaped coded (bottom).

**Figure 2:**
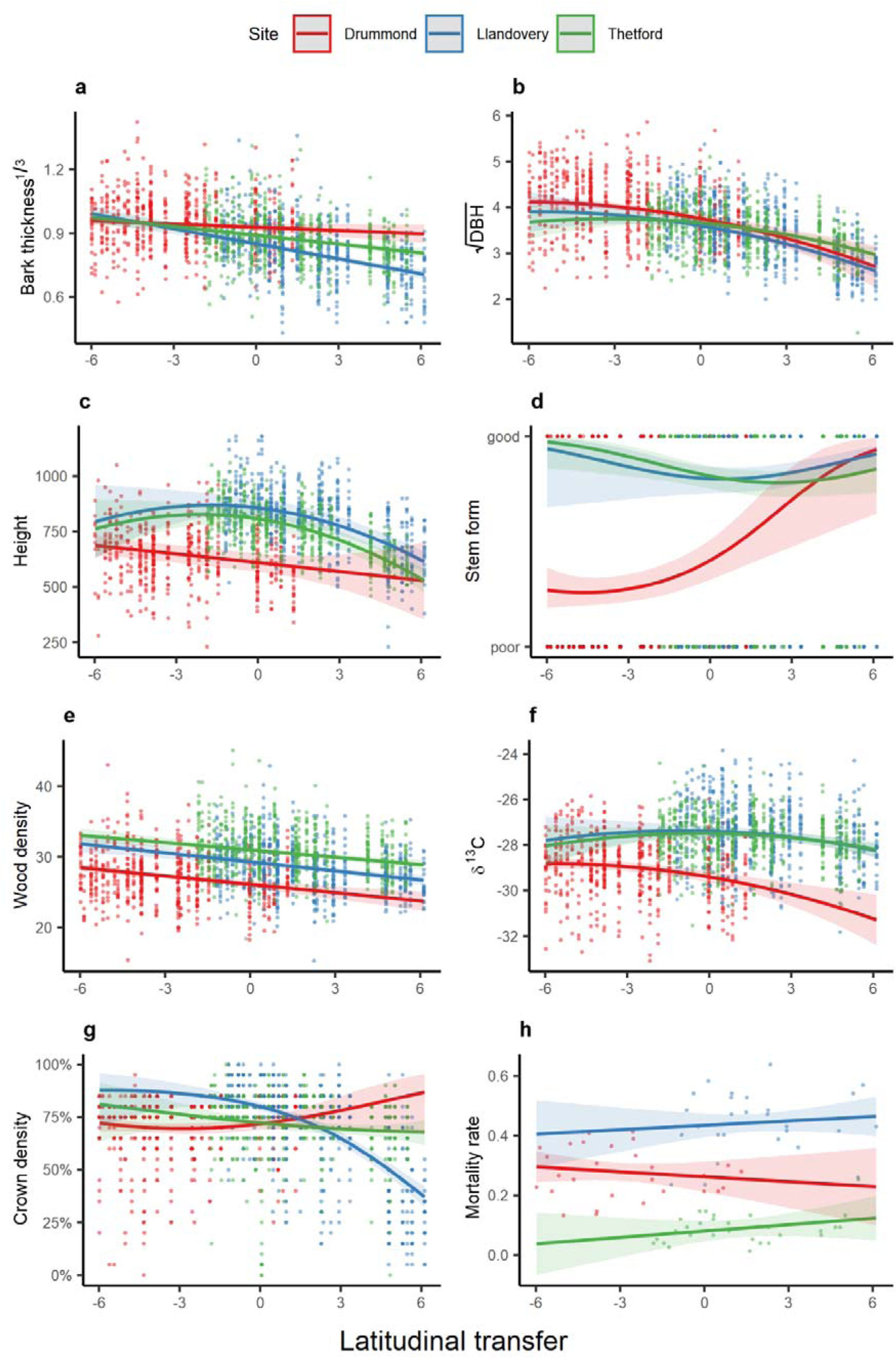
Quantitative and qualitative phenotypic trait values and their relationship with provenance latitudinal transfer and trial site. **a,** bark thickness, **b,** DBH, **c,** height, **d,** stem form, **e,** wood density, **f,** δ^13^C, **g,** crown density, and **h,** mortality rate. Slopes are colour coded by site (red = Drummond [56.6°], blue = Llandovery [52.6°], green = Thetford [52.4°]) and represent the marginal effects of statistical models, and shades represent the confidence intervals. Bark thickness and DBH were cube-root and square-root transformed, respectively. Traits **a-g** contain individual-level data, and trait **h** contains provenance-level data. Note: Site latitudes are provided within brackets. Latitudinal transfer = provenance latitude - site latitude.

Principal component analysis (PCA) of SNP data for these 2,017 trees (Fig. 1c, Extended Data Fig. 1) shows that much covariation is caused by intra-provenance relatedness. A PCA with only two individuals from each provenance showed a tight cluster for most samples but a greater spread for the plus trees (Fig. 1d).

Population structure inference using fastSTRUCTURE^20^ estimated K=10 as the simplest model for the 1,873 individuals in the provenance trials (maximum marginal likelihood), with K=29 optimal for explaining the structure in the dataset. Admixture plots (Supplementary Fig. 6) showed a similar pattern to the PCA. Removing first- and second-degree relatives and the 73 ambiguous trees (see above) reduced the dataset to 1,020 “unrelated” trees, which had optimal K of 1. Substructure in these 1,020 trees could now be explained by 17 components.

For the 1,873 trees, within-provenance genetic diversity calculated in STACKS^21^ decreased slightly with latitude (**H_E_**: F_1,27_ = 5.11, p = 0.032; for other metrics see Extended Data Fig. 2, Supplementary Fig. 7, Supplementary Table 2). Genetic differentiation (F_st_) between pairs of provenances was low, ranging from 0.0079 (Bovington Camp vs. Llanidloes) to 0.0294 (Spinningdale vs. Sand Hutton) (Extended Data Fig. 2h). Genetic diversity statistics for the 1,020 unrelated individuals are shown in Supplementary Fig. 8).

### Phenotypic traits

Mean values for 1,861 phenotyped provenance trial trees are shown in Figure 2 for height at 8 years, and for the traits measured after 20 years’ growth: diameter at breast height (DBH), carbon isotope composition (δ^13^C), wood density (measured indirectly as detrended debarked drilling resistance), bark thickness, crown density, stem form, and mortality. Correlations among these are shown in Supplementary Fig. 9. A latitudinal cline was seen for DBH, δ^13^C, height, bark thickness and wood density, suggesting southern provenances have higher values for these traits (Fig. 2). DBH, δ^13^C, crown density, stem form and height had significant quadratic relationships with latitudinal transfer, a pattern consistent with a degree of local adaptation (Fig. 2, Supplementary Table 3). For DBH, δ^13^C, height and crown density the local optima were south of the Thetford (England) and Llandovery (Wales) sites. For stem form, the optimum was north of Drummond (Scotland). We did not find a significant association between mortality rates and latitudinal clines.

### Genome-wide association study

We performed GWAS on the 1,861 trees. Using GLM, which does not account for relatedness and population structure, we found hundreds, sometimes thousands, of SNPs associated with most traits (Fig. 3). Genomic inflation was very high (Extended Data Fig. 3). Many top SNPs were shared among traits and these often showed latitudinal clines in their allele frequencies (Supplementary Fig. 10). A dense ∼400 Kbp peak in chromosome 6 was associated with multiple phenotypes (Fig. 3). PCAs summarising variation of SNPs contained in this peak show three clusters likely representing haplotypes (Fig. 4a, Supplementary Fig. 11), one of which is very uniform and common, and its presence follows a latitudinal cline (Fig. 4b). This pattern suggests a recently formed inversion^22^. Compatible with the expected reduction in recombination rates inside inversions, 78% of SNPs within this region (4,152/5,330) were monomorphic in putative inverted homokaryotypes compared to only 3.6% (190/5,330) in standard homokaryotypes, with mean minor allele frequencies of 0.012 and 0.222 respectively. Some genes inside the inversion have homologs previously implicated in cellular processes, apoptosis, and root morphogenesis (Fig. 4c, Extended Table 2). Taking trial site effects into account, individuals homozygous for the inversion had consistently lower DBH, wood density, and δ^13^C, compared to individuals in the same provenance but without the inversion (Fig. 4d).

**Figure 3:**
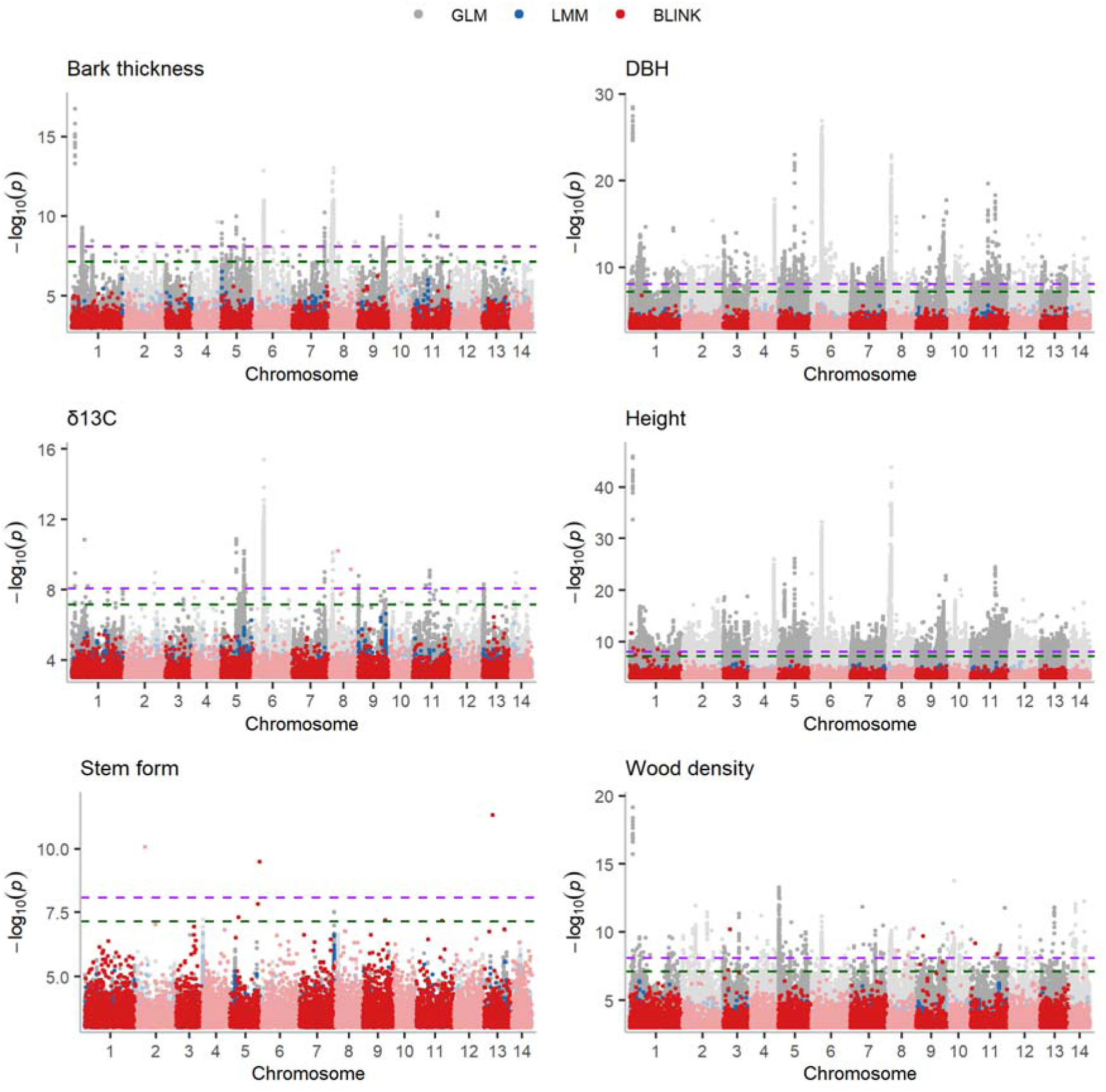
Genome-wide associations between SNPs and six traits of interest in silver birch (*Betula pendula*). Dots are colour coded according to the statistical method used i.e. dark grey = (generalised) linear model, red = linear mixed model, and dark green = BLINK. Dashed purple and red lines represent the Bonferroni and modified Bonferroni significance thresholds, respectively.

**Figure 4:**
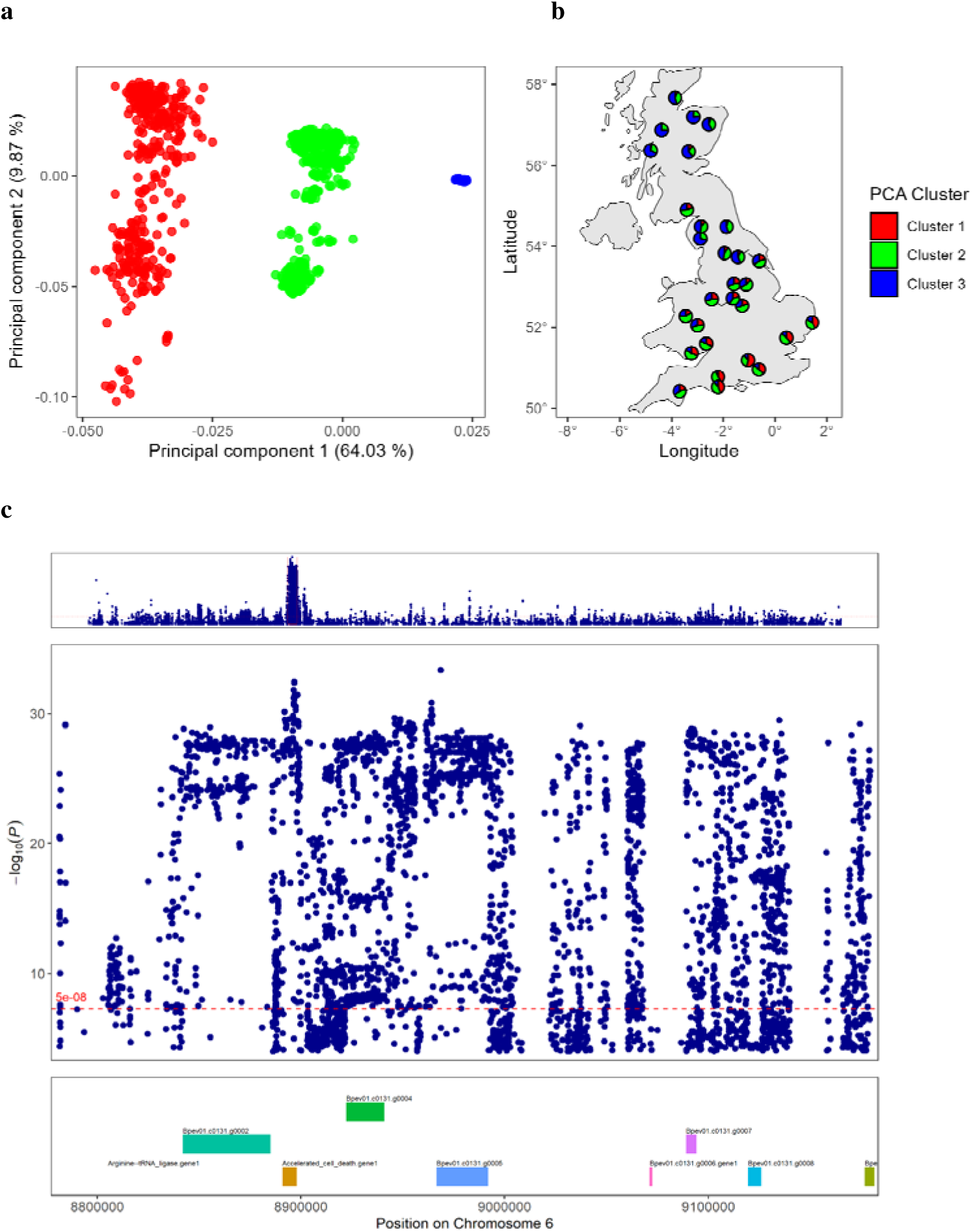
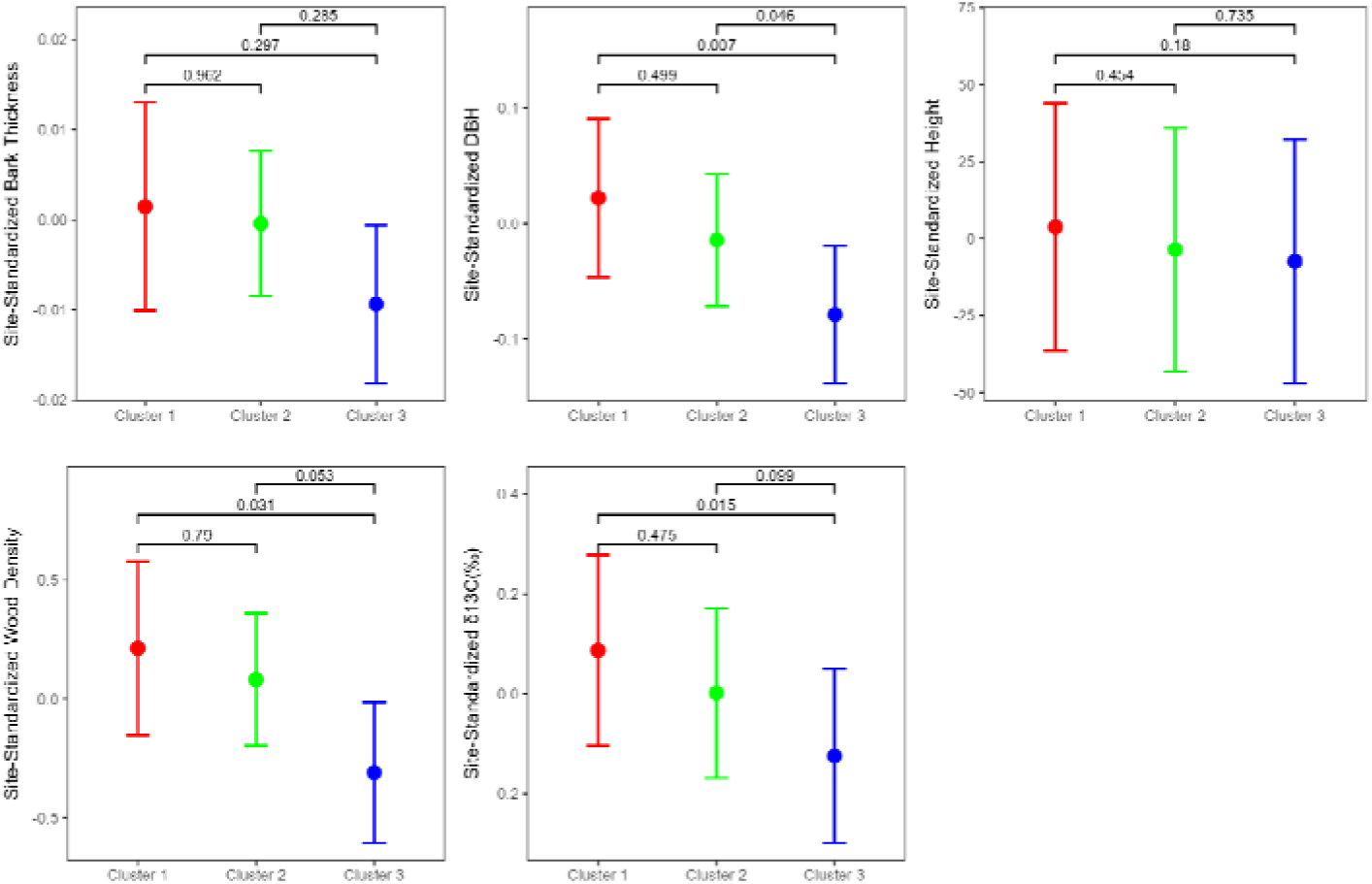
Genomic inversion within a ∼400 Kbp region of chromosome 6, its frequency across UK silver birch populations, and its effect on phenotypic traits. **a,** the first two principal components summarising variation in this region. The blue cluster contains 809 individuals; **b,** geographical location of provenances with the proportion of individuals belonging to each PCA cluster represented by a pie chart; **c,** GWA between SNPs in chromosome 6 and tree height using GLM. The main panel highlights SNPs located in the putative inversion, with genes found in the region represented by rectangles in the bottom panel. The putative function of these genes are provided in the Extended Table 2; **d,** Differences in marginal means between individuals with different inversion haplotypes. Phenotypes were standardised by the site mean grouped by provenance. P-values refer to post-hoc pairwise comparisons. Only sites with occurrences of all three haplotypes were included.

A GWAS using linear mixed models (LMMs) in LDAK^23^, fitting a genetic relatedness matrix (GRM) as a random effect (this minimises confounding caused by close kinship, the main source of structure in our dataset; see *Methods*), did not show genomic inflation (Extended Data Fig. 3) but found only one significant SNP. This was associated with δ^13^C (-log10(p) = 7.54) (Fig. 3), inside the inversion on chromosome 6, and 741bp away from a protein domain of unknown function (DUF1644), which has been associated with drought and salinity stress responses in rice^24^.

Another LMM incorporating genome by environment (GxE) interactions in GCTA^25^ showed some genomic inflation and identified 1,132 unique SNPs exceeding the significance threshold across five phenotypic traits (Supplementary Fig. 12). Of these, 24 were in protein coding regions (Extended Table 2), one of which was DUF1644. Of 1,061 height-associated SNPs, and 9 DBH-associated SNPs, 878 and 7 respectively were located within the chromosome 6 inversion. Significant for the GxE interaction term were 43 SNPs (associated with DBH (1), stem form (20), height (21), and δ^13^C (1)) (Supplementary Figure 12b). These were outside the inversion and three were within protein coding regions (Extended Table 2) and 37 were near genes (Supplementary Table 4).

We finally used the BLINK algorithm which has been shown to increase power compared to LMMs^26^. BLINK identified seven SNPs associated with height, 18 with wood density, three with δ^13^C, and seven with stem form, none of which inside the inversion (Fig. 3, Supplementary Table 4). These included SNPs located inside 9 protein coding regions (Extended Table 2).

### SNP-Heritability of traits

Using the REML method and the 1,861 trees (Table 1), we estimated high SNP-heritability for wood density (0.62 ± 0.06 standard deviation), height (0.62 ± 0.06) and DBH (0.53 ± 0.06); intermediate heritability for bark thickness (0.37 ± 0.06) and δ^13^C (0.32 ± 0.06), and low heritability for stem form (0.13 ± 0.05) (Table 1). Whilst inclusion of relatives is normally expected to bias heritability estimates upwards, REML on the 1,020 “unrelated” trees led unexpectedly to higher heritability estimates, which correlated with previous results (r^2^ = 0.95). QuantHer also gave higher estimates that were strongly correlated (r^2^ = 0.97). High heritability of traits despite few GWAS hits (after controlling for relatedness) suggests that these traits are under polygenic control. Thus, they may be suitable for genomic prediction.

**Table 1:**
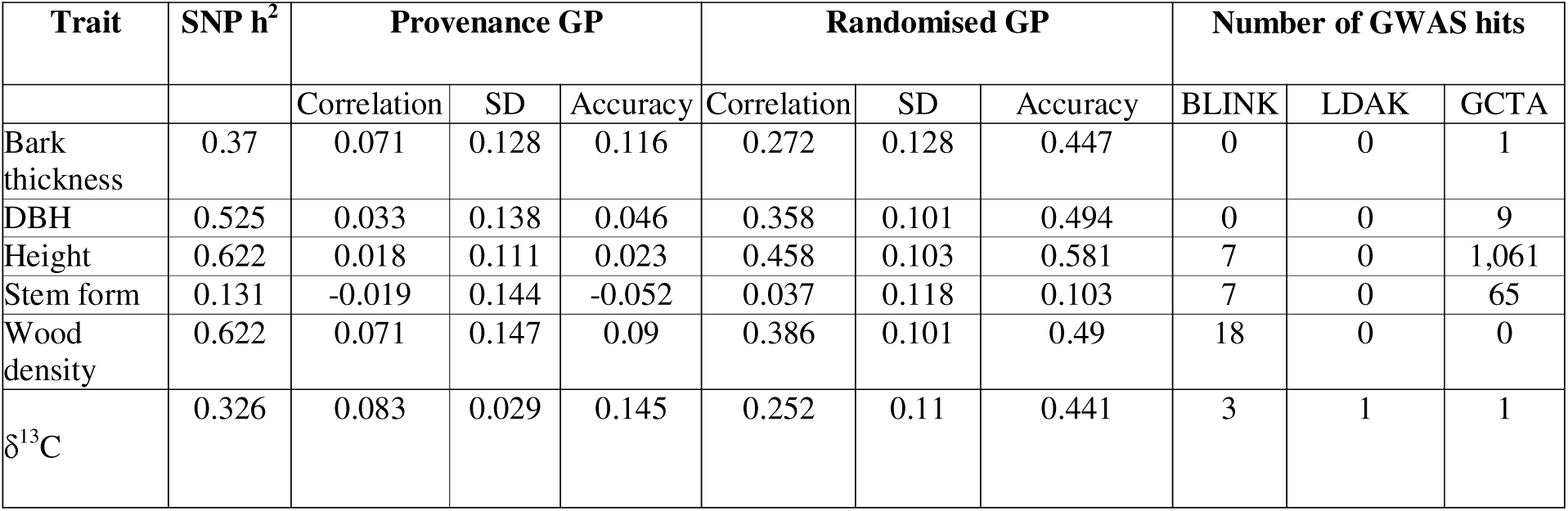
Summary statistics of heritability and genomic prediction results for traits in silver birch (*Betula pendula*). In the Provenance GP, individual provenances (n = 29) were omitted from the training set and used as the test set. In the Randomised GP, a random subset of individuals were omitted from the training set and used as the test set. Prediction correlation is the correlation between observed trait and genomic estimated breeding values. Accuracy is the correlation divided by the square root of the heritability.

### Genomic prediction

We trained and tested genomic prediction models using all ∼5.9 million SNPs on the 1,861 phenotyped trees from the provenance trials. When random subsets were used for testing, prediction accuracies ranged from 0.58 for height to 0.10 for stem form, with a mean of 0.43 across all traits (Table 1). Restricting the analysis to 1,020 “unrelated” trees, provided similar results (Supplementary Table 6). When all but one provenance was used as the training set and the other provenance as the test set, accuracies were much lower, ranging from 0.15 for δ^13^C to -0.05 for stem form, and averaging just 0.06 across all traits.

For each of the 148 plus trees we calculated GEBVs using a model trained on all 1,861 provenance trial samples. We compared the GEBVs of plus trees with provenance trial trees sourced from latitudes above 53.83 (the lowest latitude recorded for a plus tree in our dataset). The slope of a linear regression of latitude on GEBVs tended to be less steep for plus trees than for provenance trees for height, DBH, stem form and δ^13^C; i.e. the interaction term Latitude*Dataset was statistically significant (Extended Data Fig. 4 and Extended Data Table 3). This indicates that, compared to provenance trial trees from similar latitudes, plus trees tended to have better GEBVs for most traits. Of the 144 plus trees, 81 were homokaryotypes for the inversion on chromosome six, 54 were heterokaryotypes, and nine did not carry the inversion.

### Genome Environment Association

Environmental variables for the seed source and trial planting sites are summarised in Supplementary Figs. 13-15 and Supplementary Table 7. Analysing the 1,020 “unrelated” trees from the provenance trials with latent factor mixed models (lfmm2 in *LEA*^27^) at K=11 (Supplementary Figure 16c and 16d), 65 were significantly associated with mean temperature of the wettest quarter, 21 with slope, 15 with precipitation of the warmest quarter, 5 with isothermality, 4 with soil invertebrate taxa richness, 3 with mean temperature of the driest quarter, 3 with annual mean days of snow lying and 1 with northing (Extended Fig. 5). Nine of these 117 SNPs fell within the coding region of seven genes (Extended Table 2 and Supplementary Text 2). Two of these genes were also found in the GxE GWAS analysis.

This SNP set of 117 significant loci was reduced to only 22 following pruning for linkage disequilibrium (r=0.4, 50,000bp window) and a PCA of samples based on their genotypes at these 117 loci revealed distinctive clustering suggestive of association with structural variation (Supplementary Figure 21). Although lfmm2 has lower false positive rates than other GEA methods like lfmm^28^ and RDA^29^, we found that repeating our lfmm2 analyses with random permutations of environmental data allocated to seed source locations ^as^ ^in^ ^12^, resulted in similar numbers of significant hits (Supplementary Figure 22). Examination of allele clines for these SNPs suggested that they were often driven by a single seed source site at an extreme of the environmental distribution (Supplementary Figure 15 & 23). These combined observations led us to have little confidence in these 117 SNPs.

In order to assess the genomic variance attributable to environmental factors, we considered all 5.9M SNPs for which an effect size was estimated by lfmm2. We found that 19% of the relative variance in effect sizes could be attributed to precipitation seasonality, 12.6% to mean temperature of the driest quarter, 10.3% to precipitation of the warmest quarter, 8.5% to phosphorus availability, 8% with isothermality and lower values for other climatic factors (Supplementary Fig. 17a). Precipitation seasonality and mean temperature of the driest quarter showed the highest signal redundancy (Supplementary Fig. 17c). The majority (n = 10) of our environmental variables exceeded by more than two standard errors (SE) the average variance explained by 10 permuted dummy variables (3.5± 0.24%; Supplementary Figure 18), providing evidence of genuine genomic signals.

Two out of three soil variables (phosphorus availability and soil C:N,) explained moderate variation in effect sizes (8.5% and 5.9% respectively) (Supplementary Fig. 17a). Removing soil variables redistributed the explained variance among correlated predictors, resulting in a decrease in the relative contribution to total variance of isothermality and TPI, but an increase in unique variance suggesting they covary with soil variables (Supplementary Fig. 17). One soil variable (invertebrate taxon richness = 2.2%) and three terrain variables (northing = 2.8%, slope = 4.3%, easting = 3.6%) fell within or below one SE of the dummy average, so didn’t meaningfully differ from the null (Supplementary Fig. 17a and 18). Inclusion of first- and second-degree relatives made little difference to the pattern of these results (Supplementary Fig.19 and Supplementary Text 1).

### Predicting and testing genetic offset

Genetic offset — the degree of genetic mismatch between source populations and trial site environments — differed according to the SNP sets and methods we used. Alleleshift^30^ results using the 117 statistically significant GEA SNPs (22 after LD pruning) showed the Thetford site to have the largest genetic offset for almost all provenances (Extended Table 4 and 24a). This was likely caused by the model failing to predict a meaningful relationship between allele frequency and the environment at Thetford for ten provenances (GAM.rsq ∼ 0), with another 15 showing very low values (GAM.rsq =< 0.05), which inflated the allele shift estimation. Latitudinal transfer between seed source and trial sites had a positive correlation with this genetic offset measure (Supplementary Figure 25).

In contrast, AlleleShift^30^ results with 10,000 top GEA SNPs (∼2.8K after LD pruning) and LEA genetic gap and RONA^27^ results with ∼739K SNPs showed higher genetic offsets for most provenances at Drummond (Extended Table 4). The genetic gap and RONA genetic offset measures correlated with environmental dissimilarities between provenance source locations and trial sites, and Alleleshift results with ∼2.8K SNPs correlated with these somewhat too (Supplementary Figure 25). These three polygenic genetic offset methods were highly correlated (r > 0.85) (Supplementary Figure 25).

The three polygenic methods gave mean genetic offset predictions for the (now retired) worst-case climate change scenario (RCP8.5) at each provenance source location in 2070-2100 that were lower than the mean genetic offset at Drummond under current climatic conditions (Fig. 5). Alleleshift with 22 SNPs gave higher offset values for the current Thetford climate than for RCP8.5 at source locations (Fig.5c)

**Figure 5:**
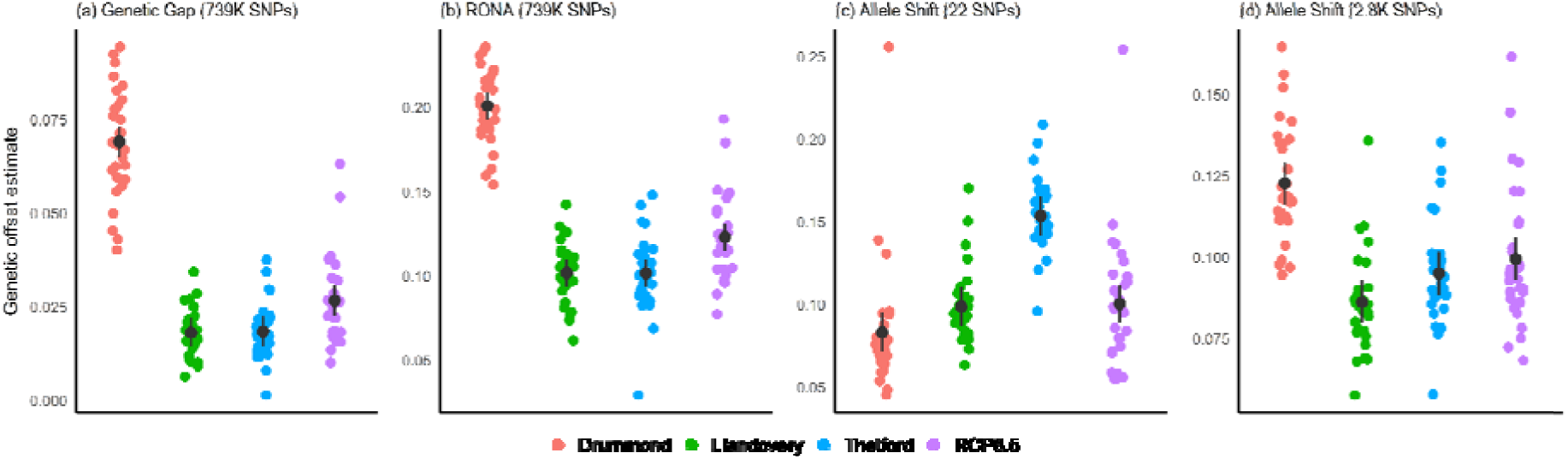
Genetic offset required for adaptation to three trial sites in the present and future climate at provenance source in 29 UK silver birch populations (*Betula pendula*). Each panel depicts a different genetic offset estimate: **a,** genetic gap, **b,** modified RONA, **c,** allele shift estimated with 22 statistically significant SNPs from GEAs, and **d,** allele shift predicted with ∼2.8k top GEA SNPs by p-value. Genetic gap and RONA were estimated using 738,833 genome-wide SNPs. Black dots and bars represent the predicted means and confidence intervals, and coloured dots show the raw values coloured by trial site (Drummond = red, Llandovery = green, Thetford = blue) or predicted future environment at the source (purple). SNPs used in the analyses were pruned to approximate linkage equilibrium from an original set of 117 (Allele Shift, 22 significant SNPs), 10k (Allele shift, top ∼2.8K SNPs) and ∼5.9M (genetic gap and RONA) SNPs, respectively.

Seeking to empirically validate our genetic offset predictions, we compared them to measures of phenotypic performance, particularly DBH (see Table 2; Fig. 6; Supplementary Table 8; Supplementary Figures 26-31). We focus here mainly on results for Thetford and Llandovery because these are within the climate envelope of the source sites (Supplementary Fig. 14), and had some signal of local adaptation in tree performance data (see above). We found negative associations (p < 0.05) between DBH and genetic offset for the three polygenic measures, but not for the results with 22 SNPs (Fig. 6). Largest R^2^ values and best fit by AIC scores were found for genetic gap. Height gave broadly similar results. Bark thickness showed a negative correlation with the polygenic genetic offset predictions at Thetford. Crown density showed a negative correlation with the polygenic genetic offset predictions at Llandovery. Wood density and δ13C did not correlate with any genetic offset measures.

**Figure 6:**
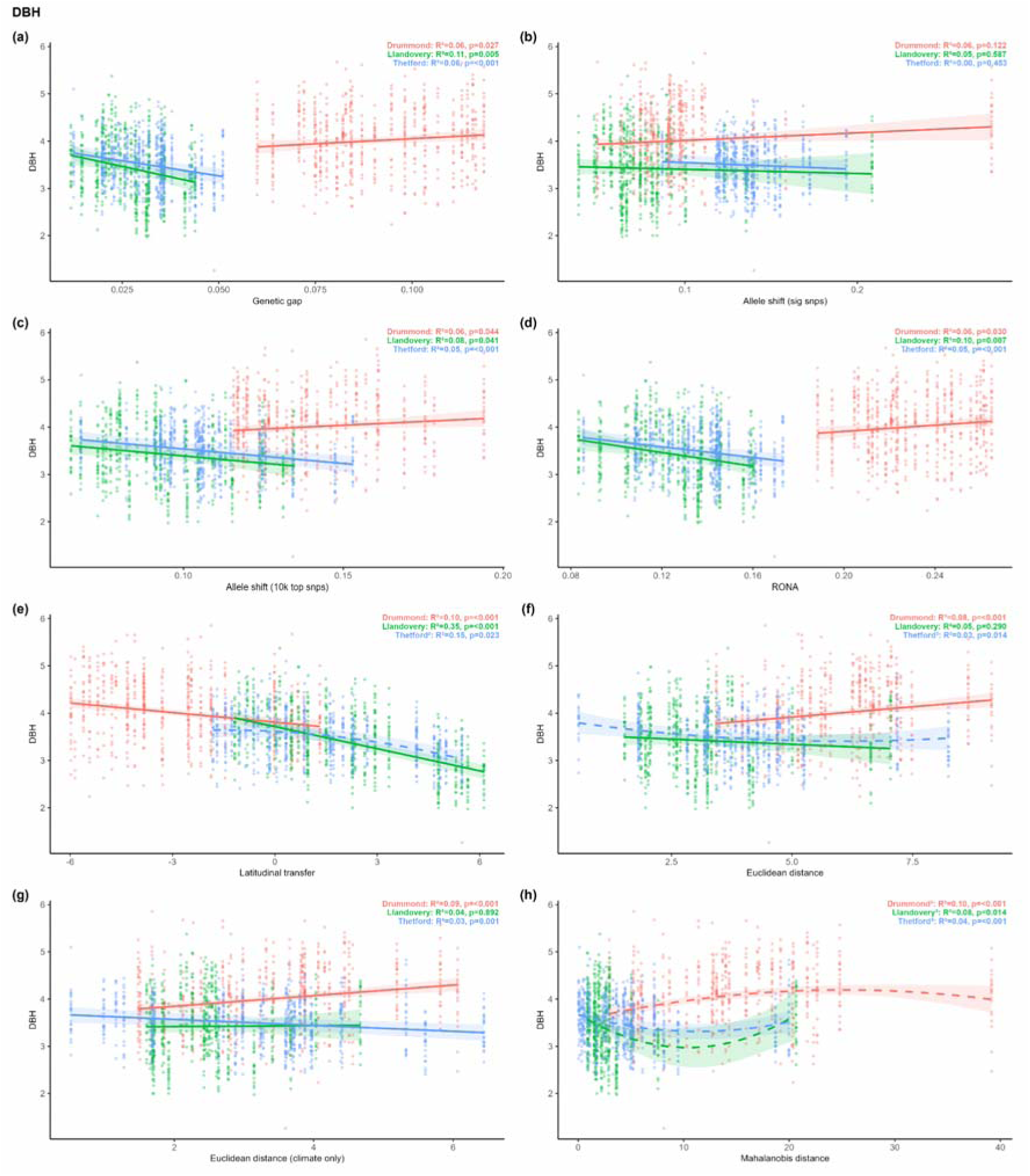
Relationship between diameter at breast height (DBH) after 20 years’ growth and (a-d) measures of genetic offset (e-f) measures of environmental distance. **a,** Genetic gap calculated with LEA using ∼739K SNPs, **b**, allele shift calculated with 22 SNPs, **c**, allele shift calculated with ∼2.8K SNPs, **d,** RONA calculated with LEA using ∼739K SNPs **e**, latitudinal transfer, **f**, euclidean distance with all environmental variables **g,** euclidean distance without soil variables, **h,** mahalanobis distance. Similar figures for other phenotypic traits can be seen in Supplementary Figs. 26-31. SNPs used in the analyses were pruned to approximate linkage equilibrium from an original set of 117 (Allele Shift, 22 significant SNPs), 10K (Allele shift, top ∼2.8K SNPs) and ∼5.9M (*LEA*) SNPs, respectively.

**Table 2:** Best simple environmental and genetic predictors of tree growth traits at each trial site according to model-estimated AICs. Predictors tested include the difference in latitude between source and planting site (Latitudinal transfer), environmental similarity between source and planting site determined by the (i) Euclidean distance in PCA space using all environmental, terrain and soil variables included in the GEA (EucDistance), (ii) only the climate variables (EucDistanceClimate), or (iii) the Mahalanobis distance (MahDistance) including all variables. Four metrics of genetic offset between provenance and trial site were calculated using (1) the AlleleShift package on 22 SNPs significantly associated with environmental variables (AlleleShiftSigSNPs), (2) the top 2.8K GEA SNPs ordered by p-values (AlleleShift2.8K), and using the LEA package on 738,833 unlinked SNPs to determine (3) the genomic gap, weighted by effect-size (GeneticGap) and (4) Risk Of Non Adaptedness (RONA). Results for all models can be found in Supplementary Table 8). SNPs used in the analyses were pruned to approximate linkage equilibrium from an original set of 117 (AlleleShiftSigSNPs), 10K (AlleleShift2.8K) and ∼5.9M (GeneticGap; RONA) SNPs, respectively.

| Site | Trait | Best environmental predictor | $r^2$ | AICc | Best genetic predictor | $r^2$ | AICc | Environmental-Genetic $\Delta\text{AICc}$ |
| --- | --- | --- | --- | --- | --- | --- | --- | --- |
| Thetford | DBH | Latitudinal transfer | 0.15 | 794.85 | GeneticGap | 0.06 | 843.64 | -48.79 |
|  | Height | Latitudinal transfer | 0.37 | 7689.42 | GeneticGap | 0.16 | 7743.62 | -54.21 |
|  | Wood density | Latitudinal transfer | 0.04 | 3463.25 | AlleleShiftSigSNPs | 0.02 | 3472.03 | -8.77 |
|  | Crown density | EucDistanceClimate | 0.46 | -1000.67 | AlleleShift2.8K | 0.14 | -1007.96 | 7.28 |
|  | Bark thickness | Latitudinal transfer | 0.07 | -1127.62 | AlleleShift2.8K | 0.02 | -1096.42 | -31.20 |
| | $\delta^{13}\text{C}$ | Latitudinal transfer | 0.02 | 1804.64 | RONA | 0.01 | 1808.80 | -4.17 |
|  | Stem form | MahDistance | 0.03 | 602.50 | AlleleShift2.8K | 0.09 | 597.73 | 4.77 |
| Llandovery | DBH | Latitudinal transfer | 0.35 | 908.82 | GeneticGap | 0.11 | 968.70 | -59.88 |
|  | Height | Latitudinal transfer | 0.32 | 7433.52 | GeneticGap | 0.08 | 7470.45 | -36.93 |
|  | Wood density | Latitudinal transfer | 0.07 | 3198.30 | RONA | 0.02 | 3213.94 | -15.65 |
|  | Crown density | Latitudinal transfer | 0.90 | -637.57 | GeneticGap | 0.75 | -585.56 | -52.02 |
|  | Bark thickness | Latitudinal transfer | 0.21 | -932.49 | GeneticGap | 0.06 | -893.91 | -38.58 |
| | $\delta^{13}\text{C}$ | Latitudinal transfer | 0.06 | 1977.49 | AlleleShift2.8K | 0.02 | 1985.89 | -8.40 |
|  | Stem form | Latitudinal transfer | 0.05 | 639.72 | RONA | 0.04 | 642.77 | -3.05 |
| Drummond | DBH | MahDistance | 0.10 | 1095.42 | GeneticGap | 0.06 | 1112.57 | -17.15 |
|  | Height | Latitudinal transfer | 0.07 | 6188.23 | GeneticGap | 0.02 | 6199.54 | -11.30 |
|  | Wood density | MahDistance | 0.07 | 3044.14 | AlleleShift2.8K | 0.01 | 3058.83 | -14.70 |
|  | Crown density | MahDistance | 0.26 | -679.73 | GeneticGap | 0.20 | -677.31 | -2.42 |
|  | Bark thickness | EucDistanceClimate | 0.03 | -788.15 | AlleleShiftSigSNPs | 0.02 | -786.00 | -2.15 |
| | $\delta^{13}\text{C}$ | EucDistanceClimate | 0.08 | 1789.83 | AlleleShiftSigSNPs | 0.05 | 1804.65 | -14.82 |
|  | Stem form | EucDistanceClimate | 0.10 | 658.32 | AlleleShiftSigSNPs | 0.12 | 659.81 | -1.49 |

However, latitudinal transfer correlated better with DBH and height at Thetford and Llandovery than the genetic offset predictions, with much lower AIC scores (ΔAICc > 30; Table 2, Supplementary Table 8). All traits showed a much stronger correlation with latitudinal transfer than any measure of genetic offset at Thetford and Llandovery, with the exceptions that crown density at Thetford was best described by allele shift with 2.8K SNPs and stem form was best described with genetic gap. At Thetford and Llandovery, environmental dissimilarity measures (Euclidian and Mahalanobis) performed less well than the polygenic genetic offset measures at predicting DBH. At Drummond, which is outside the provenances’ climate envelope, estimates of environmental dissimilarity were the best predictors of provenance performance for 6 of 7 traits.

## Discussion

Whole genome sequencing of over 2000 trees in a series of 20-year-old silver birch provenance trials allowed us to: (1) discard triploids, tetraploids and close relatives from analyses in which they could have contributed to spurious conclusions; (2) estimate the heritability of phenotypic traits, discover candidate genes associated with them and train polygenic genomic prediction models; (3) conduct in silico assessment of selected plus trees; (3) detect genome-wide variance associated with environmental factors; (4) estimate genetic offset at planting sites and show with phenotypic data that estimates based on many loci perform better than estimates based only on a few loci that are significantly associated with environmental variables; (5) show that trees survive and grow in conditions predicted to have larger genetic offsets than a worst case climate change scenario.

Our analyses were hampered by the fact that latitudinal clines are a persistent feature of the British silver birch study system. Many of the trait variables showed phenotypic clines, with southern provenances performing better at all sites even when there was some signal of local adaptation. Genome-wide within-provenance genetic diversity decreased slightly with latitude. In GWAS that did not account for population structure, many loci, and a large structural variant, that were significantly associated with growth differences showed latitudinal clines. In GWAS that did account for population structure, few SNP loci were found significantly associated with traits. Incorporating a GxE term in the GWAS led to the identification of three putative genes whose effects are dependent on the environment. GEA that accounted for population structure found few significant candidate loci, and genetic offset calculated with these could not reliably predict traits. Genome-wide signals from these GEA provided measures of genetic offset that were corroborated by performance data but were out-performed by latitudinal cline as a predictor of performance data.

These latitudinal clines are patterned by both drift and selection: drift because silver birch recolonised Britain after the last glaciation in a northward expansion^31,32^; selection because of clines in current environmental variables. This alignment makes it hard to disentangle the signal of adaptation from neutral signals in GWAS and GEA^33–35^. Our analyses that sought to include neutral population structure appear to have been overly conservative and eliminated loci that are truly associated with phenotypic trait variation. This is a well-known problem^33–35^ and might be overcome in future by more explicit models of the size of drift effects.

In GEA studies it is common to focus on statistically significant loci^12,36–38^, though these do not always perform much better than random exome loci^15^. We found 117 significant SNP loci but further investigation suggested they were unreliable. Our study is unusual in that we then used the effect sizes estimated for 5.9M genome-wide SNPs to show considerable effects of environmental variables on genomic variation. In doing so, we apply a logic that has been widely applied in the fields of genomic prediction^39^ and polygenic risk scores ^40^: although individual locus effects may be estimated imprecisely, genome-wide summaries are expected to be substantially more robust because shrinkage regularisation reduces variance in extreme coefficient estimates, while aggregation over many loci causes much of the remaining approximately unbiased estimation error to average out. The relative size for different environmental variables is particularly informative. We found temperature and precipitation variables had the largest associated genetic variance in UK silver birch, but soil phosphorous availability was also important. Soil is used by silviculturists as a major factor in deciding which species to use when establishing new woodlands ^41^, but has only occasionally been included in GEAs on trees ^42–44^.

Estimates of genetic offset are increasingly used to predict levels of adaptation to different sites in the present and future. We had the valuable opportunity to test such estimates using phenotypic data^45–48^. We found that the 117 significant SNPs from our GEA did not give genetic offsets that correlated with tree growth, but effect sizes from larger numbers of SNPs did. This gives support to the logic about aggregation over many loci outlined above. Our results also caution against over-interpretation of high genetic offset scores. Our genetic offset estimates for the Drummond population were higher than for extreme climate change scenarios at seed source sites, yet trees are still surviving and growing well at Drummond. Our study adds to the evidence that genomic forecasting methods may be unreliable in environments outside the sampled range^15,36,46–48^.

Our results can help inform the selection of seed sources for silver birch planting in Britain. The phenotypic dataset on its own suggests that, if planting for rapid growth, the ‘local is best’ rule only applies in the south of England and that to the north and in Scotland it is better to plant more southerly provenances. Our genomic data do not contradict this conclusion, as latitude was a better predictor of tree performance than any of our measures of genetic offset. However, if tree planters are seeking ecological restoration, caution may be needed. Growth traits are sometimes good fitness proxies^49^ but it is also possible that lower growth in the north reflects their conservative adaptation such that it could contribute to fitness if, for example, it protects against rare events such as unseasonably cold conditions^50^. It is possible that our genetic offset results provide better predictors of ecological adaptation than the growth data do alone, but we have no way of testing this hypothesis at present.

Our results can also inform tree improvement. We show that key growth traits have strong heritability, and thus could be improved by selection. Key candidate loci for growth could be selected individually, and the large inversion on chromosome 6 could be selected against. We also train genomic prediction models that are accurate -- as is typical^51–53^ -- within the same genetic background. Our genomic predictions suggest that previously selected plus trees are superior, and ongoing half-sibling progeny testing will further test this hypothesis. As Britain represents a small corner of the current global distribution of silver birch, it may contain variants that are rare elsewhere and could be of value in gene conservation or breeding programmes in other countries.

## Data availability

DNA read data generated by this project are available at https://www.ncbi.nlm.nih.gov/bioproject/PRJNA947136/

## Acknowledgments

We thank the Forest Research Technical Services Unit for work conducted in the field during this project and for long-term maintenance of the trial sites. We thank the owners of the trial sites for permission to access. This project was funded by the Department for Environment, Food & Rural Affairs via the Center for Forest Protection (Ref. CFP2202). KRG and ES acknowledge funding from the Research Council of Finland (343059). We thank Laszlo Csiba, Robyn S Cowan at the Jodrell laboratory and the Plant Health and Adaptation team at Kew for valuable advice on this study. We used Queen Mary’s Apocrita computing cluster (http://doi.org/10.5281/zenodo.438045) for data analysis, and their Research-IT team’s support is much appreciated.

## Author contributions

Fieldwork was conducted by DF and MC with assistance from Forest Research’s Technical Support Unit. DNA lab work was conducted by MC. Statistical analyses were conducted by DF, EA, RC, PS. Bioinformatic analyses were conducted by MC, RC, PS. GWAS and GP were conducted by RC. GEA was conducted by RC and PS. Carbon isotope analysis was conducted by KRG and ES. Artwork was produced by PS and RC. The project was conceived, proposed and managed by JC, RW and RB. All authors assisted in writing the manuscript.

## Online Methods

### Forestry provenance trials

The silver birch (*Betula pendula* Roth) provenance trials used in this study are described in detail by Lee et al.^18^, in which they were referred to as Series 2. Briefly, these trials were established from a collection of 50 catkins of *B. pendula* from 14-36 well-separated open-pollinated mother trees. Provenances were selected in woodlands that were considered to be autochthonous and located at 29 sites across Britain (Fig. 1a). The locations of these populations span all four Regions of Provenance (ROPs) in Britain and represent most of the 24 native seed zones therein. Seeds were collected in 1995 for the provenances in Scotland, Castle Howard and Sand Hutton, and in 2001 for the other provenances.

Prior to sowing, seeds from all mother trees within a provenance were bulked to capture the genetic diversity of the population. Seeds from all the sampled provenances were surface sown on compost held in Rootrainers™ in a glasshouse at Alba Trees’ commercial tree nursery in East Lothian, Scotland. The surface of the compost was then covered with a water-porous cloth to maintain moisture and avoid overheating or light saturation. When large enough to handle, seedlings were pricked out into individual containers and placed outdoors to harden off. Seedlings that had a morphology suggestive of *B. pubescens* (rather than *B. pendula*) were rogued out of the containers following the standard operating procedure at Alba Trees. The 1-year-old saplings were kept outdoors during the winter, before being planted out.

Planting took place in spring 2003 at three trial sites: Drummond Hill (Perthshire, Scotland), Llandovery (Carmarthenshire, Wales), and Thetford (Suffolk, England) (Fig. 1a). These sites were chosen to represent the different climatic conditions in which silver birch grows in Britain. Trials were planted in complete randomised blocks - a standard approach in forestry studies - with each site composed of three fully replicated blocks (Supplementary Fig. 4). Square plots for each provenance contained 25 trees (5 x 5) at Drummond and Thetford, and 36 trees (6 x 6) at Llandovery. Trees were planted at 2 x 2 m spacing. At Thetford only, a boundary of two edge trees was planted. Fences were erected around the perimeter of each site to minimise herbivore damage.

Upon revisiting these sites at the beginning of 2022, mature trials significantly differed in terms of level of maintenance, social structure, integrity of fencing, and density of undergrowth, which influenced the ease of orientation. The boundary fencing at Llandovery had been removed and the whole site was heavily overgrown with brambles. Several plot marking posts were missing and had to be reestablished. Block 3 at Llandovery was unavailable for sampling because it had been removed by the landholder. Both Llandovery and Drummond trials were established on a slight slope, resulting in flooding of several areas following periods of prolonged precipitation. Moreover, land erosion caused trees at Drummond to shift downhill, which caused an undesirable u-shaped formation at the base of the trunk of some trees. Additionally, slope and uneven terrain at Drummond caused by the use of continuous mounding for ground preparation resulted in poor row and column alignment of some trees, which caused issues during plot identifications. Conditions at the flatter, more uniform Thetford trial site were good.

Silver birch plus trees (*in-situ* phenotypic selections of individuals with outstanding form) were selected in Scotland and Northern England under the auspices of the British and Irish Hardwood Improvement Programme (latterly, Future Trees Trust) (Fig. 1b). Grafted trees made with scion material collected from 157 plus trees were planted at the Glencorse clonal archive site (55.85°, - 3.23°) in Scotland in 2010 (see: https://ukfgr.org/research-trials/). Each plus tree was represented by 1-3 ramets, with 401 trees in total arranged in 2 x 3 m spacing.

### Phenotypic assessments and sampling of genetic material

Material for DNA extraction from the provenance trials and the clonal archive of plus trees was sampled between May 30th and November 9th, 2022. At Drummond and Glencorse telescopic pole pruners were used to remove small branches from trees to provide a source of leaf samples, while at Llandovery and Thetford (where the leaf canopy was beyond reach) sterile scalpel blades were used to sample a small section of bark termed the phelloderm. Scalpel blades were cleaned with 95% EtOH between trees and discarded upon the completion of each provenance plot sampling. At Drummond and Thetford, 8 trees per provenance plot per block were randomly selected and sampled, totalling 24 trees per provenance per site. Due to the earlier removal of all trees in block 3 at Llandovery, 12 trees per provenance plot were randomly sampled in blocks 1 and 2 only. At Glencorse clone bank 148 grafted plus trees were sampled. The total sampling effort resulted in over 2,200 samples for DNA extraction, representing approximately 70 individuals per provenance distributed evenly across trial sites and blocks, and 148 plus trees from Glencorse. Leaf and phelloderm samples were immediately stored in silica gel desiccant with indicator beads. Dried samples were transported to RBG Kew for subsequent DNA extraction (see section DNA extractions). Sampled trees from provenance trials were tagged and their locations within the plots recorded to ensure that phenotypic assessments were conducted on the same individuals during subsequent site visits. Tree height was measured in 2008 when they were eight years old as outlined in Lee et al.^18^. We conducted additional survival and phenotypic measurements (i.e. DBH, mortality rate, stem form, crown leaf density, wood density, bark thickness, and δ^13^C, Extended Table 1) at all three provenance trial sites in 2022 and 2023. Diameter at breast height of 1.3 m (DBH) was measured in centimetres using a measuring tape. Survival of trees in each plot was assessed by counting surviving trees. To assess tree stem form, a simple ordinal scale was applied as follows: 1) a single, straight clear stem and leader; 2) a single stem and leader with kinks in the main stem and / or heavy branching; 3) neither 1 or 2, poor form. Stem form was evaluated by different assessors at each site. Crown leaf density, which can be used as a proxy for tree health (see below), was measured in August 2023 as the percentage of leaves still retained in the crown to an accuracy of 5% increments using the method described by Innes^54^. An indirect measure of wood density was obtained by resistance drilling. Bark-to-bark resistance traces were collected using the IML-RESI PD300 PowerDrill® (IML, Germany) positioned 1.3 m above ground level from the west side of the trunk. Mean drilling amplitude (power use by the drill) was estimated by adjusting bark-to-bark drilling profiles using an in-house processing application. The application removes low density bark from the profile and applies a linear correction function to account for the increase in friction (and therefore drill amplitude) as it passes through the stem. The summary score for a given tree was then the mean of all detrended amplitude values and is expressed as a percentage. Bark thickness on exit was measured as distance between the cambium position and the outer cork position when the drilling needle exited the tree stem (exit was used to reduce bias associated with the starting position of the drilling needle).

Carbon isotope composition (δ^13^C) was used to estimate intrinsic water-use efficiency (iWUE), which reflects the ratio of photosynthetic rate to stomatal conductance^55^. In August 2023, wood cores of the outside of the main stem were taken at 1.3 m above ground level from the west side of all genotyped trees using a 0.5 cm increment borer. The cores provided enough material for δ^13^C analysis of the tissue produced during the most recent growing season (2023, which was not a dry year). To facilitate the analysis of a large number of samples (∼2000) in a reasonable time frame, we targeted the most recent growth of the trees with the following analysis scheme. Using a scalpel, bark was removed from each core and the cores were reduced to 5 mm length and slid into the sample holders with the tissue from the most recent growing season facing upwards. The δ^13^C measurements for the tissue formed during the 2023 growing season were performed at the Stable Isotope Laboratory of Luke (SILL) of Natural Resources Institute Finland (Luke) using a laser ablation system coupled to an isotope ratio mass spectrometer (LA-IRMS). The novel approach of sampling the most recent wood growth without identifying annual tree rings was tested in a pilot study, where different lasing settings (laser energy, firemode, spot size, spot pattern, scan speed) were tested to achieve a penetration depth of the laser beam in order to target only the most recent growth ring of the trees.The following settings were used: Laser energy 40%, spot size 150 µm, laser pattern: line, He-flow 40 ml/min. With these settings the average penetration depth on the silver birch samples in the pilot study was 425 µm with standard deviation of 38 µm. Note that the laser ablates softer areas and areas closer to the sample surface more efficiently. Therefore, the study assumes that increment during the 2023 growing season would not be less than 0.425 mm. Details of the LA-IRMS methodology followed in this study are described in Saurer et al^56^.

### DNA extractions

Genomic DNA extractions were performed using an adapted CTAB protocol^57^. Dried plant material weighing 35-50 g was added to labelled Eppendorf tubes and homogenized using sterile ball bearings and a Spex® SamplePrep 2010 Geno/Grinder® set at 1500 strokes per minute for up to 3 minutes. TNE buffer was added to each tube and samples were kept on ice for 10 minutes. The supernatant was discarded after centrifugation and a solution of 800 µL of 2 X CTAB buffer, 120 µL 10% Sarkozyl solution and 20 µL Proteinase K was added. Samples were incubated in a water bath at 65°C for at least 2 hours, during which they were periodically inverted. To remove protein contaminants, multiple Sevag washes (24:1 chloroform:isoamyl alcohol) were performed, after which the upper aqueous phases were twice transferred to new tubes. Between washes, 5µl of RNAse (100 mg/ml) were added and samples were incubated in a water bath at 37°C for 30 minutes, to catalyse the breakdown of RNA molecules. DNA was precipitated in a solution of 5M sodium chloride and ice-cold isopropanol at -20°C overnight. Extractions were spun down and the resulting DNA pellet was cleaned with 75% and 95% ethanol respectively, before being vacuum centrifuged (Genevac™ miVac Duo Concentrator) and resuspending the pellet in 100 µl nuclease-free water. Quality control assessments were performed using Nanodrop (Thermo scientific, NanoDrop 2000) and agarose gel electrophoresis, and concentrations measured using a compact Quantus™ fluorometer (Promega Corporation).

### Whole-genome sequencing, SNP discovery and removal of polyploids

DNA samples were sent to Edinburgh Genetics Limited (UK) for 150 bp paired-end whole-genome sequencing using DNBseq technology on a DNBSEQ-T7 platform. Library preparations were performed using their egSEQ Enzymatic DNA library preparation protocol. We aimed to generate 20X coverage of the whole genome per sample. Raw reads were assessed for quality with FastQC v0.12.1 (https://www.bioinformatics.babraham.ac.uk/projects/fastqc/) and trimmed^58^ before being mapped to the silver birch reference genome^59^ using the BWA-MEM algorithm with the -M option. Single nucleotide polymorphisms (SNPs) were called using GATK v4.2.6.1 HaplotypeCaller and GenotypeGVCFs pipeline^60^.

To check for the presence of tetraploids (likely to be *Betula pubescens*) and triploids (likely to be *B. pubescens* x *B. pendula* hybrids) at the trial sites and FTT archives, we plotted allelic balance ratios at heterozygous sites^61^. Individuals with an approximate 0.5/0.5 allele balance ratio were considered diploids, whereas those with a ratio closer to 0.33/0.67 or 0.25/0.75 were considered triploids and tetraploids, respectively^62^. Preliminary analyses of allelic balance identified 2,042 diploids, 64 tetraploids, 89 triploids and five individuals of unknown ploidy (Supplementary Fig. 1). The vast majority of polyploid individuals belonged to southern provenances (Supplementary Fig. 2). No polyploid individuals were found among the plus trees. We assume the tetraploids are *B. pubescens* that were misidentified when the trials were established, and the triploids are hybrids between *B. pendula* and *B. pubescens*. Since GATK default options assume that individuals are diploid, we repeated the SNP calling pipeline after the removal of polyploids. These SNPs were then hardfiltered using GATK (parameters outlined in Supplementary Table 1). We used BCFTools^63^ to retain only biallelic variants, and PLINK 2 v20220603^64^ was used to filter alleles with counts lower than 3, with minor allele frequency < 0.05 and strongly deviating from hardy-weinberg equilibrium (p < 1^-50^). All SNP filtering steps can be found in Supplementary Table 1. This resulted in ∼5.9million SNPs, for which the density across the 14-chromosomes were plotted with the R package *CMplot*^65^ (Supplementary Fig. 5).

### Population structure and pairwise relatedness

To investigate population structure in silver birch, we first used all ∼5.9 million SNPs to calculate whole-genome linkage disequilibrium (LD) decay using PopLDdecay^66^. We then extracted SNPs in approximate linkage equilibrium using PLINK 2^64^ function --indep-pairwise 50 5 0.2. This reduced set was then used to investigate population structure of the diploid trees in the provenance trials and the plus trees in the Glencorse clonal archives. We performed principal components analysis (PCA) (Fig. 1c,d, Extended Figure 1) and estimated relatedness coefficients using the KING algorithm^19^ implemented in PLINK 2^64^. The former aimed at identifying strong population structure based on biogeography, and the second, at identifying technical replicates and potential sequencing/labelling errors and also to investigate fine-grained levels of population structure caused by families within provenances. The relatedness matrix identified five technical replicates that had been deliberately included (kinship coefficients of 0.44 ± 0.01 SD), and 10 additional pairs which were identified as accidental double sampling (kinship coefficients of 0.45 ± 0.01). These pairs were removed. PCAs and kinship coefficients were plotted in R v4.2.2^67^ using the package *ggplot2* v3.5.1^68^.

FastSTRUCTURE v.1.0^20^ was also performed on this SNP dataset to investigate the degree of admixture in each individual and to estimate the number of populations in the provenance trial dataset. We ran fastSTRUCTURE using a simple prior, fivefold cross-validation, and number of ancestral populations (K) ranging from 1 to 29 i.e. the number of provenances in the dataset. The fastSTRUCTURE’s <u>chooseK.py</u> and <u>distruct.py</u> scripts were used to respectively estimate the K value better able to explain the components of the data, and generate admixture plots for each K value measured (Supplementary Fig. 6). We used STACKS^21^ populations to estimate several metrics of genetic diversity for each provenance i.e. mean frequency of the major allele (P), nucleotide diversity (i.e. π = ∑2p(1−p)/N_sites_), observed heterozygosity (H_O_), expected heterozygosity (H_E_), inbreeding coefficient (i.e. F_IS_= 1 - H_o_/H_e_), and the degree of differentiation (i.e. pairwise F_ST_) between them (Extended Data Fig.2). STACKS was run on a 25% random subset of LD-filtered variable sites (341,081 SNPs) with the --smooth and --bootstrap flags. Finally, we investigated whether genetic diversity followed a latitudinal cline e.g. due to postglacial recolonisation, by regressing the provenance’s latitude on each genetic diversity metric (Supplementary Fig. 7). P-values were estimated by F-tests (Supplementary Table 2).

We repeated fastSTRUCTURE and STACKS analysis on a set of trees that excluded first and second-degree relatives (Supplementary Fig. 8).

### Provenance performance

To compare patterns of covariation among traits within the provenance trials, we calculated Pearson’s correlation coefficients between continuous phenotypic traits, based on 1,861 trees (Supplementary Fig. 9).

Next we investigated the relationship between individual tree performance at the three trial sites and latitudinal transfer i.e. site latitude - provenance latitude. Here, higher performance is defined as higher growth rates and denser crowns. Under the expectation of local adaptation, provenances closer to the trial site (i.e. latitudinal transfer ∼ 0) should perform better. We built linear mixed models (LMMs) in lme4 v1.1-36 ^69^ for the following continuous traits: DBH at age 19 (DBH); height in year 8 after the establishment of the trials (Height); average detrended microdrill resistance at age 20 (Wood density); bark thickness at the exit (Bark thickness); δ^13^C measured at age 20. Stem form, a categorical variable with tree levels, was transformed into a binary variable by grouping categories 1 and 2 (section *Forestry provenance trials*) as good form, and category 3 as poor form (i.e. 0 = poor form, 1 = good form). This binary variable was fitted in a generalised linear mixed model (GLMM) in lme4 v1.1-36 with a binomial distribution and logit link function. Crown density, a proportion variable ranging from 0 to 1, was fitted in a beta regression mixed model implemented in glmmTMB v1.1.8^70^. To avoid issues stemming from boundary values, 1 was adjusted to 0.99 and 0 to 0.01. For all variables, the trial site, provenance latitudinal transfer and their interaction were fitted as the main fixed effects. A quadratic effect of latitudinal transfer and its interaction with the trial site were also included. The number of neighbouring trees (discrete, 0-8) and whether the tree was planted at the edge of the trial site (binary, 0 = no edge, 1 = at the edge) were entered as covariates as these factors are known to affect phenotypic traits. We included the plot identity nested within the block identity as random effects to account for microenvironmental variation within sites (full model in lme4 syntax: Trait ∼ Site * (Lat_transfer + I(Lat_transfer^2)) + Edge + Neighbours + (1|Block/Plot)). Finally, provenance mortality rate was fitted in a linear model without covariates or random effects. This was necessary since we lacked information on both the exact number of dead and planted trees and their specific location on the trial site.

P-values were estimated using likelihood ratio tests (LRT) for generalised, linear and beta regression mixed models, and F-tests for linear regressions. Higher order interactions were dropped from the full model if p > 0.05. Significance tests were then repeated with the main effects. Model residuals were inspected using the DHARMa v0.4.6 (https://CRAN.R-project.org/package=DHARMa) package, which revealed that bark thickness and DBH had right-skewed distributions. A cube and a square root transformation were sufficient to correct for the skewness of bark thickness and DBH respectively.

### Genome-wide association study

We investigated associations between genetic markers and phenotypes using three different statistical models, namely (1) a (generalised) linear model (G)LM, (2) a linear mixed model (LMM), and (3) BLINK^26^. The first two model approaches were implemented in LDAK v6^23^, whereas BLINK was implemented in the R package GAPIT3^71^. For traits which showed evidence of non-normality in the previous section, we used the transformed phenotypic data instead. This is because linear models assume data are normally distributed, and fitting phenotypes with skewed distributions could lead to spurious associations. Similarly, stem form was transformed into a binary variable with both 1 and 2 scores considered as success (1) and form 3 considered as failure (0).

#### (G)LM

Linear models are the simplest approach to GWAS but do not directly account for population structure or relatedness. The model formula can be summarised as:

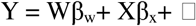

where Y is a vector of phenotypes, W is a matrix of covariates and β_w_ the vector of the unknown fixed effects, X is a matrix of genetic markers, β_x_ are its additive genetic effects, and is a vector of the residual errors. For continuous traits, the model is a simple linear regression whereas for binary traits, a logistic regression is fitted. These models were run without correction for population structure e.g. by including PCs as covariates, since mixed model approaches (see below) may increase the occurrence of type II errors^72^. (G)LMs served as baselines to which the other models were compared.

#### LMM

Linear mixed models (LMMs) control for subtle population substructure and relatedness by fitting a genetic relatedness matrix (GRM) as a random effect in the model. This is recommended since our dataset contains high levels of relatedness within provenances, which could lead to spurious associations if not controlled for^73^. Most of our traits were continuous variables, with the exception of stem form, which was binary. While logistic regressions are more appropriate for binary traits, linear mixed models are a close approximation, particularly when the data lack strong population stratification^74^. Linear mixed models can be described by the formula:

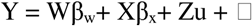

where Y is a vector of phenotypes, W is a matrix of covariates, β_w_ is the vector of the unknown fixed effects, X is a matrix of genetic markers and β_x_ are its additive genetic effects, Z is an incidence matrix that relates observations to random effects, and u are the random effects which follow a multivariate normal distribution 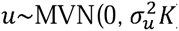, where K is the GRM and is a vector of the residual errors, which are also assumed to have a normal distribution 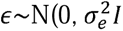, where I is the identity matrix. The GRM generated in LDAK v6 has additional assumptions that take into account the effects of LD and MAF on SNP heritability^23^. For example, estimated heritability can be inflated due to multi-tagging in LD regions, so we manually set 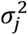 to 0 if SNP_j_ had high correlation (r^2^ > 0.98) with SNP_j’_ and 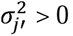. Moreover, because SNPs with low MAF are expected to contribute less to heritability, LDAK weights SNPs as a function of MAF such that:

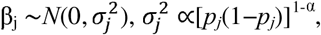

where β_j_ is the effect of SNP_j_ and 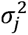 its variance, p is the MAF and α is the power scaling parameter. While a power of -0.25 has been shown to be a good approximation for many human traits^23^ we set α at 0 since it is the most commonly used parameter value in crop and animal research^75^.

#### GCTA

Genome-by-environment (GxE) associations are less often incorporated into GWAS, mostly due to the high power required to identify interactions robustly. Due to our moderately large dataset, we implemented LMMs including GxE effects using the software GCTA v1.95.1^25^. A sparse GRM was generated using the --make-bK-sparse 0.05 flag, and models were run using the --fastGWA-mlm flag. Since GCTA does not accept discrete environmental variables i.e. site identities, we implemented a PCA including 40 climatic, soil, and terrain variables from all three trial sites (see *Genome environment association* below for more information on these variables) and used the first PC as a continuous environmental variable summarising the main axis of variation between sites. Effect sizes and p-values were extracted for three hypothesis tests derived from the same model: the main genetic effect (P_G), the genotype-by-environment interaction effect (P_G_by_E), and a joint 2-degree-of-freedom test combining both (P_2df). We focused primarily on P_G_by_E to identify SNPs whose effects were environment-dependent. Any SNP that deviated more than 0.1 in frequency between sites was excluded from results, as this difference may be the consequence of early selection at the trial sites or variation in sample size, which may cause false positives.

#### BLINK

In order to reduce false negatives in linear mixed models and increase power, Huang et al.^26^ developed BLINK, a multivariate GWAS model which takes into account linkage disequilibrium between markers. In the first step, the method ranks markers based on p-values, with the most strongly associated marker serving as the reference. Any markers in LD with this reference marker are then removed. The process repeats iteratively using a Bayesian information criteria optimizer, selecting the next most significant marker as the new reference and eliminating markers in LD, until no further markers can be excluded. It also iteratively includes associated markers as covariates when testing other markers, thereby reducing confounding effects from relationships among individuals. Contrary to LMMs, BLINK does not require a GRM.

#### Covariates

For each model (i.e. (G)LM, LMM, BLINK, and GCTA) and phenotype, we included data relating to the position of each sampled tree in the trials relative to both the edge of the site (binary), the number of neighbouring trees (discrete), and the plot and block identities as covariates (see Provenance performance section). The former two were included to account for border effects and degree of resource competition respectively. Plot and block identities were included to account for micro-environmental variation within sites. Except for the GCTA model, dummy variables representing each of the three trial sites were also included as covariates to control for site specific effects.

#### Manhattan plots and gene annotations

We plotted p-values as Manhattan plots and Quantile-Quantile (QQ)-plots for each environmental variable using the *topr*^76^ R package. For the significance threshold we used a modified Bonferroni cutoff, which divides 0.05 by the number of haplotype blocks instead of SNPs. These were used to minimise stringency of the regular Bonferroni, since not all SNPs are independent due to linkage disequilibrium, but still be more stringent than false discovery rates. Haplotype blocks were determined using PLINK 1.9 v170906^77^. The standard Bonferroni threshold was added to Manhattan plots for reference. Finally, we searched for putative genes within 5kb intervals of statistically significant SNPs using a custom search function in the R v.4.2.2 environment^67^. The function uses the gene position information in the silver birch gene annotation file^59^ to search for genes within 5kb upstream and downstream of the target SNP. If the function fails to assign any gene to the SNP within that distance, it repeats the search in incremental 5kb intervals until a gene is found. The function will then move to the next SNP until all selected SNPs are ascribed a putative gene. If more than one gene is found within the 5kb interval, all are reported in the output file.

### Investigating an inversion on chromosome 6

GWAS analyses using the GLM method revealed a dense (∼400kbp) peak on chromosome six (Fig.3, Fig.4c). We investigated genomic variation in this region by PCA in PLINK 2^64^. PCA plots revealed three haplotypes (Fig.4a), whose abundance within provenance was plotted as pie charts in the UK map (Fig.4b) using the sf v1.10-16^78^, scatterpie v0.2.4 (https://CRAN.R-project.org/package=scatterpie), and ggplot2^68^ R packages. Since controlling for population structure in the GWAS eliminated significant associations in this region (Fig. 3), and given that both haplotype abundances and phenotypic traits showed clear latitudinal clines (Fig. 2, Fig. 4b), we compared differences in phenotypic traits between haplotypes using a different approach. To help eliminate site and latitudinal effects, traits were standardised by subtracting the mean value of each trait within the combination of provenance and site. The adjusted traits were then used in a linear mixed model with the haplotype identity as the predictor, edge and neighbours and covariates (see Provenance Performance section), and the block and plot identities as random effects. Models were implemented in the lme4 package v1.1-35.1^69^. Post hoc pairwise comparisons between haplotypes were then conducted using estimated marginal means (EMMs) in the emmeans v.1.10.0 package (<https://CRAN.R-project.org/package=emmeans). Two other approaches namely, 1) using raw phenotypic traits and controlling for site and provenance effects by fitting both as random effects, or 2) using propensity score matching implemented in the Matchlt package^79^, gave qualitatively similar, but less conservative, results, so are not reported.

### SNP-heritability of phenotypic traits

We estimated the SNP heritability of each phenotype to investigate their suitability for breeding programs. Most SNP heritability tools require that samples be unrelated to minimise the confounding effects of the common environment (i) and non-additive genetic effects (ii)^80^. This poses a problem to our dataset, which contains provenances with high degree of relatedness among individuals (see Results). While (i) is not applicable to provenance trials, since all individuals are raised in a similar environment, we cannot rule out that (ii) would inflate estimates. We therefore used three approaches to estimate SNP heritability. In the first approach (1), we used the KING software^19^ to identify and exclude individuals related by 2nd or 1st degree relatedness. We then regressed the phenotypes of the remaining individuals on the genomic relatedness matrix generated in the section above (GWAS, Equations 1 & 2) using a linear mixed model with restricted maximum likelihood (REML) in LDAK v6^23^. In this approach, the GRM is fitted as a random effect to estimate the variance components that maximise the REML. Since filtering related individuals in approach (1) decreases the sample size by approximately half, which can increase the mean squared error and therefore reduce the confidence in the estimates, we repeated (1) but this time including all individuals (approach 2). Finally, in the third approach (3), we estimated SNP heritability using the –quant-her and –tetra-her functions in LDAK v6. These functions use the coefficient of relatedness between pairs of close relatives to estimate the heritability of continuous and binary phenotypes, respectively^81^. In both methods we included the covariates described above in the model, namely, the dummy variables representing each trial site, the tree plot and block identities, the number of neighbours surrounding a tree and whether a tree was located at the edge of the trial site. The variance explained by these covariates was discounted from the final heritability estimates.

### Genomic prediction

To test whether we could accurately predict phenotypic traits using genetic markers, we fitted all ∼5.9 million SNPs in elastic net (ENET) prediction models using LDAK v6^23^. ENET is a penalised regression which combines the ridge and LASSO penalties^82^ and can therefore capture both sparse (few large-effect SNPs) and polygenic (many small-effect SNPs) architectures, thereby offering improved prediction accuracy, especially when the underlying genetic architecture is unknown. In the context of genomic prediction, the formula can be expressed as follows:

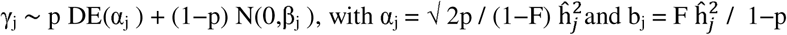

where DE(α_j_) stands for a double exponential distribution with rate α_j_. The relative contributions of the double exponential and normal distributions are regulated by the p and F parameters, respectively^83^. Suitable prior parameters were chosen by cross-validation.

Since the degree of relatedness between individuals in the training and test sets have been shown to heavily influence prediction accuracies, we implemented two approaches for genomic prediction (GP). Firstly, we trained models using a leave-one-provenance-out approach. Models were trained with one provenance excluded from the training set, and models were then used to predict the phenotypes of the individuals from the excluded provenance. We reasoned that genomic prediction accuracies for these models should be low, since the average relatedness between individuals in the training and test sets is also low. Secondly, we used a classical cross-validation method in which individuals were randomly ascribed to the test set until all individuals were included in both sets. Since the sample size of the test and training sets can affect accuracy^84^, randomised test sets contained a similar number of individuals found in the 29 provenances i.e. the 1,861 individuals were equally distributed into 29 sets.^84^ We compared the mean accuracy across the 29 models generated for each approach. Accuracy of predictions was determined by the correlation between observed phenotypes and genomic estimated breeding values (GEBVs), divided by the square root of the heritability of approach 2 in the section above.

Finally, we trained an ENET model for each phenotype including only 1,861 provenance trees to predict the GEBVs of plus trees from the Glencorse clonal archives. This allowed us to investigate whether subjective assessments of tree quality used on plus tree selection correlate well with our predicted genetic merit of these trees. The GEBV for plus trees was compared to the GEBVs estimated for the provenance trees in the randomised cross-validation approach as described above.

### Genome-environment association studies (GEAs)

Genome-environment association (GEA) studies were performed using a Latent Factor Mixed Model (LFMM). Such models offer a simple and quick approach to assess locus-specific linear associations for each environmental variable independently, after first fitting a multivariate model of all environmental variables simultaneously^85^. LFMMs allow neutral genetic structure to be accounted for through latent factors which represent hidden sources of population structure, removing the requirement to first identify loci under neutral genetic structure^28,37^. This method was shown to have low false positive rates and was successfully used in *Fagus*, *Alnus,* and other tree species^86–89^.

#### Environmental Variable Selection

Nineteen 1 km-resolution climatologies were downloaded from version 2.1 of the Climatologies at High Resolution for the Earth’s Land Surface Areas (CHELSA) global database^90^. Environmental variables were annual averages between 1981-2010, and were derived from temperature (n = 11) and precipitation (n = 8) records, reflecting annual trends, seasonal variation, and climate extremes. A further four datasets for the period 1981-2010 were downloaded from www.catalogue.ceda.ac.uk which detailed ground frost, snow lying, wind speed, and sunshine duration at 1 km resolution^91^. Altitude data were used in the R package Raster v.3.6.26^92^ to extract terrain characteristics, including slope, aspect, roughness, topographic position index (TPI), and terrain ruggedness index (TRI). Aspect was converted from radians to northing and easting using cos() and sin() transformations respectively.

Soil data were downloaded from the UKCEH CountrySide Survey (https://catalogue.ceh.ac.uk/documents/467ada26-957a-40df-a60a-f385c828a923). Variables extracted were soil moisture, carbon concentration, carbon stock, bulk density, pH, carbon:nitrogen ratio, nitrogen concentration, phosphorus concentration, bacterial Shannon diversity index, invertebrate taxa richness and invertebrate Shannon diversity index. Data are representative of 256 0-15cm soil cores taken in 1998 (apart from soil bulk density and microbe Shannon index for which data were only available for 2007 when 591 soil cores were assessed).

Correlations between all 40 environmental variables were investigated using the R packages *virtualspecies* v.1.6^93^ and *corrplot* v.0.92 (https://github.com/taiyun/corrplot) (Supplementary Fig. 13). A custom R script was developed to iteratively remove one variable from each highly correlated pair until all remaining variables were only loosely correlated with each other (i.e. r =< 0.7). This resulted in a set of 14 variables (Supplementary Table 7), which were standardised to have a mean of 0 and standard deviation of 1.

#### LFMM

The lfmm2() and lfmm2.test() functions within the R package Landscape and Ecological Association Studies (*LEA*) v.3.10.2^27^ were used to perform genome-environment associations. Lfmm2() uses a frequentist method as opposed to lfmm() which is Bayesian ^28^. The former was chosen because it is less computationally intensive and has been shown to be less prone to false positives ^28^. A preliminary analysis of our data comparing both methods using 420,165 SNPs on chromosome 9 confirmed these earlier suggestions, as the amount of genomic inflation, and therefore the number of false positives, was much lower with lfmm2(), even when genomic control was implemented (Supplementary Fig.32 and Supplementary Fig.33).

Lfmm2() requires that all SNPs be genotyped and that a K value is provided to represent the number of latent factors (assumed number of ancestral populations/sources of hidden population structure). Missing genotypes were imputed using BEAGLE 5.4^94^. To determine an appropriate K value, a sparse non-negative matrix factorisation (SNMF) analysis was performed on a reduced set of 738,833 SNPs in approximate linkage equilibrium (see above) and without SNPs inside transposable elements, which were removed using bedtools^95^. SNMF was run at K=1 to K=30 for 7 repetitions (Supplementary Fig.16).

To complement the SNMF analysis, principal component analysis (PCA) of the genotype data was performed using LEA, and the Catell’s rule was used to identify the number of principal components explaining meaningful population structure (Supplementary Fig. 16). These analyses were run on two datasets. The first included all provenance trial individuals (n = 1873) excluding technical replicates. Eleven trees which were more closely related to provenances other than their own, and for which we could clearly identify a sampling error, were reassigned to their correct provenance. The second dataset only included individuals that showed 3^rd^ degree relatedness or less (see Methods, *SNP-heritability of phenotypic traits*). Individuals found to be more closely related to individuals from provenances other than their own, including the 11 mentioned above, were also removed (see Results, *Genomic diversity of samples*). This led to a reduced set of 1020 “unrelated” individuals. The latter dataset was created because 1) LFMMs do not explicitly control for close relatedness among samples, which can confound associations, and 2) to remove potentially mislabelled samples.

For the full dataset (n = 1,873), neither SNMF or PCA eigenvalues provided a clear indication of the ideal K value (Supplementary Figure 16a and 16b). On the other hand, analysis of the alternative dataset of 1020 ‘unrelated’ individuals showed K=11 as the clear minimum in the SNMF cross-entropy criterion (Supplementary Fig. 16c and 16d). We therefore fitted lfmm2() models with the 1,020 ‘unrelated’ individuals, the 14 environmental variables and all ∼5.9M SNPs. Nevertheless, we investigate how close kinship may affect GEA signals by comparing the full model with the “unrelated” model and present results in a Supplementary Figure 19 and Supplementary Text 1).

We also repeated the GEAs with the 1,020 ‘unrelated’ individuals but this time excluding soil variables, which are not often included in GEAs. We did so to assess how much unique genetic variation they explain compared to climatic and terrain variables (Supplementary Figure 17). All lfmm2() models were run with effect.size = TRUE, and p-values were generated using lfmm2.test() with linear = TRUE and genomic control = TRUE.

To investigate the contribution of each environmental variable to genetic variance, and find genetic correlation between variables, the matrix of genome-wide SNP effect sizes (β) was extracted from the lfmm2 output and transformed into a covariance matrix (Σ_β_) using a custom R script, which was used to derive the following metrics:

(1) Var_total_(j)=(Σ_β_)_jj_ is the diagonal element of the covariance matrix, or the total variance in effect sizes for environmental variable j across all loci.
(2) (Σ_β_)_reg_ = Σ_β_ + ridge × I_p_ is the ridge-regularized Σ_β_, where ridge = 10^-6^ and I_p_ is the p × p identity matrix.
(3) Var*_Con_*(j) = 1 / ((Σ_β_)_reg_ ¹)_jj_ = Var(β_j_ | β_k_ for k≠j) is the conditional R^2^, or the variance in effect sizes for environmental variable *j* that remains after accounting for correlations with all other environmental variables’ effects (k).
(4) Redundancy(j) = Var_total_(j) - Var*_Con_*(j) or the portion of variance in effect sizes for variable *j* that is explained by correlations with other environmental variables’ effects.
(5) Relative Unique(j) = Var*_Con_*(j) / Var_total_(j), or the portion of variance in effect sizes for variable j that is independent of other environmental variables.

The metrics above incorporate smaller, genome-wide effects and therefore do not rely on arbitrary SNP significant thresholds. This approach is more appropriate to polygenic adaptation, since in a medium powered design most of the important SNPs are expected to be non-significant. Metrics were calculated for all models i.e. with and without soil variables, and with and without related individuals.

To formally assess whether environmental variables explained variation in SNP effect sizes beyond random noise, we generated 10 dummy variables, each retaining provenance-level population structure such that all individuals within a provenance were assigned the same value. These were permuted alongside the 14 real environmental variables in lfmm2() to define the null expectation. The mean and standard errors (SE) of the total variation explained by the dummies across the 10 permutations were compared to the total variation explained by each environmental variable. Values exceeding 2 SE of the dummy mean were considered significant.

Manhattan plots (Extended Data Fig.5), and searches for candidate genes, were performed as above (Section *Manhattan plots and gene annotations, GWAS*). QQ-plots (Supplementary Fig. 20) were plotted using the R package GWASTools v1.44.0^96^. Allele frequencies of SNPs identified in the LFMMs were calculated per provenance using PLINK 2^64^ with the -freq flag (Supplementary Figure 23).

### Predicting and testing local adaptation

We trained models using genomic data and provenances’ source environmental conditions to predict how (mal)adapted they are to each of the three trial sites they were planted. Assuming growth-related phenotypes are reasonable fitness proxies^49^ allows us to test whether our predictions match the actual performance of the provenances at these sites. We used two different methods to predict local adaptation: allele shift and genomic gap.

#### AlleleShift

We used R package AlleleShift^30^ to train models using SNPs identified as associated with the environment in lfmm2(), which were then used to predict the optimum allele frequency of each SNP at each trial site. The AlleleShift pipeline is comprehensively described in Kindt^30^. Briefly, we first pruned closely linked statistically significant SNPs using SNPRelate v1.32.2 (10.1093/bioinformatics/bts606) with a correlation threshold of 0.4 within a sliding window of 50,000bp to reduce pseudo-replication and collinearity in the prediction models. This reduced the SNP set from 117 to 22. The R packages *vcfR*^97^ and *adegenet*^98^ were then used to generate allele counts per provenance for each of the 22 SNPs. An RDA model was built to characterise the relationship between provenance allele frequencies and the 14 uncorrelated, standardised environmental, terrain and soil variables. This model was further calibrated with a GAM model to eliminate occurrences of negative allele frequencies. The calibrated model was then used to predict the optimum allele frequencies expected under the environmental conditions at each trial site. The allele shift (often referred to as the residual or offset) was calculated as the absolute difference between the predicted allele frequency at the trial site condition and the actual, baseline allele frequency at each provenance’s local conditions. The mean absolute allele shift across all 22 SNPs was then used as a proxy of provenance mal-adaptedness, and was calculated for each provenance trial site combination.

We also repeated the pipeline on the 10,000 SNPs with the smallest p-values in the GEAs. After pruning for linkage disequilibrium, the number of SNPs decreased to ∼2.8K. This new analysis aims at capturing a more comprehensive genome-wide signal without focusing on statistical significance, which we believe is more appropriate to polygenic adaptation as stated above.

#### Genetic Gap

As an alternative approach to allele shifts we implemented the genetic.gap() function within the *LEA* R package. This approach provides a genome-wide estimate of (mal)adaptation using the effect sizes of all SNPs fitted in lfmm2()^99^. This genome-wide method has the advantage of explicitly dealing with correlations among environmental variables and SNPs with near-zero effects. This function also produces a ‘distance’ metric, representing a modified version of the Risk Of Non-Adaptedness (RONA), which includes correction for confounding factors and analyses multiple predictors simultaneously. Due to high computation requirements to implement genetic.gap(), we used the reduced dataset of 738,833 unlinked, outside of retrotransposons, SNPs as described above (Section, *LFMM)*. Genetic gaps were estimated per each provenance for each trial site.

#### Validating models

We tested theoretical expectations from AlleleShift and genetic.gap() (which will be jointly called “genetic offset” henceforth) by investigating whether provenances requiring larger genetic offsets to be adapted to a specific site would have lower growth performance at that site. We compared the predictive power of models using genetic offset estimates as predictors with models using latitudinal transfer or the environmental dissimilarity between provenance source location and trial sites as predictors. In other words, we ask the question “are genetic offset methods more accurate at predicting growth performance than simpler approaches relying solely on environmental or geographic information?”. We quantified differences in environmental conditions between each trial site and provenance combination using principal component analysis (PCA) on the 14 environmental variables used in the GEA. The first five principal components (PCs) explained over 71% of total environmental variation. Environmental dissimilarity was calculated as Euclidean and Mahalanobis distances between each provenance and trial site projected within the multi-dimensional PC space.

We repeated the estimation including only climatic variables (i.e. with terrain and soil variables removed), as climate is the aspect of the environment more commonly analysed in GEA studies.

Estimates of genetic offset and environmental dissimilarity, and latitudinal transfers were regressed on growth traits using linear mixed models in R. Models were fitted as described in the *Provenance performance* section, but since predicted slopes for the Drummond site differed drastically from the other two sites, we fitted models separately per site i.e. Trait∼ Predictor + Edge + Neighbours + (1|Block/Plot) instead of fitting a Predictor x Site interaction. P-values were estimated using LRT tests and adjusted using an FDR approach at alpha = 0.05 to correct for multiple testing caused by sites being tested separately. Model fits were compared using Akaike Information Criteria (AIC) adjusted for small sample sizes i.e. AICc.

Finally, we compared the different estimates of genetic offsets for each trial site with those predicted for each provenance under future environmental conditions, based on the worst-case climate change scenario (Representative Concentration Pathway (RCP) 8.5). This aimed at answering the question “Will provenances require greater genetic change to remain adapted to their source environments under future climate change than is currently required for adaptation at non-native sites?”.

Annual averages for each provenance were extracted for each 1 km-resolution CHELSA climatology for a period of 2071-2100. Data from the UKESM1-0-LL model was accessed from scenario SSP5-8.5 (RCP8.5). The MetOffice UK Climate Projections V.2.13.0 (UKPC) database was used to extract predicted wind speed at 10m for a time period of 2071-2081 at RCP8.5, or the equivalent worst case scenario assuming 4.0°C of warming by 2100 (https://ukclimateprojections-ui.metoffice.gov.uk/ui/home). The number of days with snow lying was not available on the UKPC database so could not be included in the future genetic offset estimations. Terrain variables were included as above (see *Environmental Variable Selection* section). No future predictions were available for the soil data, so these variables were excluded from this analysis.

Future, predicted variables were subset to those that were used in the original lfmm2() after removal of correlated variables, resulting in 10 variables. As before, variables were standardised and used in the LEA genetic.gap() command with 738,833 unlinked genome-wide SNPs, and in AlleleShift with sets of 22 unlinked and statistically significant SNPs and the top 2.8K unlinked GEA SNPs, for determination of the genetic maladaptation of each provenance to future conditions under a worst-case climate scenario at the provenance-source site.

Genetic offset estimates of each provenance to each trial site were re-estimated for just these 10 environmental variables using current climate data so that offsets were comparable between trial sites and RCP8.5 future scenarios. Genetic offset estimates were fitted in linear mixed effect models in the lme4 package with a categorical variable representing the trial sites and the RCP8.5 future scenario as the fixed effect, and provenance identity as a random effect i.e. GeneticOffset∼ Site + (1|Provenance). Post hoc pairwise comparisons between categories were then conducted using estimated marginal means (EMMs) in the emmeans v.1.10.0 package (<https://CRAN.R-project.org/package=emmeans).

## Extended Data

### Extended Figures

**Extended Figure 1:**
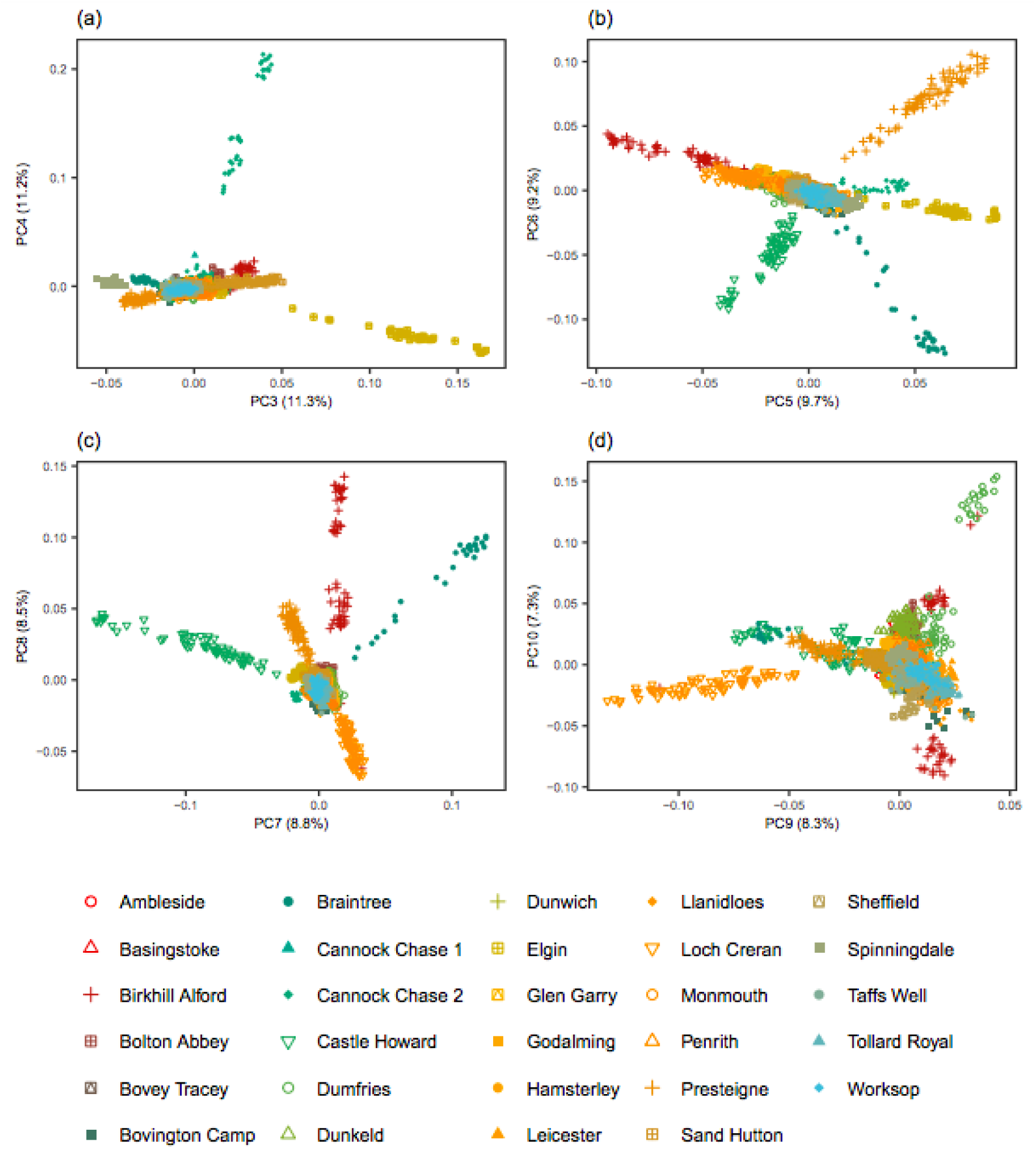
Genetic diversity among 29 silver birch (*Betula pendula*) provenances in the UK. Principal component analysis separated provenances with more related individuals. **a,** PC3 and PC4. **b**, PC5 and PC6. **c**, PC7 and PC8. **d**, PC9 and PC10. Individuals are colour and shape coded by provenance. Analysis was performed on 1.3 million LD-filtered SNPs from 1,873 provenance trial samples prior to removal of within- and between-provenance relatedness.

**Extended Figure 2:**
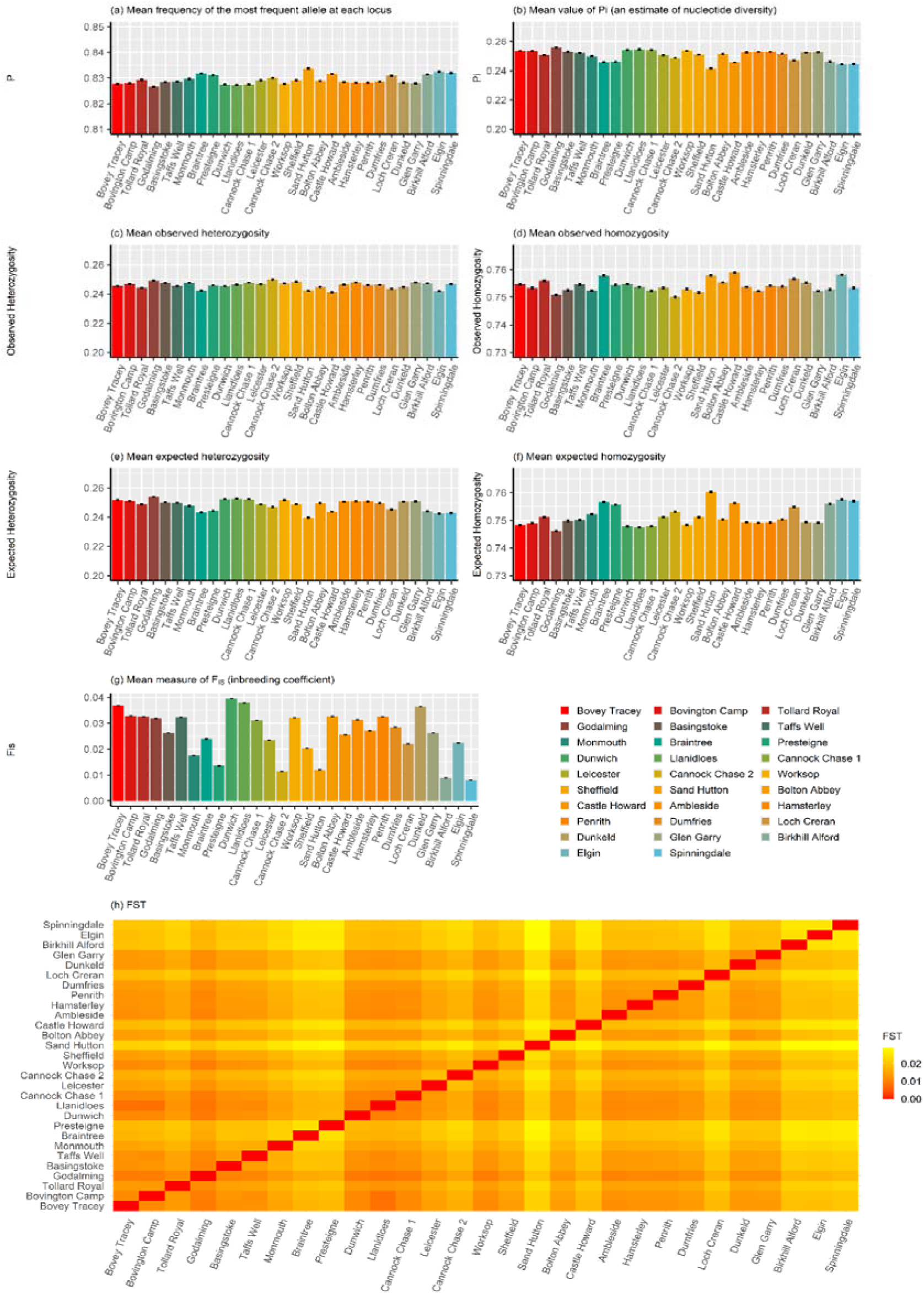
Population-level genetic diversity and between-population genetic differentiation in silver birch (*Betula pendula*). **a,** mean frequency of the most frequent allele at each locus (P). **b,** mean value of Pi (nucleotide diversity, π). **c,** mean observed heterozygosity (Obs_Het). **d,** mean observed homozygosity (Obs_Hom). **e,** mean expected heterozygosity (Exp_Het). **f,** mean expected homozygosity (Exp_Hom). **g,** inbreeding coefficient (FIS). **h,** heatmap of pairwise fixation index (F_ST_) values between each pair of provenances. Provenances are ordered by ascending latitude. Note the differing y-axis scales for each plot. Analysis was performed using STACKS populations software on the full dataset of 1,873 samples from 29 provenances, on a random, 25% subset of LD-filtered SNPs (136,185 SNPs). It should be noted that π was calculated on biallelic variable sites only, which produces higher values than genome-wide per-base-pair estimates. A similar figure is shown in Supplementary Fig. 8 for the 1020 “unrelated” trees.

**Extended Figure 3:**
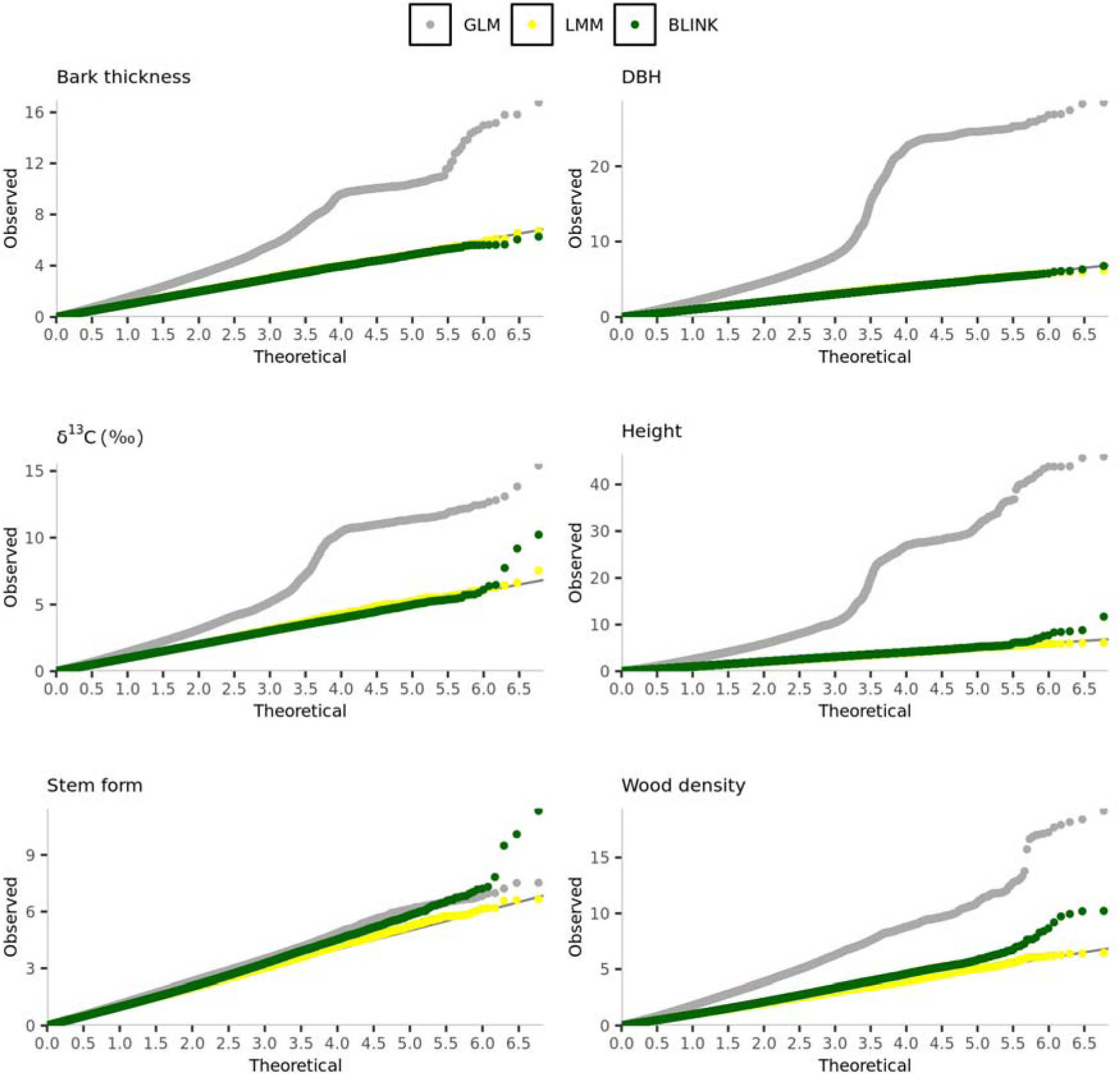
QQ-plots of GWAS results for multiple traits and association methods. Points are colour coded according to the GWAS method: Generalized Linear Model (GLM, dark grey), Linear Mixed Model (LMM, yellow), and Bayesian Least Absolute Shrinkage and Selection Operator (BLINK, green). GLM models show strong genomic inflation due to uncorrected relatedness within provenances.

**Extended Figure 4:**
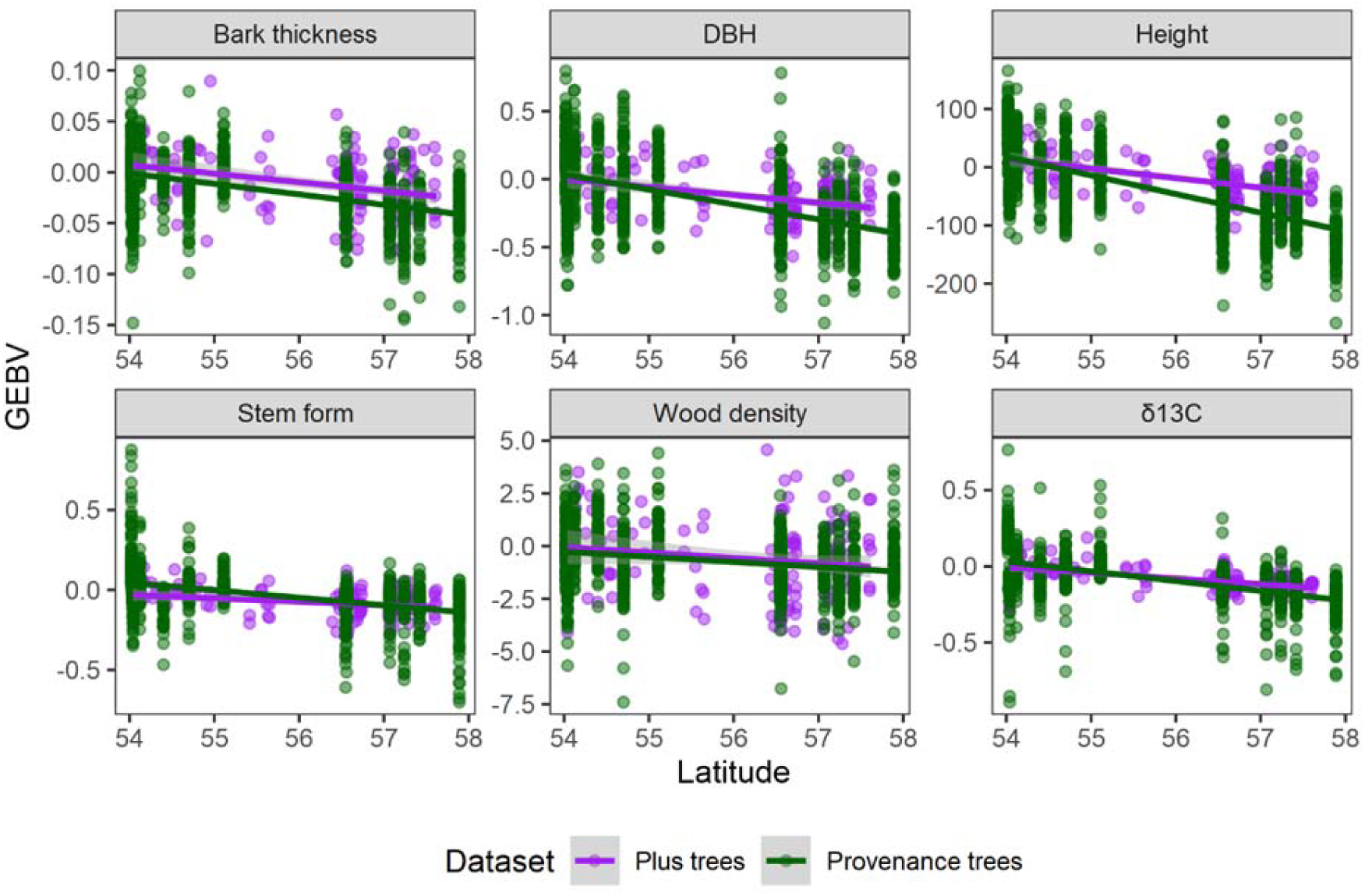
Relationship between latitude and genomic estimated breeding values (GEBVs) for individuals belonging to 14 northern provenances and plus trees from four collection locations. Plus trees are colour coded in purple and provenance trees in green. Lines represent the slope of a linear regression and shaded areas the confidence intervals.

**Extended Figure 5:**
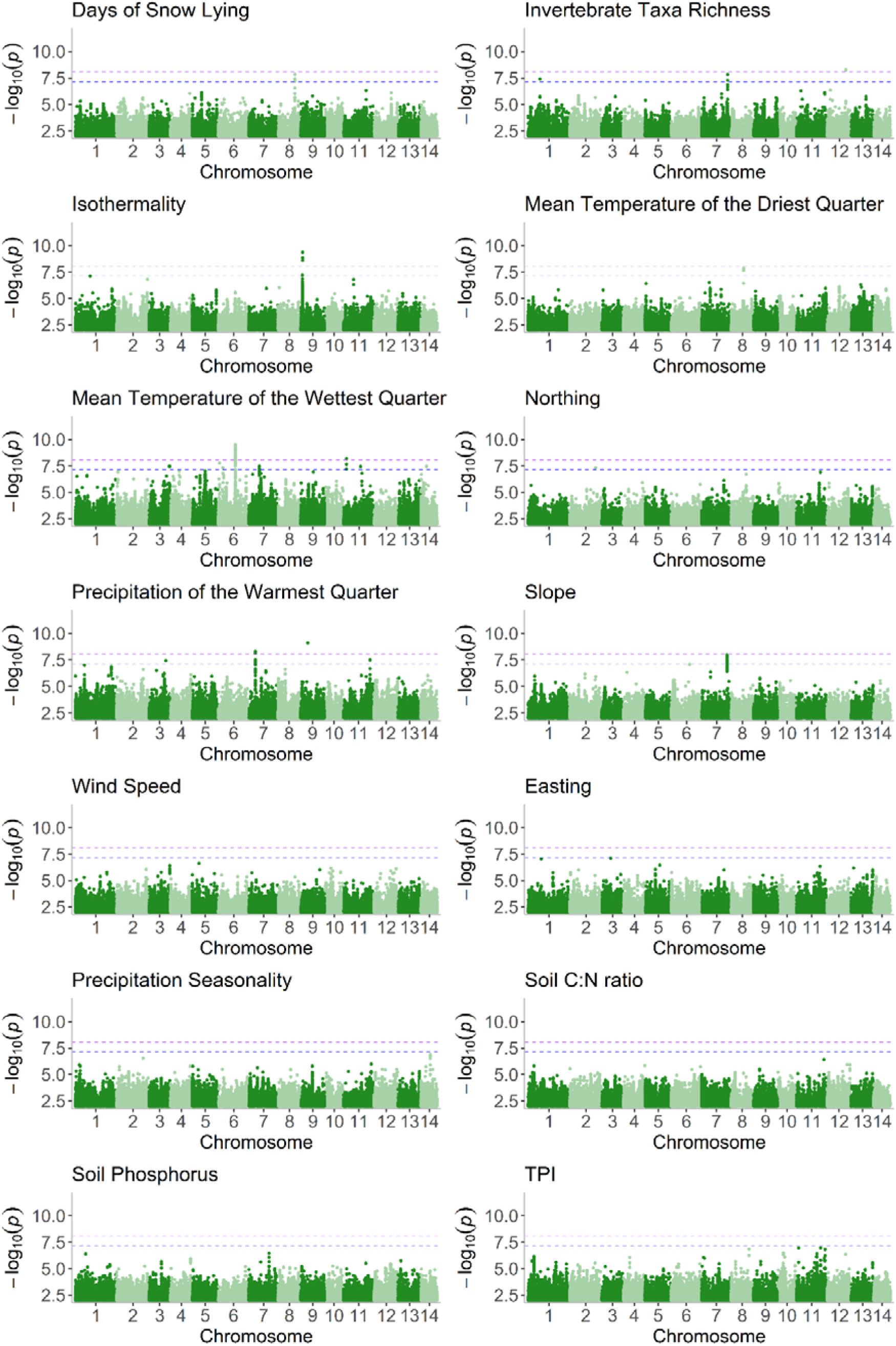
Genome-environment associations in silver birch (*Betula pendula*). Panels show Manhattan plots for each environmental variable (n = 14), with chromosome numbers on the x-axis and -log10 p-values on the y-axis. Each green dot represents a SNP, and coloured lines indicate significance thresholds (purple = Bonferroni, blue = modified Bonferroni). Associations were performed using LFMM2().

### Extended Tables

**Extended Table 1:** Summary statistics for silver birch (*Betula pendula*) phenotypic traits across three trial sites.

| Trait | Unit of measurement | Year of assessment | Trial site |  |  |
| --- | --- | --- | --- | --- | --- |
|  |  |  | Drummond | Llandovery | Thetford |
| DBH | cm | 2022 | 16.8 ± 5.32 SD | 12.1 ± 4.41 SD | 12.4 ± 3.34 SD |
| Height | cm | 2008 | 664 ± 126 SD | 804 ± 142 SD | 761 ± 110 SD |
| Bark thickness on exit | cm | 2023 | 0.912 ± 0.355 | 0.577 ± 0.277 SD | 0.685 ± 263 SD |
| Wood density | % amplitude | 2023 | 27.2 ± 3.59 SD | 28.3 ± 3.38 SD | 30.7 ± 3.34 SD |
| $\delta^{13}\text{C}$ | (ratio) | 2023 | -28.9 ± 1.3 SD | -27.5 ± 1.38 | -27.7 ± 0.963 SD |
| Crown condition | % | 2023 | 71.6 ± 14 SD | 67.8 ± 22.4 | 72.8 ± 13.7 SD |
| Stem form | count | 2022 | 1 = 19<br>2 = 148<br>3 = 403 | 1 = 108<br>2 = 375<br>3 = 135 | 1 = 1<br>2 = 555<br>3 = 114 |
| Mortality (by provenance) | rate | Up to 2023 | 0.279 ± 0.086 SD | 0.439 ± 0.097 SD | 0.0895 ± 0.048 SD |

**Extended Table 2:** Within-gene SNPs significantly associated with phenotypic traits or environmental variables in silver birch (*Betula pendula*). Note: GxE = genome-by-environment; MAF = minor allele frequency; I = inside the inversion. *For annotations supported by multiple SNPs, the statistics refers to the SNP with the largest effect size. The full list of statistically significant SNPs is provided in Supplementary Tables 4 and 5.

**Extended Table 1: Summary statistics for silver birch (*Betula pendula*) phenotypic traits across three trial sites.**
| Association | Chromosome: SNP* | $-\log_{10}$ p-value* | Effect size* | MAF* | SNPs (n) | Gene ID | Putative Gene Function | Model |
| --- | --- | --- | --- | --- | --- | --- | --- | --- |
| **GWAS** |  |  |  |  |  |  |  |  |
| Height | 1:1008256 | 11.67 | 42.41 | 0.1 | 1 | Bpev01.c0038.g0086 | Protein MAK16 homolog | BLINK |
| Wood density | 3:4402485 | 7.27 | 0.6 | 0.43 | 1 | Bpev01.c0540.g0029 | Glutathione-regulated potassium-efflux system protein | BLINK |
| Wood density | 9:17903555 | 7 | 0.52 | 0.39 | 1 | Bpev01.c1093.g0013 | Protein kinase family protein | BLINK |
| Wood density | 11:23347553 | 8.4 | 0.7 | 0.43 | 1 | Bpev01.c0644.g0008 | Phosphatidate cytidyltransferase | BLINK |
| Stem form | 2:7730618 | 10.08 | 0.11 | 0.35 | 1 | Bpev01.c0145.g0032 | Senescence-associated family protein | BLINK |
| Stem form | 5:7730618 | 9.49 | 0.1 | 0.39 | 1 | Bpev01.c0974.g0002 | RNA-binding protein 1 | BLINK |
| Stem form | 9:19818293 | 7.21 | 0.1 | 0.4 | 1 | Bpev01.c1670.g0005 | Ubiquitin carboxyl-terminal hydrolase 26 | BLINK |
| $\delta^{13}\text{C}$ | 8:7174043 | 10.22 | 0.26 | 0.29 | 1 | Bpev01.c0268.g0007 | TATA-binding protein-associated factor 172 | BLINK |
| $\delta^{13}\text{C}$ | 8:18361844 | 9.18 | -0.24 | 0.25 | 1 | Bpev01.c0029.g0044 | Cytochrome P450 | BLINK |
| Stem form (GxE) | 8:22179670 | 10.35 | -0.10 | 0.06 | 1 | Bpev01.c0478.g0018 | ABC transporter family G member | GCTA |
| Height | 6:8896694<sup>1</sup> | 10.89 | 4.66 | 0.38 | 43 | Bpev01.c0131.g0003 | Accelerated cell death | GCTA |
| Stem form | 13:16043962 | 8.31 | -0.09 | 0.06 | 3 | Bpev01.c0448.g0002 | AP2 family protein | GCTA |
| Stem form (GxE) | 4:3483942 | 7.88 | -0.10 | 0.05 | 1 | Bpev01.c0947.g0002 | Basic helix loop helix (bHLH) DNA-binding superfamily protein | GCTA |
| Height | 6:9179004<sup>1</sup> | 8.68 | 5.8 | 0.42 | 10 | Bpev01.c2410.g0002 | Cationic amino acid transporter 5 | GCTA |
| Height | 6:8932571<sup>1</sup> | 9.97 | 5.83 | 0.4 | 98 | Bpev01.c0131.g0004 | Cleavage and polyadenylation specificity factor 30 | GCTA |
| DBH | 6:8934556<sup>I</sup> | 7.39 | 0.04 | 0.38 | 1 | Bpev01.c0131.g0004 | Cleavage and polyadenylation specificity factor 30 | GCTA |
| Height | 6:8979077<sup>I</sup> | 9.3 | 6.3 | 0.38 | 244 | Bpev01.c0131.g0005 | SIT4 phosphatase-associated family protein | GCTA |
| DBH | 6:8968658<sup>I</sup> | 7.54 | 0.04 | 0.38 | 2 | Bpev01.c0131.g0005 | SIT4 phosphatase-associated family protein | GCTA |
| Height | 8:3074515 | 7.71 | 8.43 | 0.17 | 1 | Bpev01.c1836.g0006 | Cytochrome P450 | GCTA |
| Height | 10:12671968 | 7.25 | 9.77 | 0.2 | 1 | Bpev01.c0713.g0012 | DDT domain-containing protein | GCTA |
| Height | 11:29295864 | 7.39 | -11.5 | 0.06 | 3 | Bpev01.c0027.g0036 | Nudix hydrolyze | GCTA |
| Height | 6:8855335<sup>I</sup> | 9.22 | 7.25 | 0.29 | 123 | Bpev01.c0131.g0002 | Outer arm dynein light chain 1 protein | GCTA |
| DBH | 6:8845038<sup>I</sup> | 7.44 | 0.05 | 0.36 | 1 | Bpev01.c0131.g0002 | Outer arm dynein light chain 1 protein | GCTA |
| Height | 12:22824582 | 7.85 | -0.07 | 0.15 | 1 | Bpev01.c0349.g0020 | Pectate lyase family protein | GCTA |
| Height | 4:18240832 | 7.66 | -0.05 | 0.07 | 1 | Bpev01.c0863.g0014 | Pentatricopeptide repeat-containing protein | GCTA |
| Height | 6:9119612<sup>I</sup> | 7.94 | 5.79 | 0.4 | 4 | Bpev01.c0131.g0008 | Protein of unknown function (DUF1644) | GCTA |
| Stem form (GxE) | 11:20415284 | 7.27 | 0.03 | 0.06 | 1 | Bpev01.c0391.g0005 | Transmembrane_amino_acid_transporter_family_protein | GCTA |
| Height | 2:9618536 | 7.46 | -10.7 | 0.07 | 1 | Bpev01.c0080.g0038 | Unknown protein | GCTA |
| Height | 6:16566364 | 7.25 | -7.88 | 0.28 | 1 | Bpev01.c1156.g0001 | Unknown protein | GCTA |
| DBH | 14:12049034 | 8.15 | -0.09 | 0.06 | 1 | Bpev01.c1402.g0013 | Heteroglycan glucosidase 1 | GCTA |
| Bark thickness | 14:12049034 | 8.84 | -0.03 | 0.06 | 1 | Bpev01.c1402.g0013 | Heteroglycan glucosidase 1 | GCTA |
| Height | 11:29326884 | 7.8 | -9.7 | 0.07 | 1 | Bpev01.c0027.g0032 | sequence-specific_DNA_binding_transcription_factors | GCTA |
| Height | 6:9090832<sup>I</sup> | 8.08 | 6.62 | 0.39 | 41 | Bpev01.c0131.g0007 | Transducin family protein/WD-40 repeat family protein | GCTA |
| **GEA** |  |  |  |  |  |  |  |  |
| Mean temperature of the wettest quarter | 3:22099726 | 7.47 | 9.571 | 0.05 | 1 | Bpev01.c0473.g0010 | Pentatricopeptide repeat-containing protein | lfmm2 |
| Mean temperature of the wettest quarter | 7:11378787 | 7.46 | 9.65 | 0.051 | 2 | Bpev01.c0118.g0001 | Protein of unknown function (DUF789) | lfmm2 |
| Isothermality | 9:1996195 | 7.20 | 8.94 | 0.19 | 1 | Bpev01.c0214.g0030 | Unknown protein | lfmm2 |
| Isothermality | 9:2256742 | 8.85 | 9.995 | 0.069 | 1 | Bpev01.c0214.g0058 | Unknown protein | lfmm2 |
| Precipitation of the warmest quarter | 9:8209694 | 9.11 | 9.803 | 0.054 | 1 | Bpev01.c0025.g0018 | Plant protein of unknown function (DUF828) with plant pleckstrin homology-like region | lfmm2 |
| Precipitation of the warmest quarter | 11:29280149 | 7.49 | 8.8 | 0.083 | 2 | Bpev01.c0027.g0037 | Pentatricopeptide repeat-containing protein | lfmm2 |
| Northing | 2:28600944 | 7.31 | 8.52 | 0.07 | 1 | Bpev01.c2063.g0004 | External alternative NAD(P)H-ubiquinone oxidoreductase B1 | lfmm2 |

**Extended Table 3:** Comparison between genomic estimated breeding values (GEBVs) of provenance and plus trees under a latitudinal gradient. Lat = latitude. Significant p-values (<0.05) are highlighted in bold. Each model was based on 1000 observations.

| Model | Fixed effects | Estimates | CI | F | p |
| --- | --- | --- | --- | --- | --- |
| **Bark thickness**<br>$R^2 / R^2 \text{ adjusted} = 0.201 / 0.199$ | (Intercept) | 0.46 | 0.21 – 0.72 | - | - |
|  | Latitude | -0.01 | -0.01 – -0.00 | - | - |
|  | Dataset [Provenance trees] | 0.09 | -0.18 – 0.35 | - | - |
|  | Lat × Dataset [Provenance trees] | <0.01 | -0.01 – 0.00 | 0.57 | 0.452 |
| **DBH**<br>$R^2 / R^2 \text{ adjusted} = 0.283 / 0.280$ | (Intercept) | 3.13 | 0.98 – 5.29 | - | - |
|  | Latitude | -0.06 | -0.10 – -0.02 | - | - |
|  | Dataset [Provenance trees] | 2.86 | 0.62 – 5.10 | - | - |
|  | Lat × Dataset [Provenance trees] | -0.05 | -0.09 – -0.01 | 6.66 | **0.009** |
| **$\delta^{13}\text{C}$**<br>$R^2 / R^2 \text{ adjusted} = 0.278 / 0.275$ | (Intercept) | 2.01 | 0.74 – 3.28 | - | - |
|  | Latitude | -0.04 | -0.06 – -0.01 | - | - |
|  | Dataset [Provenance trees] | 1.5 | 0.17 – 2.82 | - | - |
|  | Lat × Dataset [Provenance trees] | -0.03 | -0.05 – -0.00 | 5.03 | **0.025** |
| **Stem form**<br>$R^2 / R^2 \text{ adjusted} = 0.185 / 0.182$ | (Intercept) | 1.13 | -0.12 – 2.38 | - | - |
|  | Latitude | -0.02 | -0.04 – 0.00 | - | - |
|  | Dataset [Provenance trees] | 1.46 | 0.16 – 2.76 | - | - |
|  | Lat × Dataset [Provenance trees] | -0.03 | -0.05 – -0.00 | 4.73 | **0.029** |
| **Height**<br>$R^2 / R^2 \text{ adjusted} = 0.432 / 0.430$ | (Intercept) | 897.31 | 447.80–1346.83 | - | - |
|  | Latitude | -16.36 | -24.33 – -8.39 | - | - |
|  | Dataset [Provenance trees] | 851.73 | 384.10–1319.36 | - | - |
|  | Lat × Dataset [Provenance trees] | -15.7 | -24.00 – -7.40 | 13.61 | **<0.001** |
| **Wood density**<br>$R^2 / R^2 \text{ adjusted} = 0.047 / 0.044$ | (Intercept) | 13.87 | 0.27 – 27.46 | - | - |
|  | Latitude | -0.26 | -0.50 – -0.02 | - | - |
|  | Dataset [Provenance trees] | -1.47 | -15.61 – 12.67 | - | - |
|  | Lat × Dataset [Provenance trees] | 0.02 | -0.23 – 0.27 | 0.02 | 0.887 |

**Extended Table 4:**
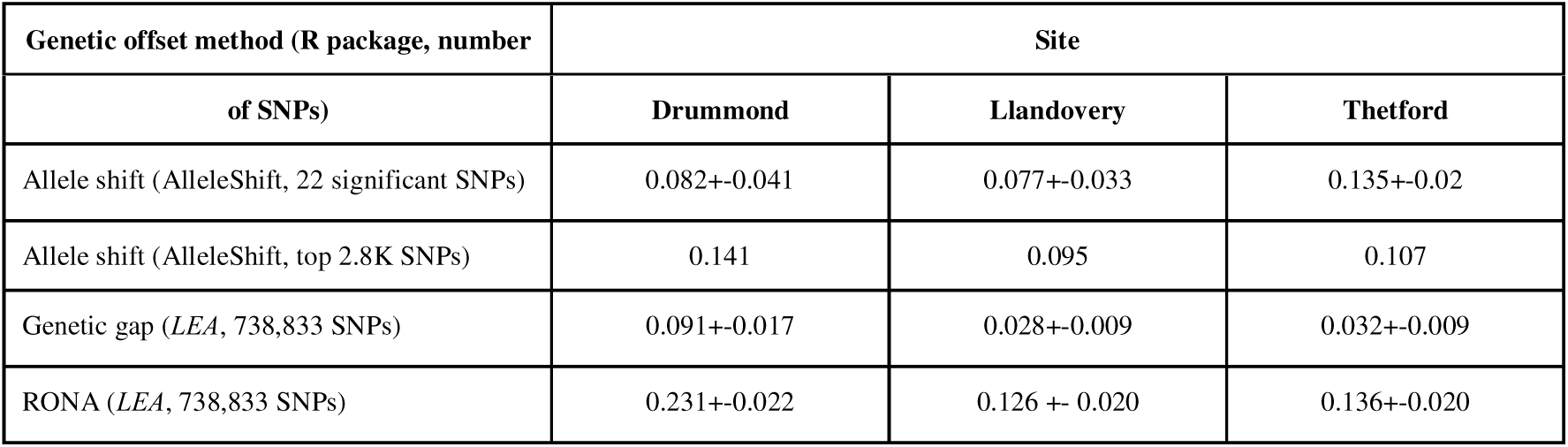
Genetic offset estimates across 29 silver birch provenance seed source locations for genetic suitability at each trial site location. Offsets were estimated between provenance and trial site local environmental conditions using the *AlleleShift* package on 22 SNPs significantly associated with environmental variables and the top 2.8K SNPs ordered by -log(p). The *LEA* package was used to estimate genetic gaps and risk of non-adaptedness (RONA) using 738,833 SNPs. SNPs used in the analyses were pruned to approximate linkage equilibrium from an original set of 117 (Allele Shift, 22 significant SNPs), 10k (Allele shift, top ∼2.8K SNPs) and ∼5.9M (*LEA*) SNPs, respectively.

## Supplementary information

### Supplementary Figures

**Supplementary Figure 1:**
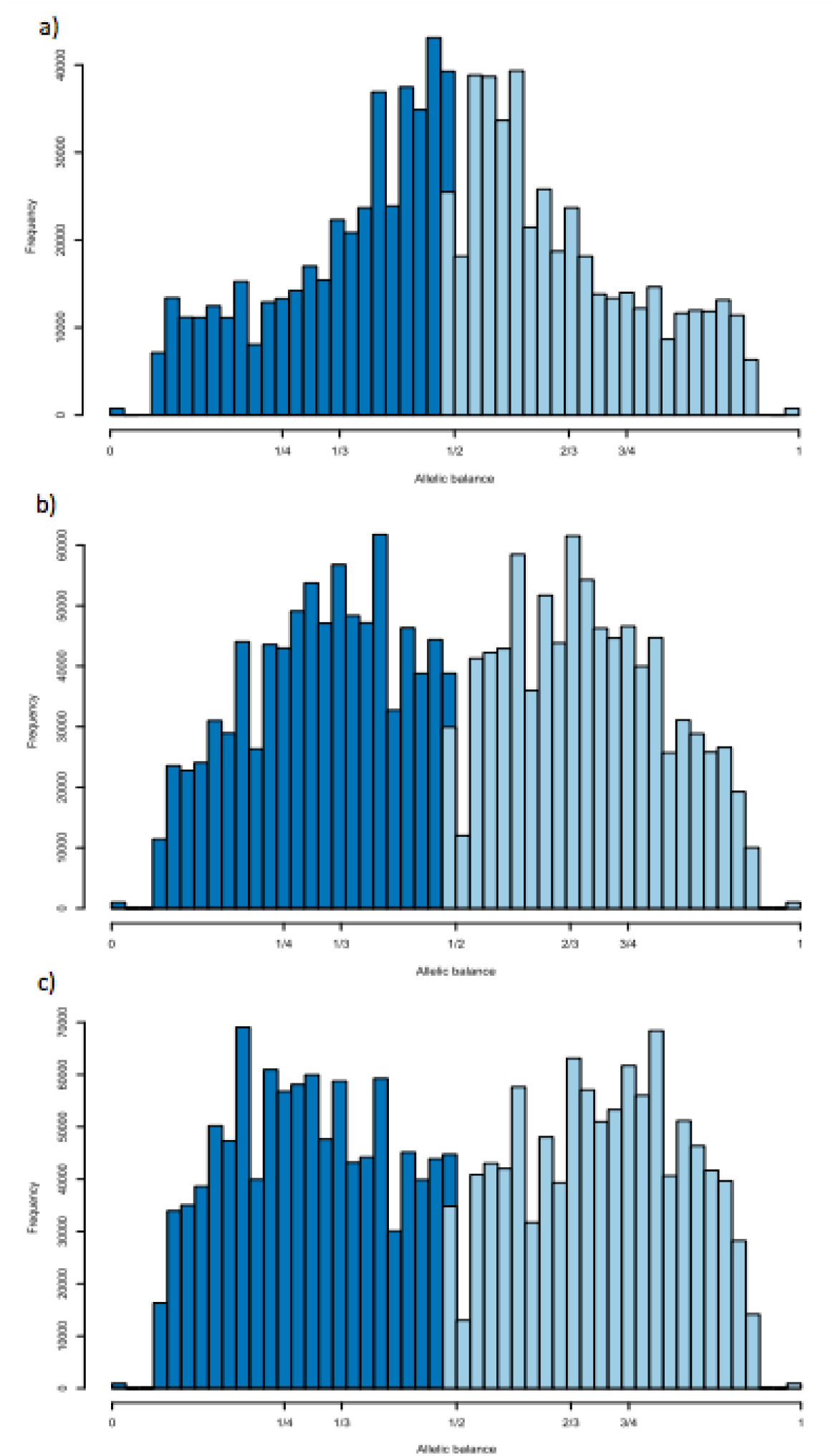
Histograms of allelic balance ratios at heterozygous sites. Alleles A and B are represented by dark and light blue bars, respectively. **a,** Diploid trees exhibit frequency peaks around 1/2 (ie. 0.5). **b,** triploids trees have peaks at 1/3 and 2/3 (ie. 0.33 or 0.66). **c,** tetraploids at 1/4 and 3/4 (ie. 0.25 and 0.75).

**Supplementary Figure 2:**
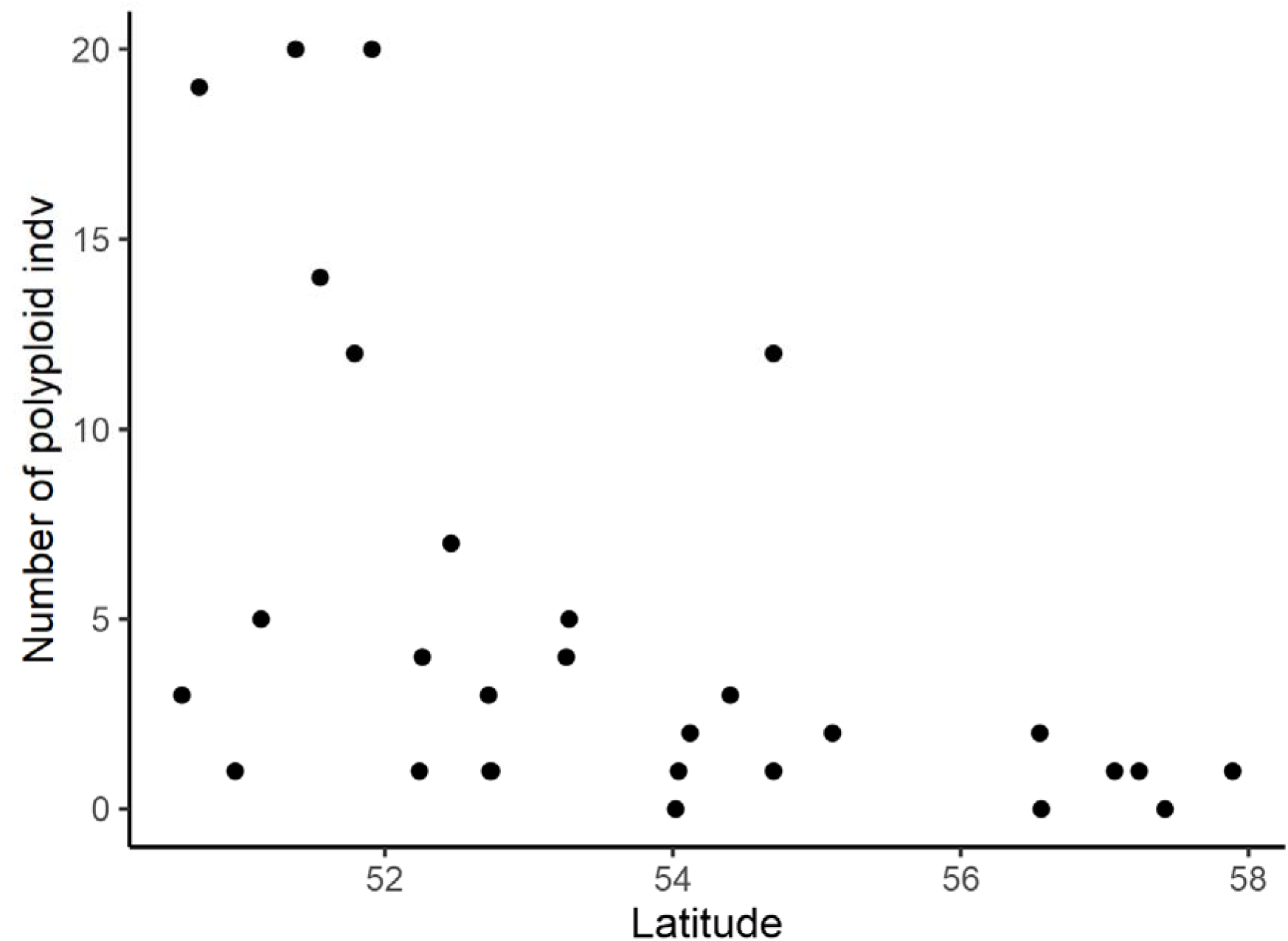
Number of polyploid individuals (i.e. triploids and tetraploids) by the latitude of its provenance of origin. Polyploids occurred more often at lower latitudes.

**Supplementary Figure 3:**
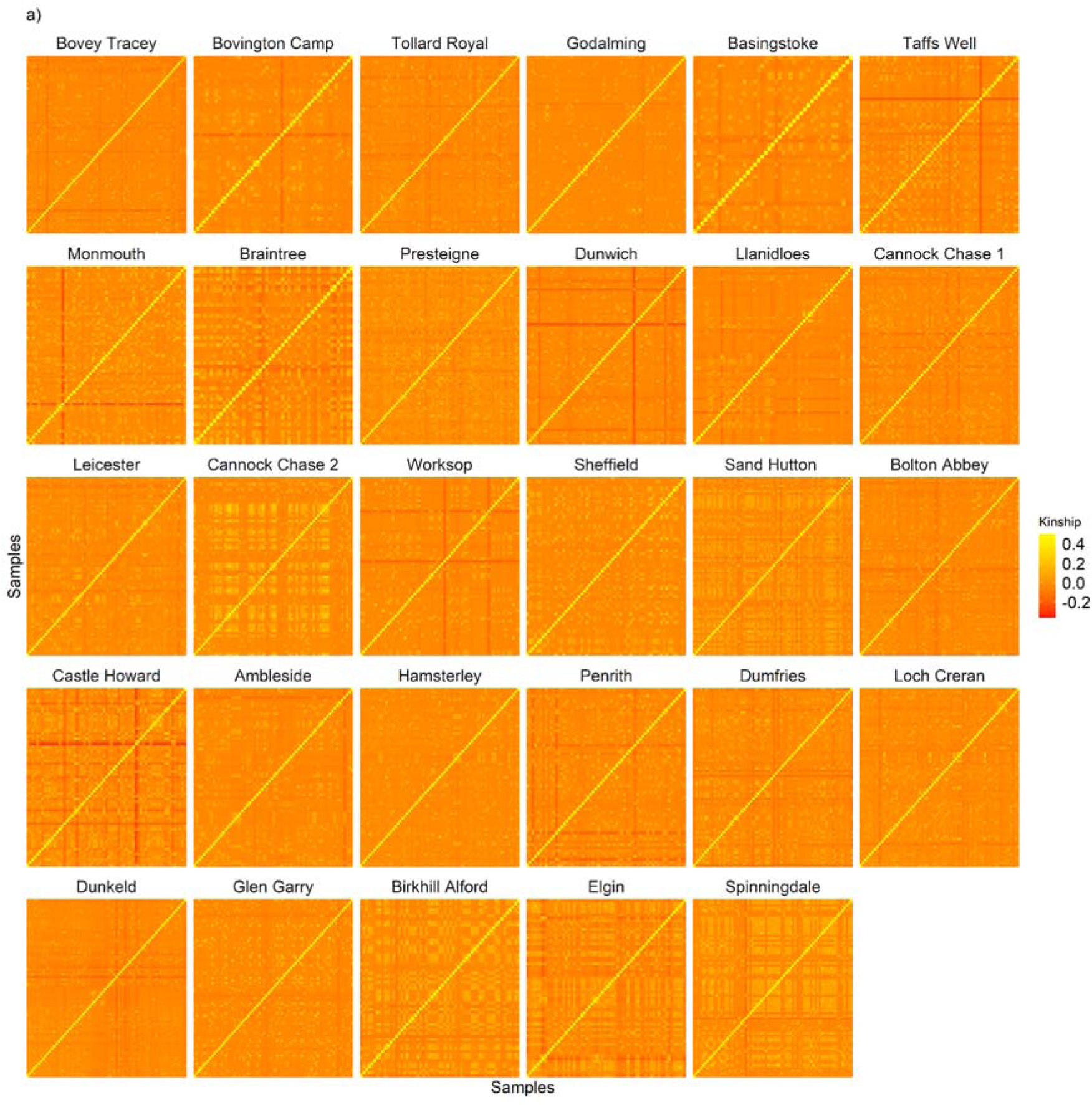
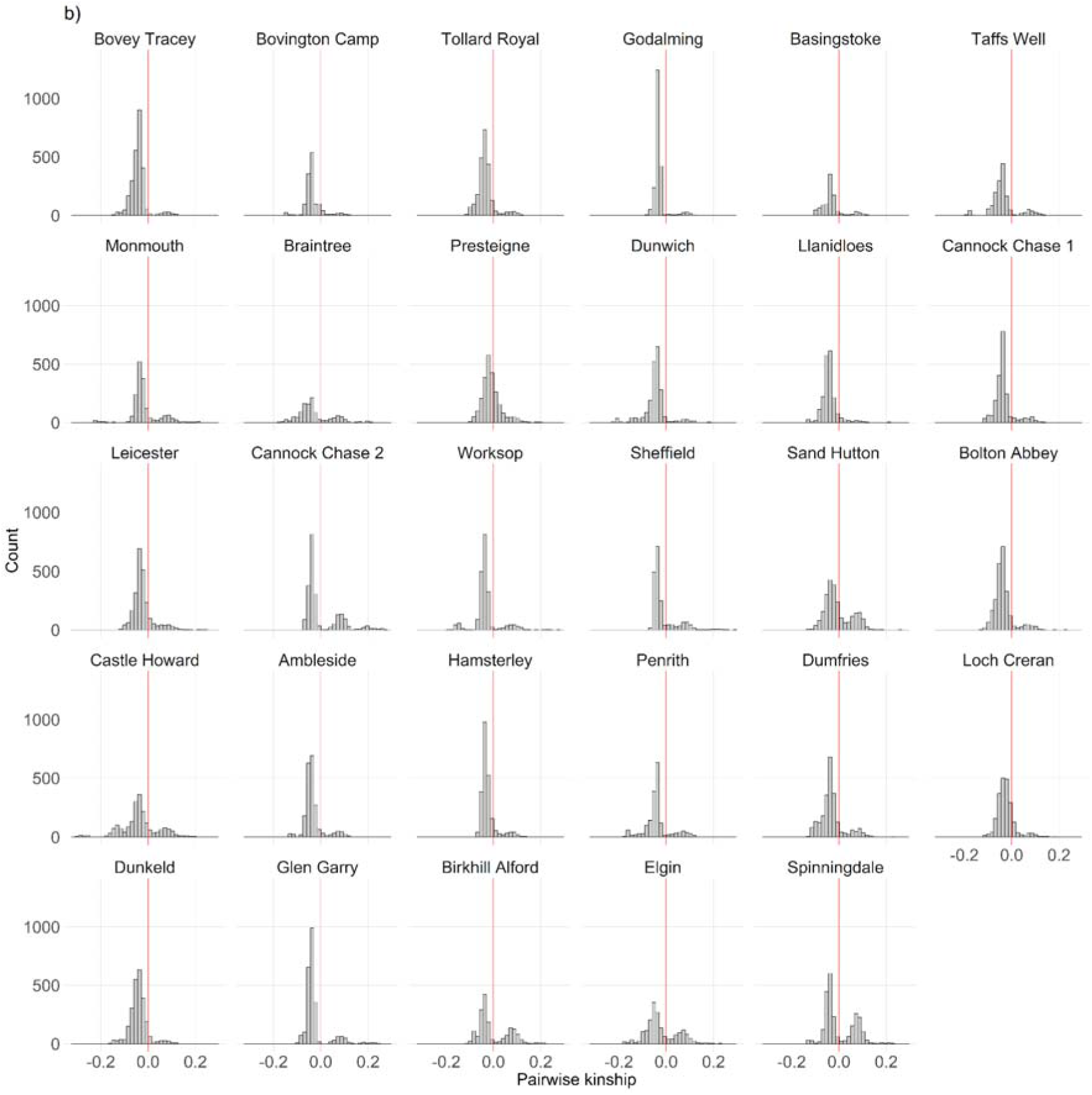
Intra-provenance pairwise kinship for each silver birch (*Betula pendula*) provenance. **a,** heatmaps of kinship with More yellowish colours indicate higher pairwise relatedness, whereas more reddish colours indicate the opposite. **b,** histograms of kinship. Provenances are ordered by latitude with the most southerly first (Bovey Tracey), and the most northerly last (Spinningdale). Kinship analysis was performed on 1,873 provenance trial samples from 29 provenances. SNPs in LD were removed prior to analysis.

**Supplementary Figure 4:**
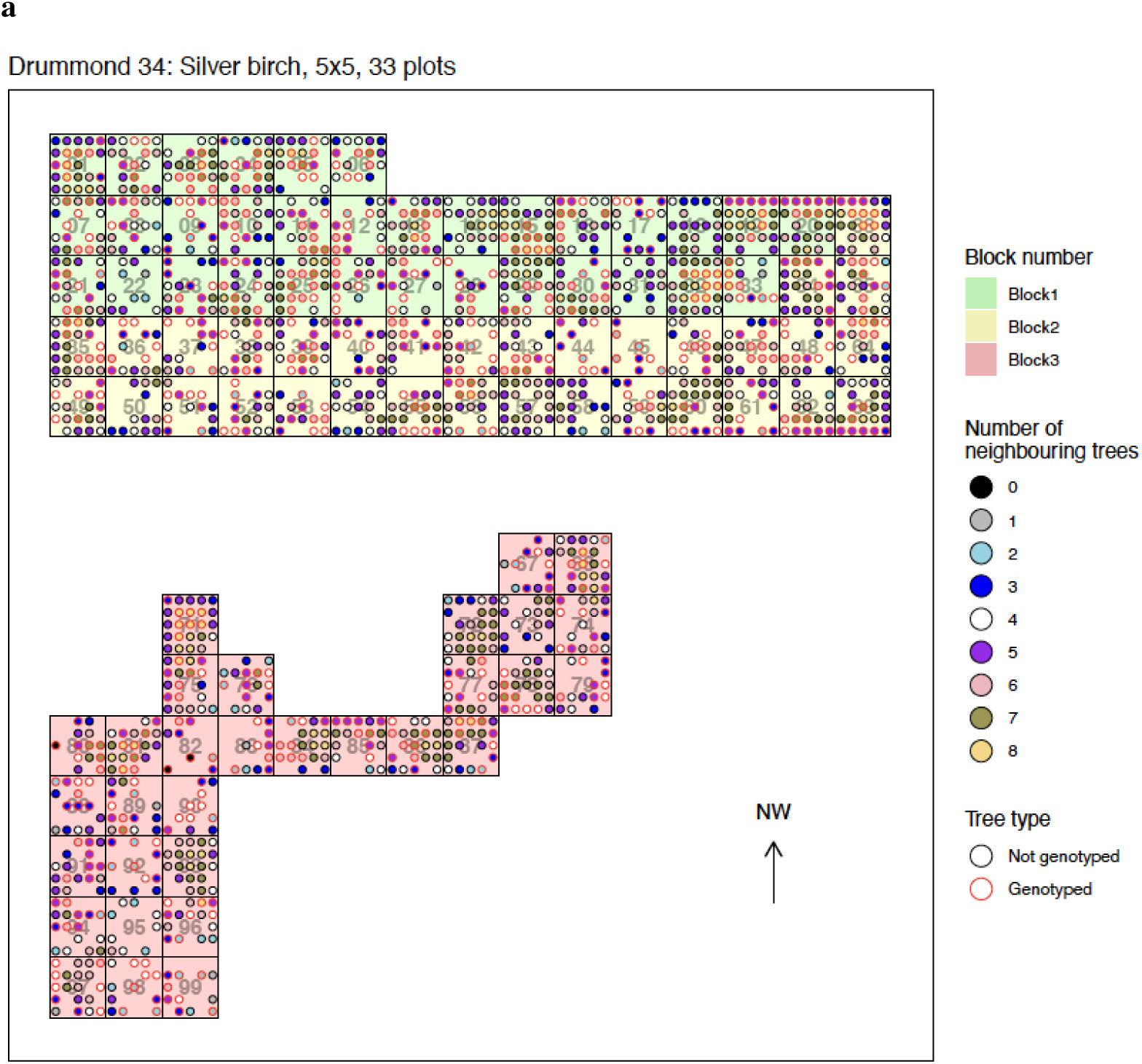
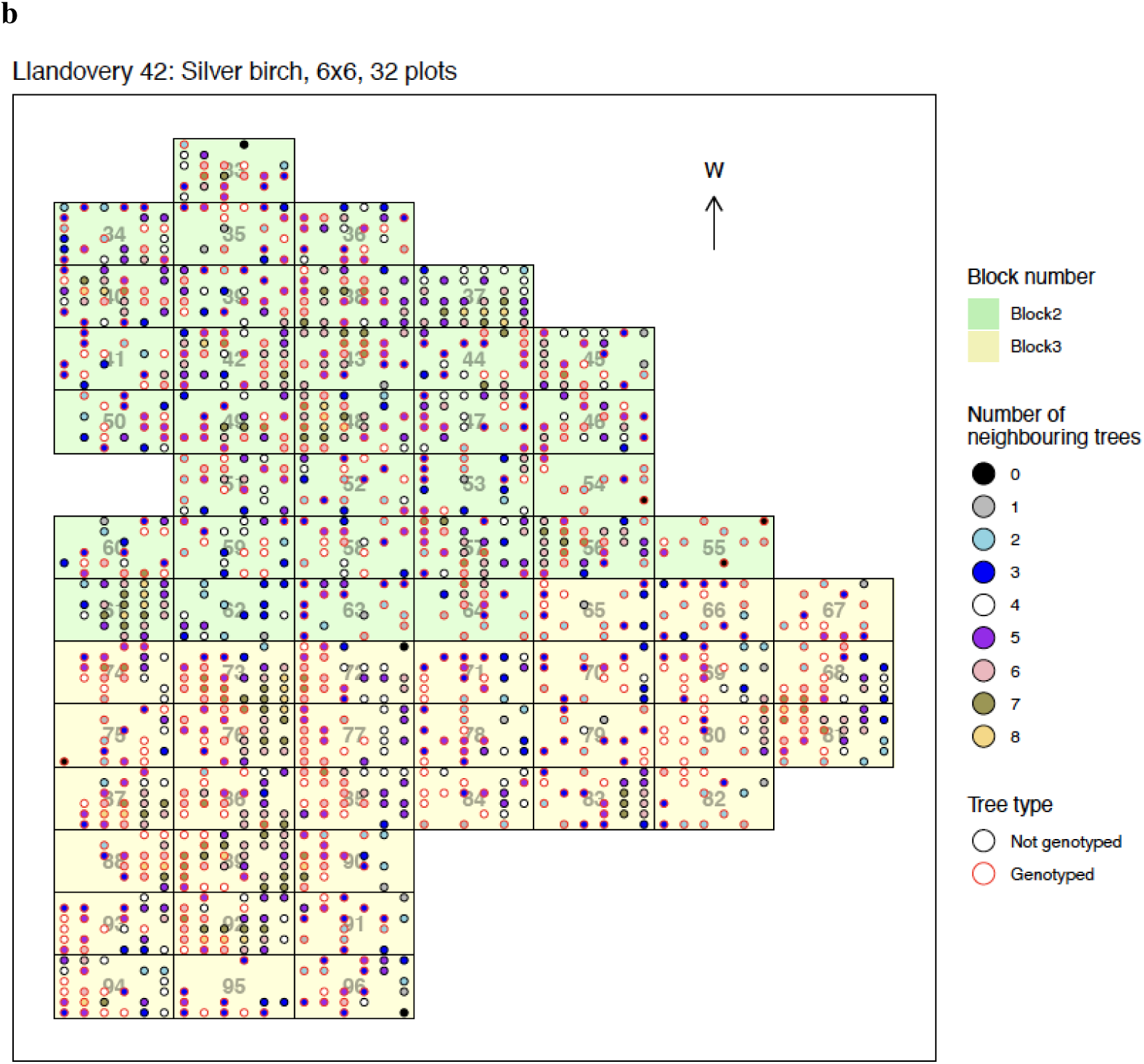
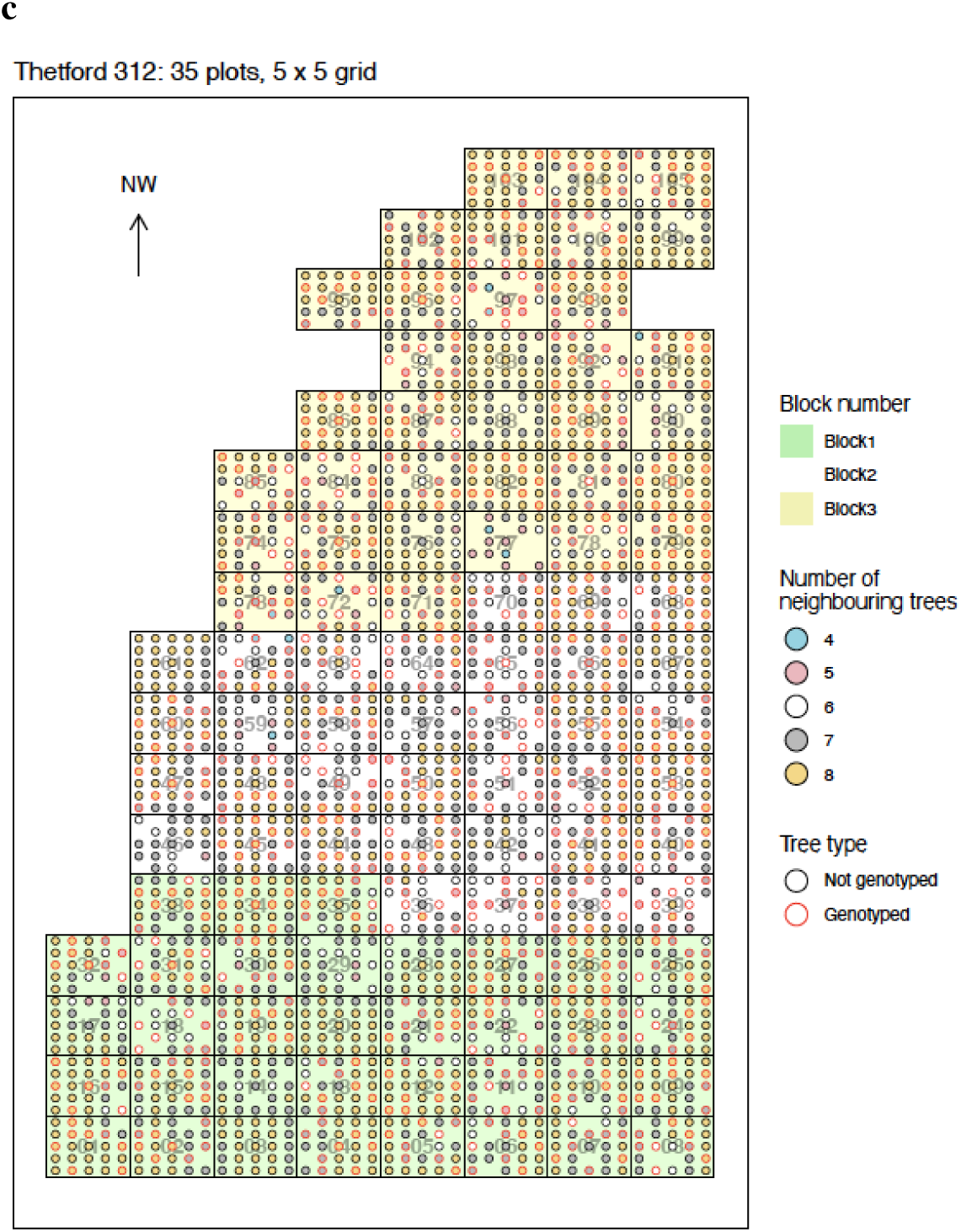
Aerial schematics of silver birch (*Betula pendula*) provenance trials showing layout, number of neighbours. Individual trees are denoted by circular points coloured according to their number of neighbours, with gaps indicating dead trees. Genotyped and phenotyped trees have red outlines. Figures show trial sites **a,** Drummond, **b,** Llandovery, and **c,** Thetford.

**Supplementary Figure 5:**
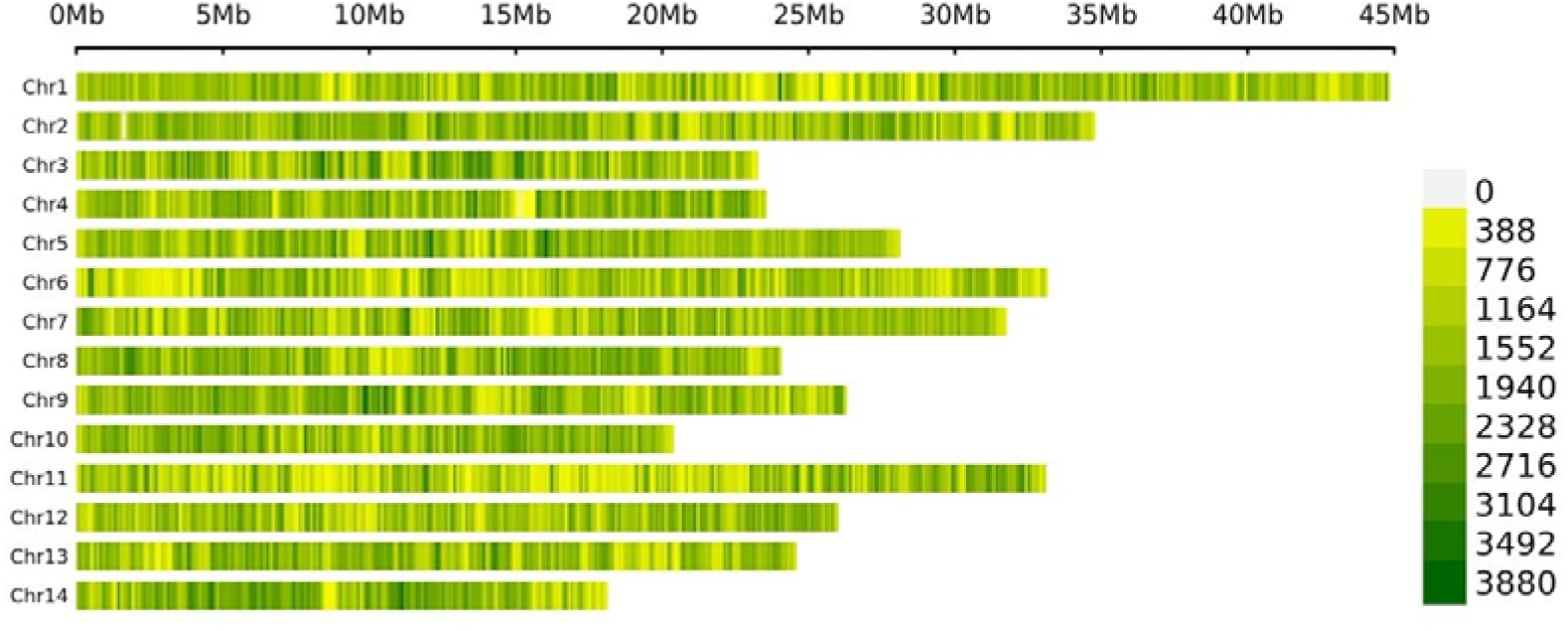
SNP-density plot by chromosome. Higher density regions are coloured in green, and lower density regions in yellow. Analysis was performed using the R package *CMplot* and included ∼5.9 million SNPs from 1,873 trees across 29 provenances.

**Supplementary Figure 6:**
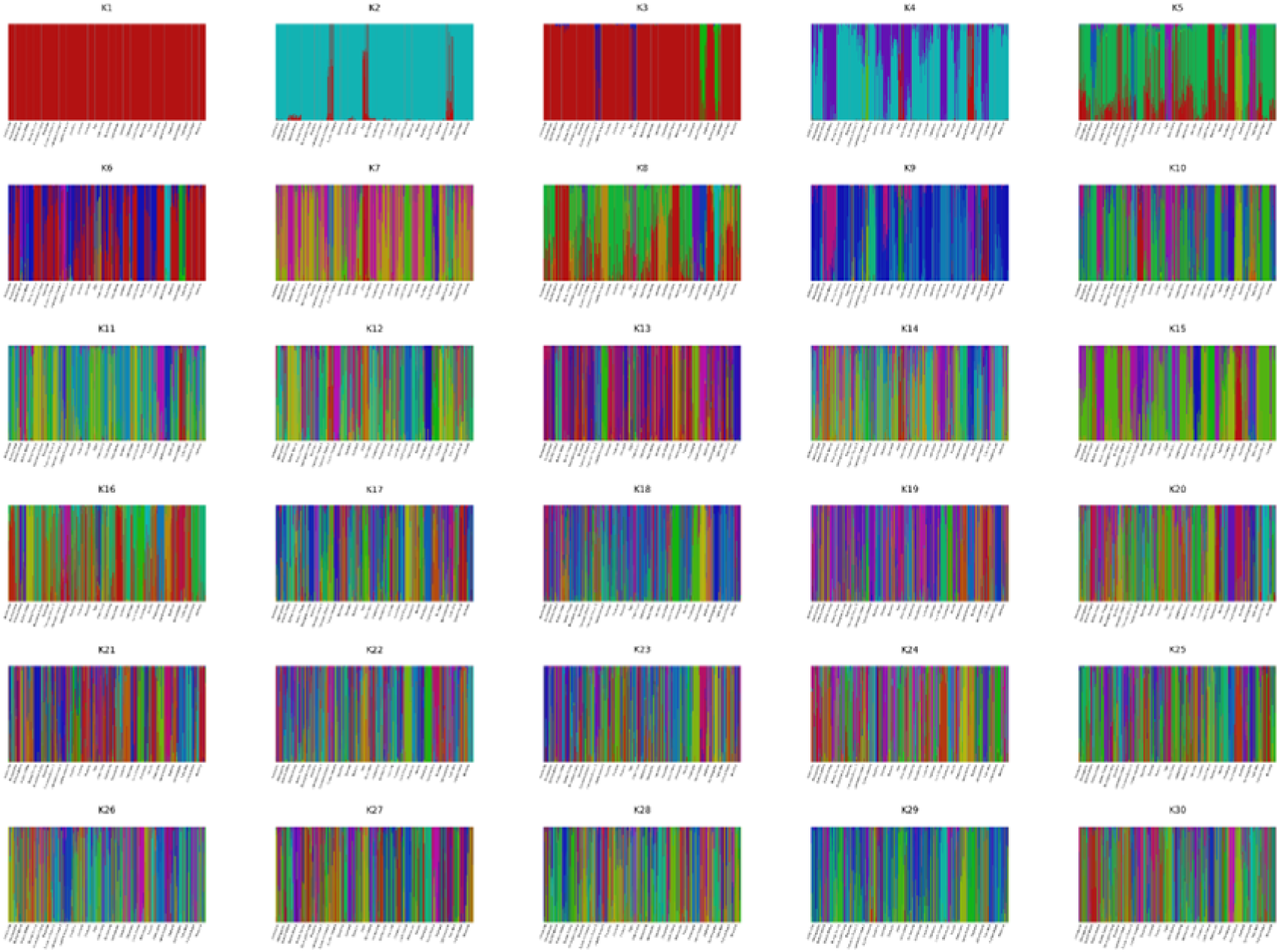
Admixture proportions estimated for different values of K in 1,873 silver birch (*Betula pendula*) individuals across 29 provenances. Analysis was performed using fastSTRUCTURE from K1-K30 with 5 cross-validation tests. SNPs in LD were filtered prior to analysis leaving 1.3 million variants. The chooseK.py script within fastSTRUCTURE identified K=10 as the simplest model (maximum marginal likelihood), and K=29 as optimal for explaining the structure in the dataset.

**Supplementary Figure 7:**
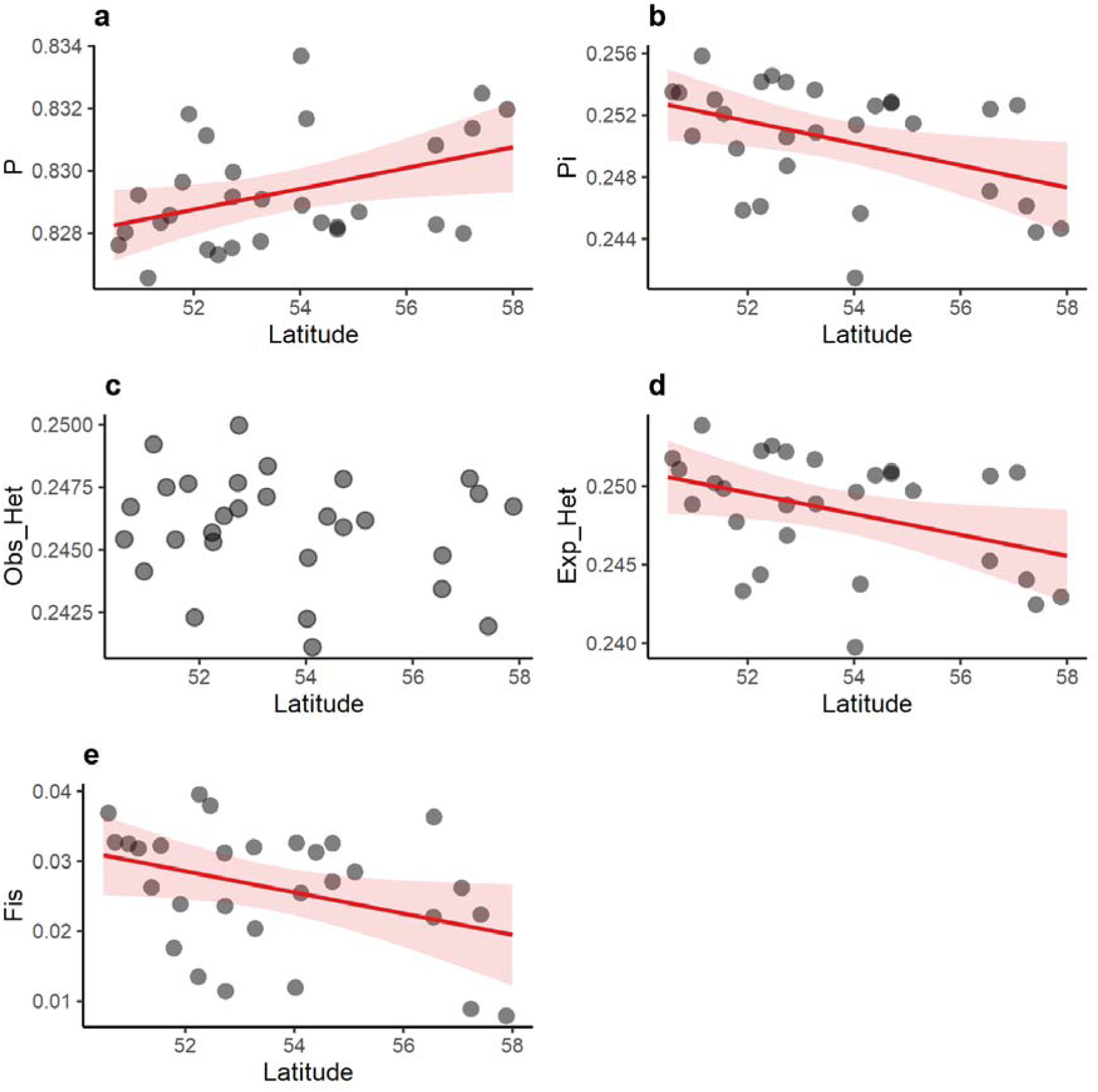
Relationship between latitude and genetic diversity metrics in silver birch (Betula pendula) provenances. **a,** mean frequency of the major alleles. **b,** nucleotide diversity. **c,** observed heterozygosity. **d,** expected heterozygosity. **e,** inbreeding coefficient. Red lines indicate the regression slope and red shades the confidence intervals. Metrics were calculated using STACKS on a 25% random subset of LD-filtered variable sites (341,081 SNPs). It should be noted that π was calculated on biallelic variable sites only, which produces higher values than genome-wide per-base-pair estimates.

**Supplementary Figure 8:**
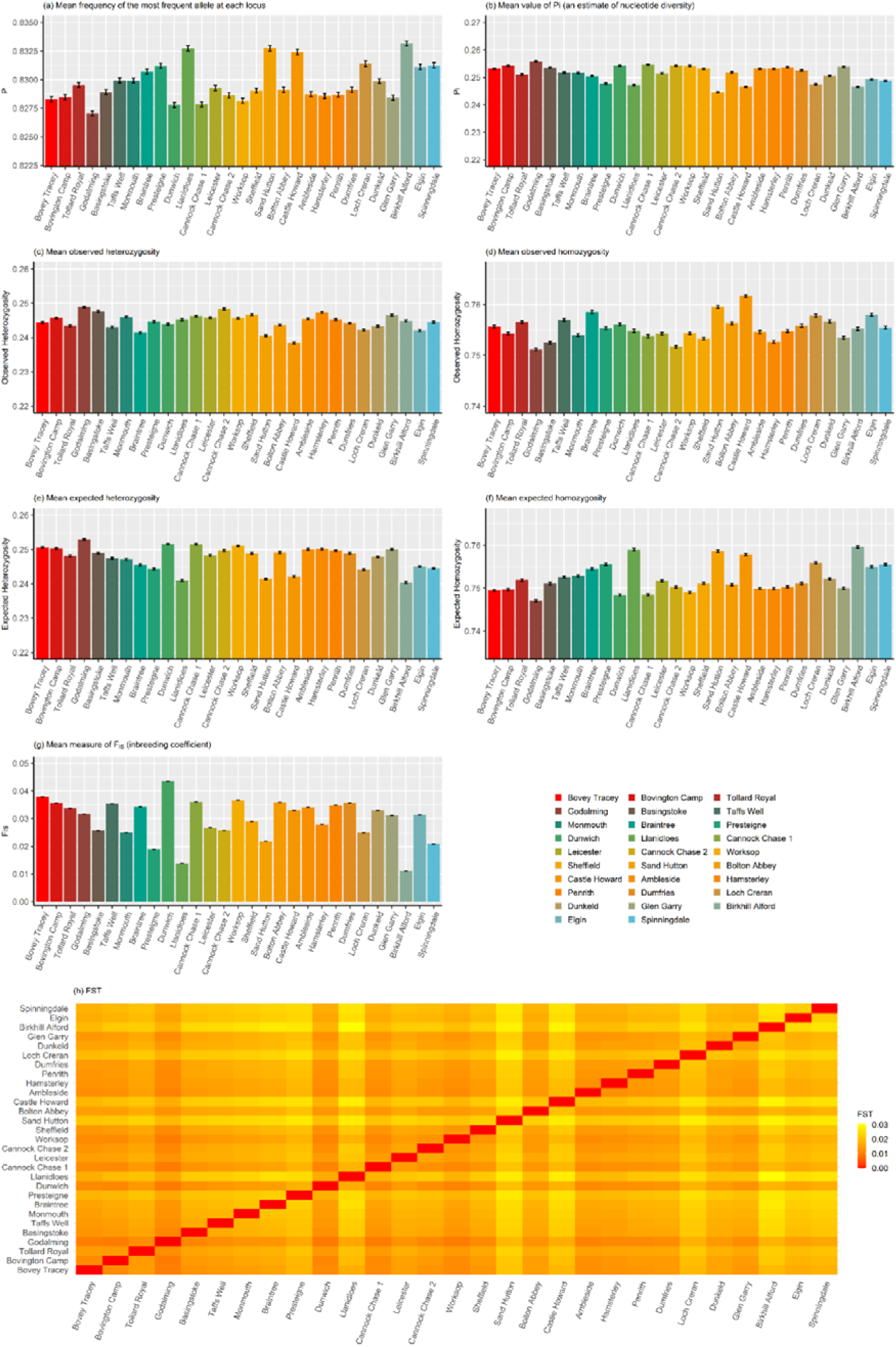
Population-level genetic diversity and between-population genetic differentiation in silver birch (*Betula pendula*) following removal of individuals related to first and second degree. **a,** mean frequency of the most frequent allele at each locus (P). **b,** mean value of Pi (nucleotide diversity, π). **c,** mean observed heterozygosity (Obs_Het). **d,** mean observed homozygosity (Obs_Hom). **e,** mean expected heterozygosity (Exp_Het). **f,** mean expected homozygosity (Exp_Hom). **g,** inbreeding coefficient (FIS). **h,** heatmap of pairwise fixation index (F_ST_) values between each pair of provenances. Provinces are ordered by ascending latitude. Note the differing y-axis scales for each plot. Analysis was performed using STACKS populations software on a related-filtered dataset of 1,020 samples from 29 provenances, on a random, 25% subset of LD-filtered SNPs (344,426 SNPs). It should be noted that π was calculated on biallelic variable sites only, which produces higher values than genome-wide per-base-pair estimates.

**Supplementary Figure 9:**
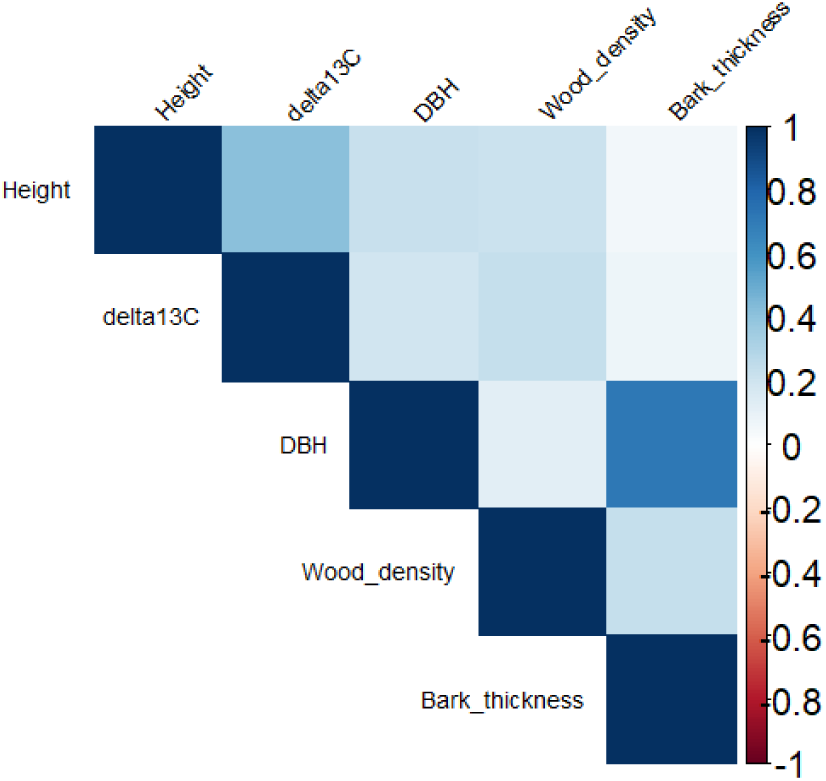
Pearson’s correlation coefficient between continuous phenotypic traits in silver birch (*Betula pendula*). Height = height in cm at 8 years after the establishment of the trials; DBH = diameter at breast height at age 19, measured in cm. δ^13^C= depletion of ^13^C isotope in parts per thousand, measured at age 20. Wood density = density proxy measured as detrended debarked drilling resistance. Bark thickness = bark thickness measured in cm at the exit of the microdrill needle.

**Supplementary Figure 10:**
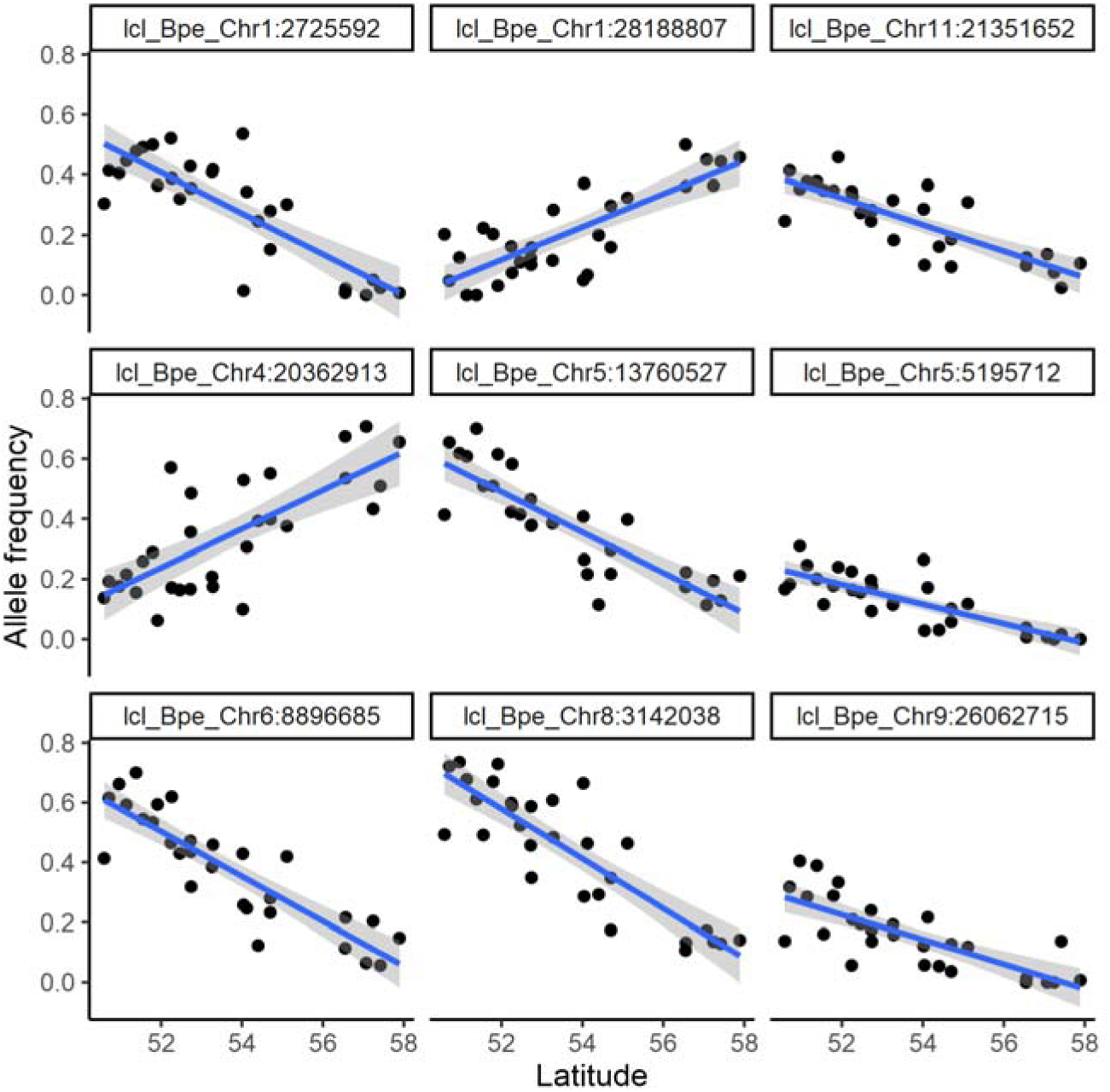
Relationship between latitude and allele frequencies of the top SNPs in peaks strongly associated with height in a GWAS not controlling for population structure and kinship. Allele frequencies were calculated for each provenance, and ordered by the provenance’s latitude of origin. Panel title refers to the chromosome identity followed by the SNP position in base pairs (e.g. lcl_Bpe_Chr1:2725592 refers to SNP at position 2,725,592 in chromosome 1). Blue lines represent the slopes of linear regressions and grey shades the confidence intervals.

**Supplementary Figure 11:**
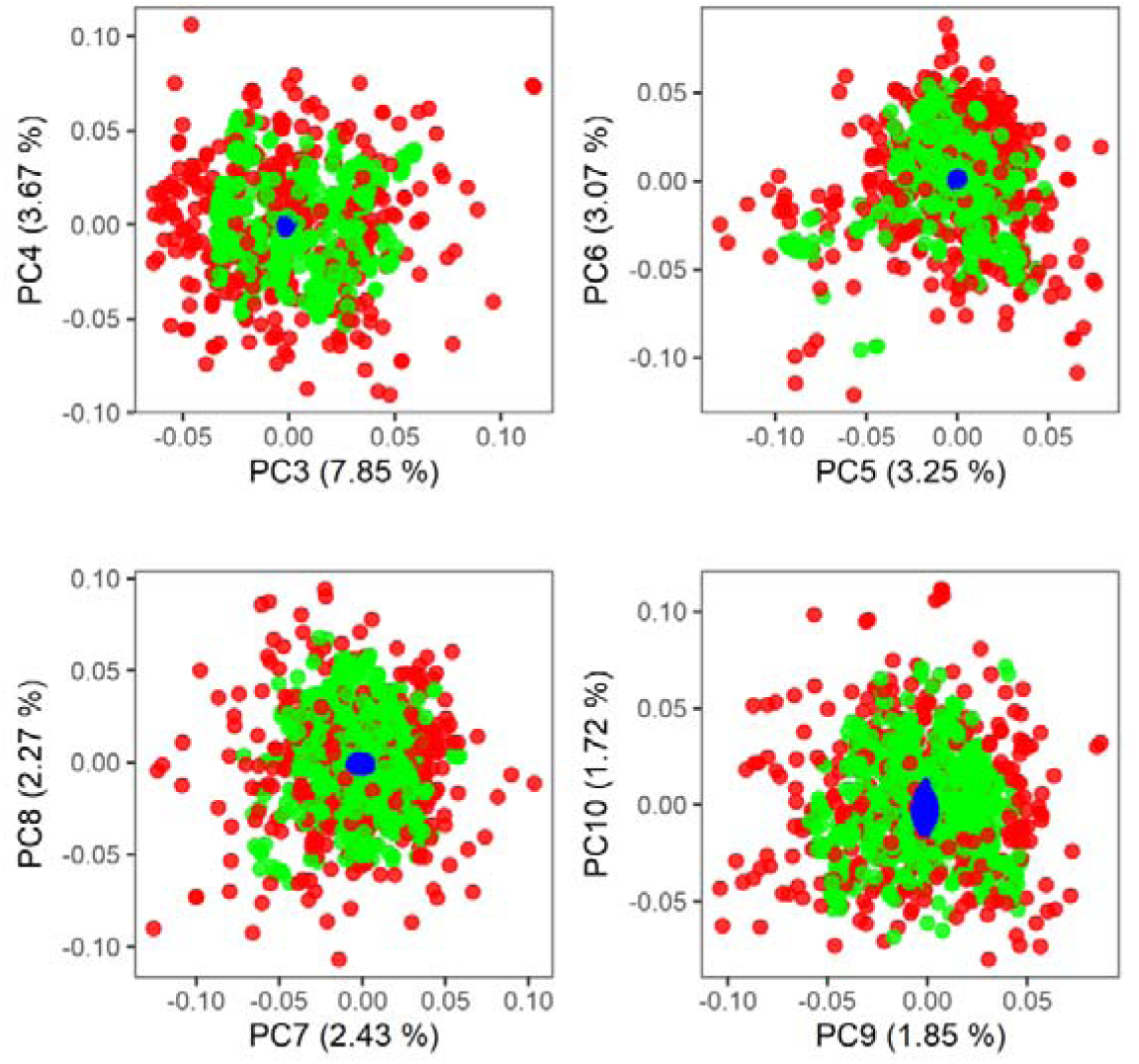
Genetic diversity in an ∼400kbp inverted region on chromosome 6. Panels depict different PC axes from PC3 to PC10. Dots are colour coded by haplotypes (red = no inversion, green = heterokaryotype for the inversion, blue = homokaryotype for the inversion, see Figure 4).

**Supplementary Figure 12:**
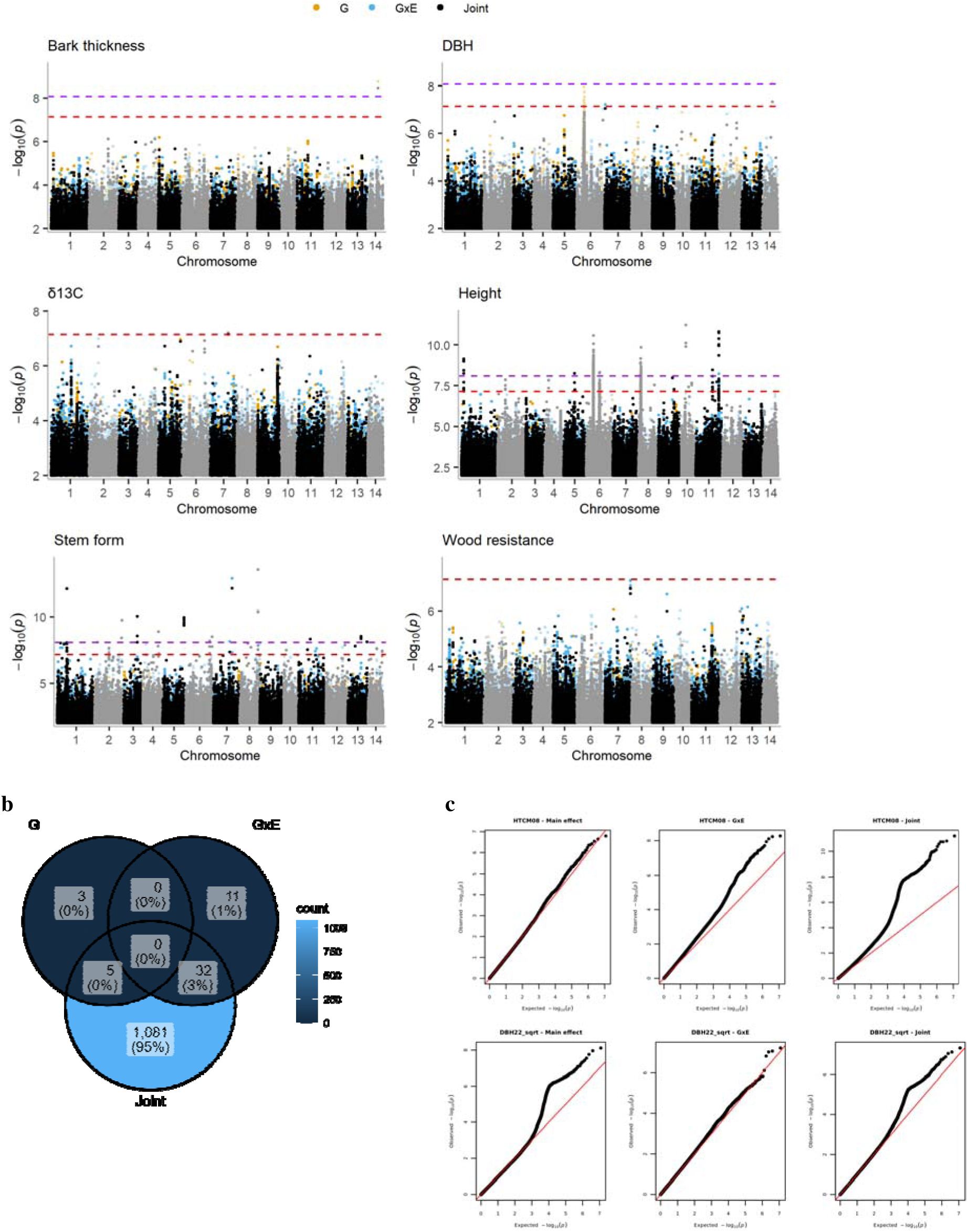
GWAS incorporating a genotype-by-environment interaction for multiple growth traits in silver birch (*Betula pendula*). **a,** Top panels depict Manhattan plots for each phenotype, coloured by model component (gold = main genetic effect, G; blue = interaction effect, GxE; and black = joint effect, Joint). **b,** Venn diagram categorising significant SNPs by model component. The blue gradient becomes lighter with an increasing number of SNPs. **c,** Q-Qplots for each model component for height (top three panels) and DBH (lower three panels). Inflation in p-values in the G component of DBH and joint component of height is mostly driven by the inversion in chromosome 6 (see main text).

**Supplementary Figure 13:**
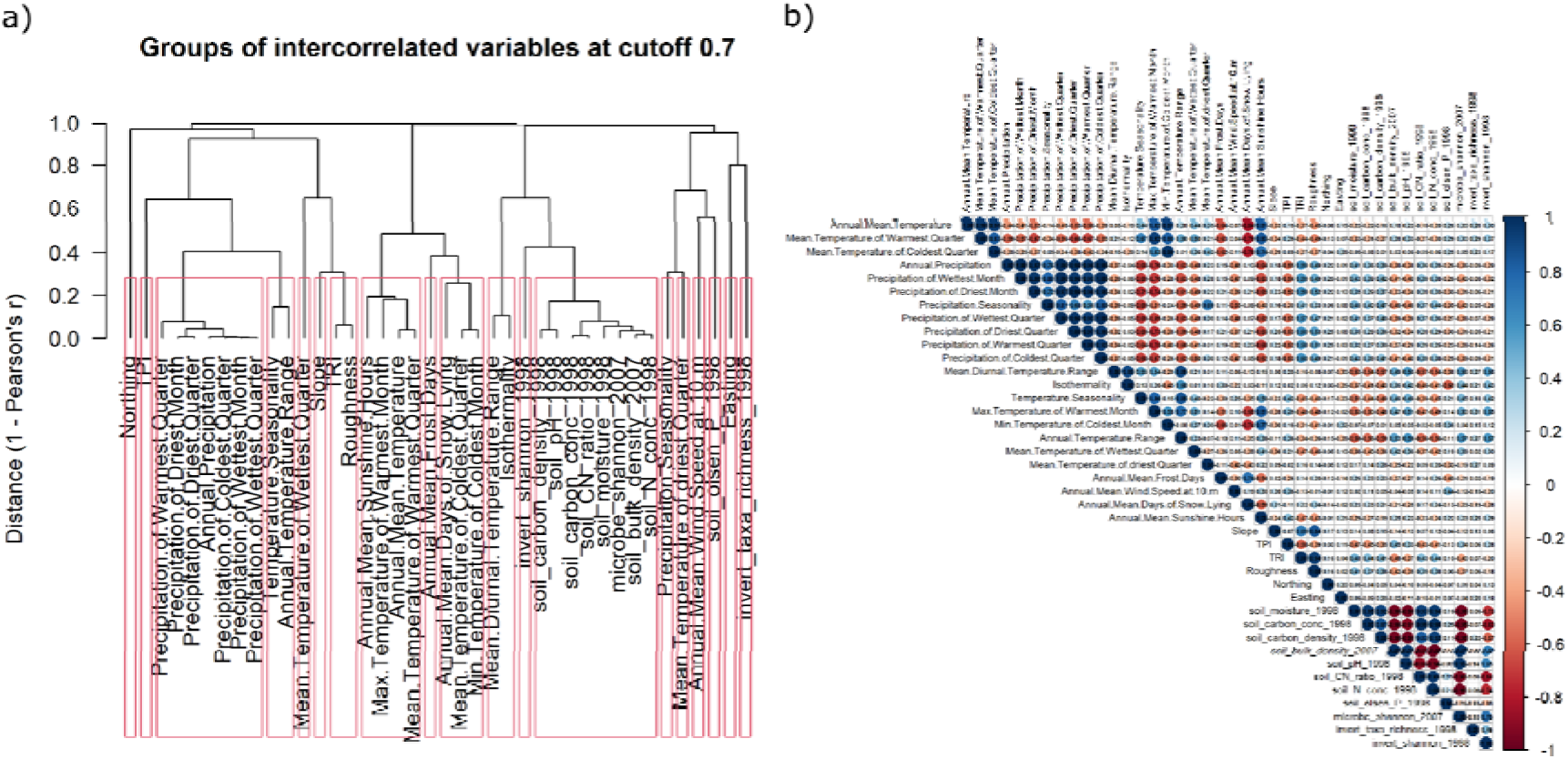
Correlation among 40 environmental, terrain and soil variables extracted at the source of 29 silver birch (*Betula pendula*) provenances. **a,** Plot produced by the R package *virtualspecies,* grouping variables with a correlation coefficient greater than 0.7. **b,** Pairwise correlation coefficients plotted using the R package *corrplot*. Variables include 19 CHELSA climatologies, 4 MetOffice variables, 7 terrain characteristics, and 11 soil attributes.

**Supplementary Figure 14:**
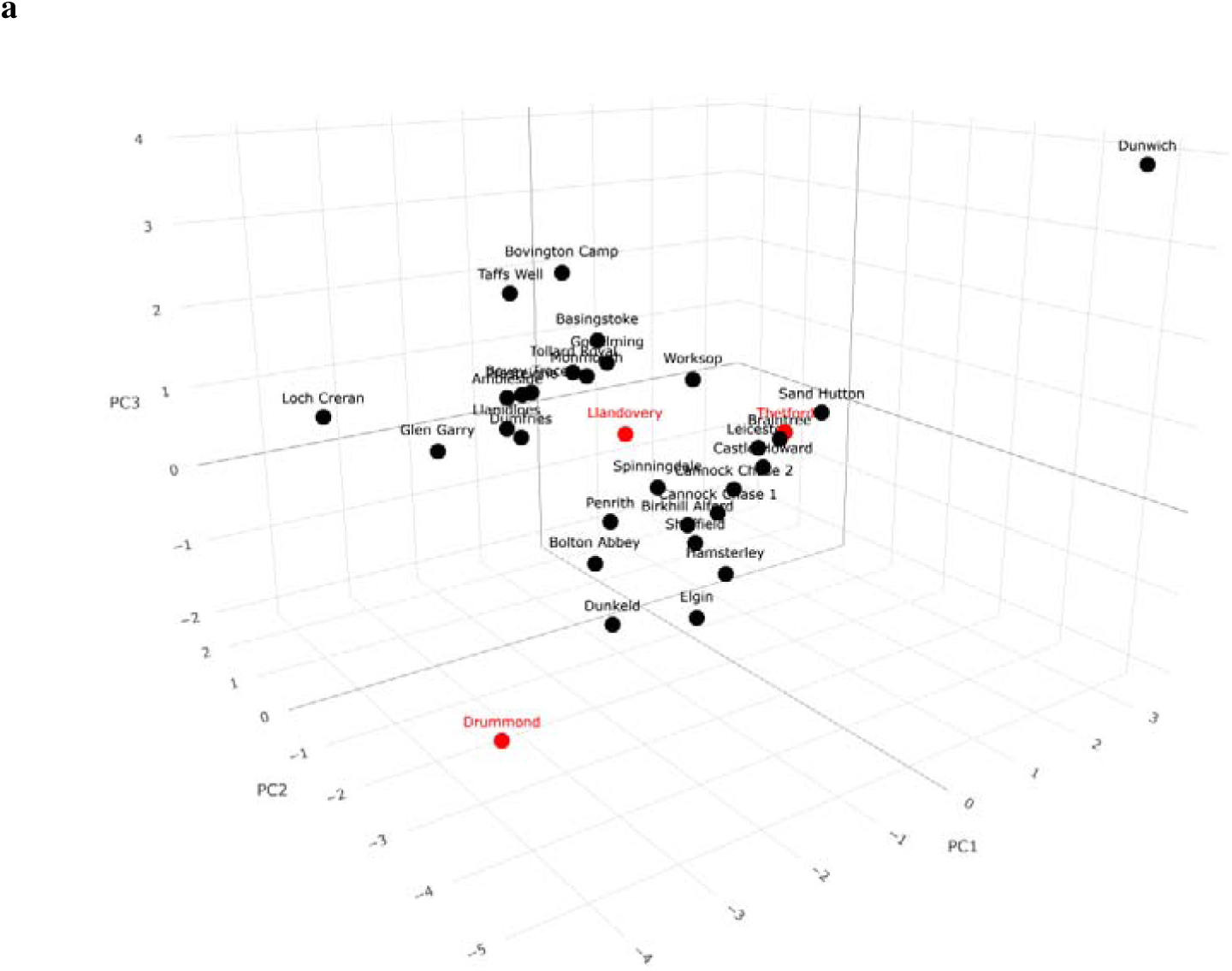
Environmental dissimilarity between trial sites and provenance sources. **a,** PCA summarising variation in 14 uncorrelated variables used in the GEA analysis (red = trial sites; black = provenances). **b,** scree plot of the percentage of variance explained by each PC.

**Supplementary Figure 15:**
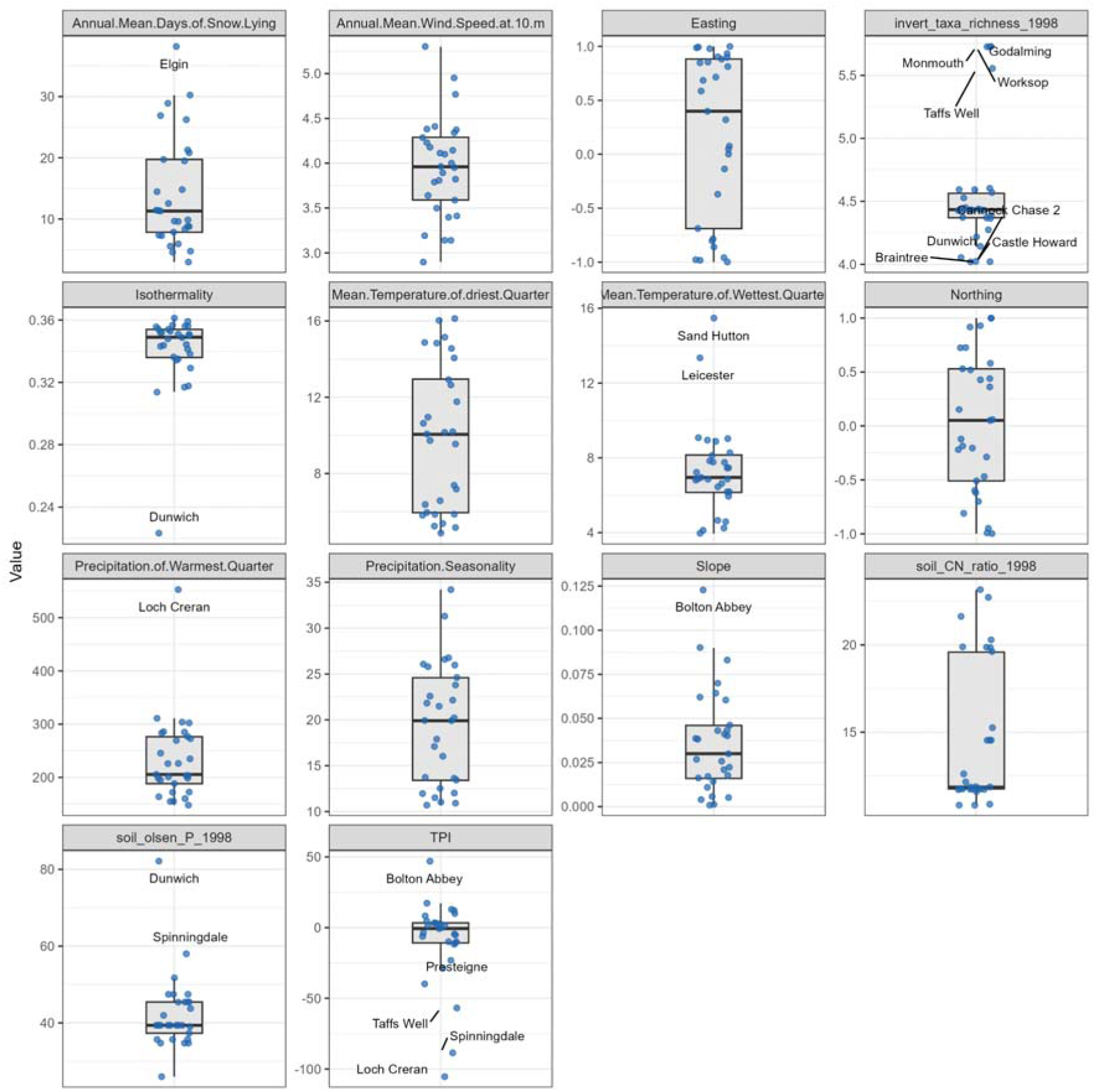
Distribution of the 14 environmental variables across the 29 UK *Betula pendula* provenances used in the GEA. Each point represents an individual provenance’s value for each variable, with provenances showing outlying values labelled (lowest and highest quartiles).

**Supplementary Figure 16:**
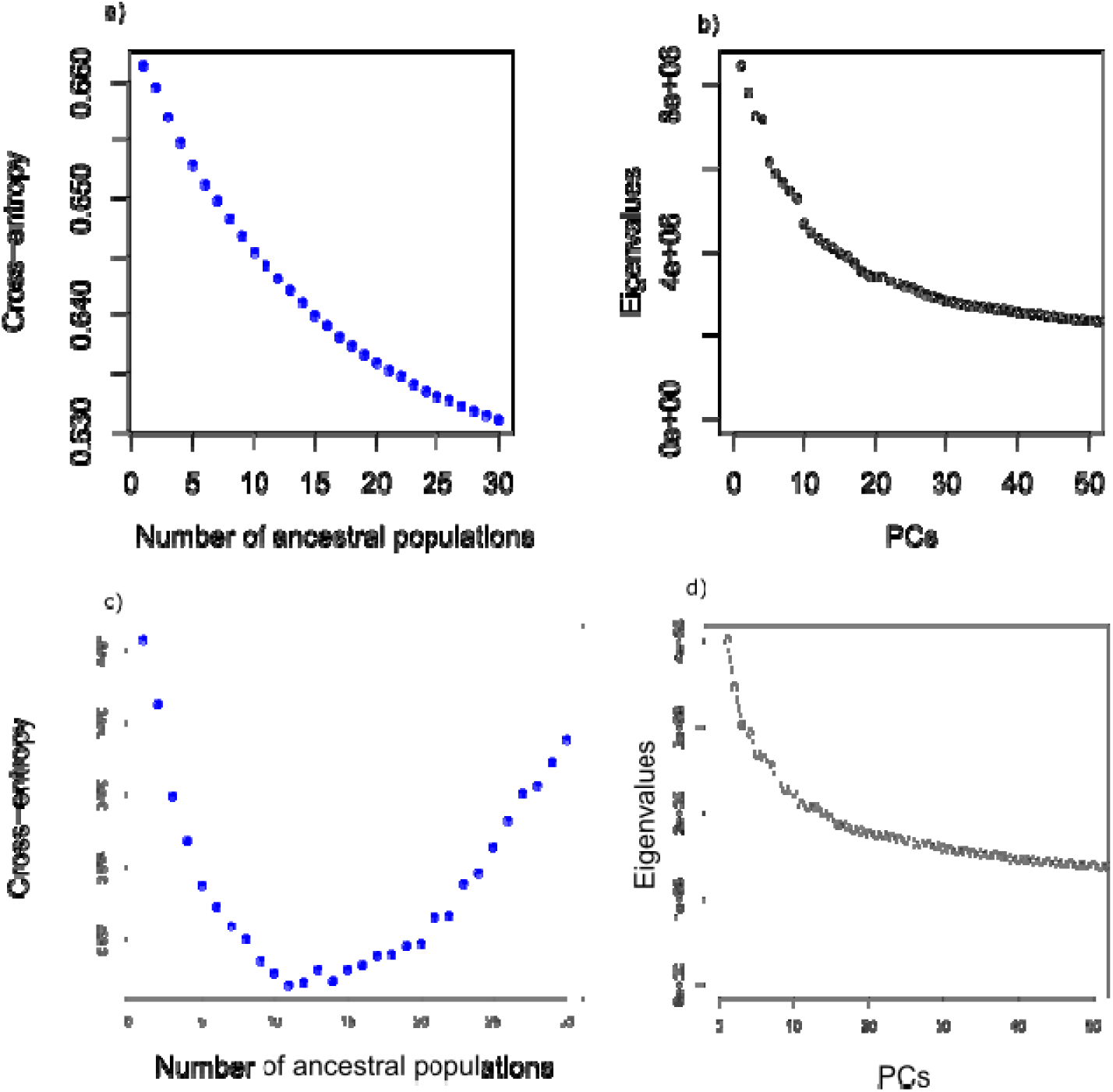
K value determination for Latent-Factor Mixed Model (LFMM) association testing. **a** and **b** show analysis performed on the full dataset of 1,873 individuals. **c** and **d** show analysis performed on 1020 individuals following removal of individuals related to first- or second-degree. **a** and **c** show the cross-entropy output of SNMF analysis run at K1-30 with 7 repetitions, indicating the likelihood of ancestral populations at explaining variance in the data. Lower cross-entropy values indicate better K value estimations. **b** and **d** show the plotting of eigenvalues following PCA, where the best estimate of ancestral populations is determined according to Catell’s rule (elbow of the curve). Analysis was performed in the LEA R package on a subset of SNPs following removal of variants in transposable elements and linkage disequilibrium to lessen computational demand, reducing the number of SNPs from ∼5.9 million to 738,833. For the full dataset, 30+ sources of variation contribute to the data. After removal of related individuals, K11 best accounts for latent structure.

**Supplementary Figure 17:**
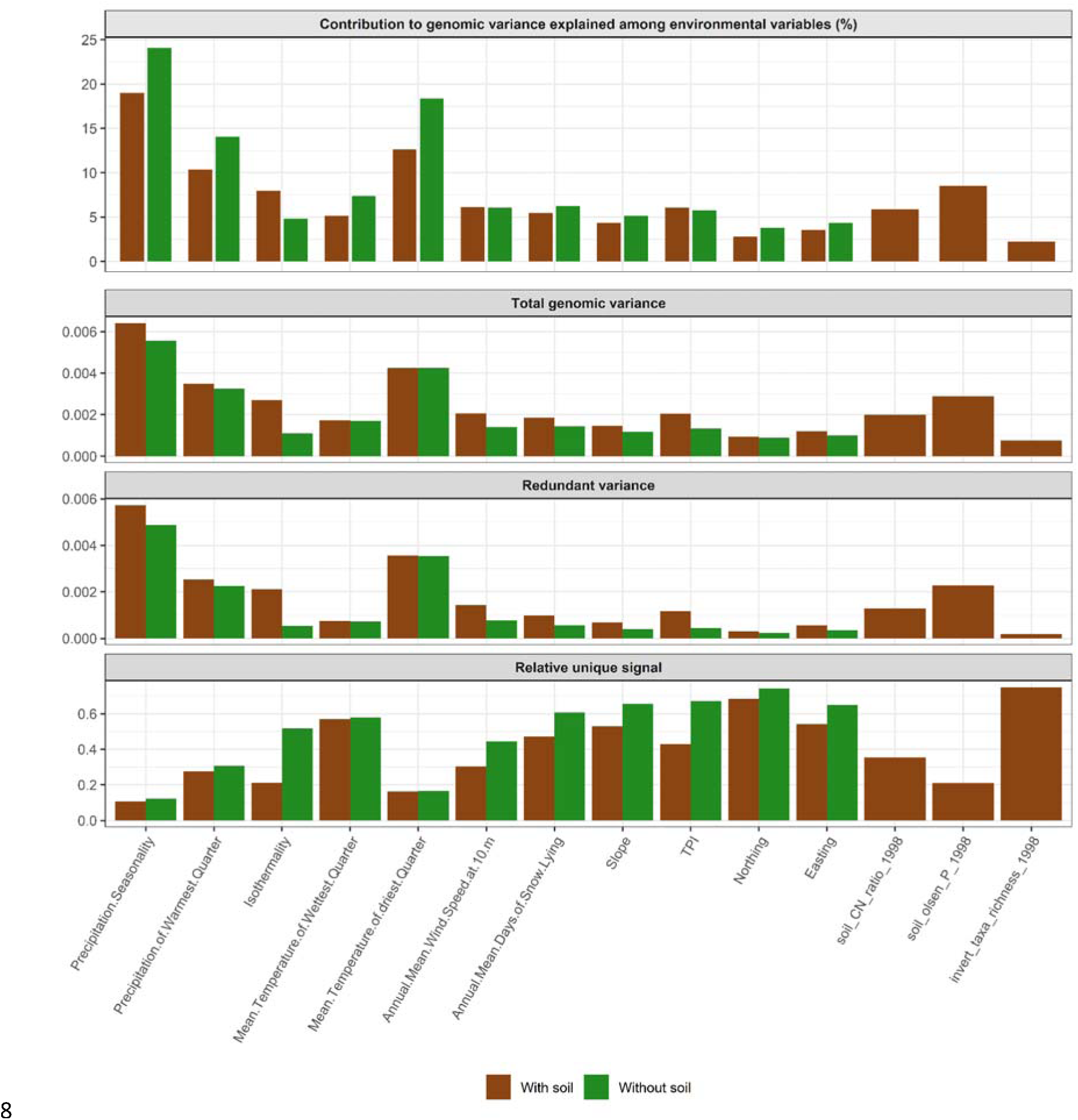
Assessment of the unique and shared contribution of each environmental variable to genetic variation with and without soil. Values were calculated from the effect size matrix (β) computed by lfmm2(), where **a,** shows the proportional contribution of each environmental variable to the total genomic variation explained across all other variables **b**, shows the raw total genomic variance of each variable. **c**, shows the redundant variance meaning the variance of each variable which can be explained by other variables, and **d,** shows the relative unique signal representing the truly independent genomic signal of each variable. Analysis was repeated with (brown bars) and without (green bars) soil data to investigate the effect of including the below ground component, and thus a greater number of variables, on the shared and independent genetic variance explained by each variable.

**Supplementary Figure 18:**
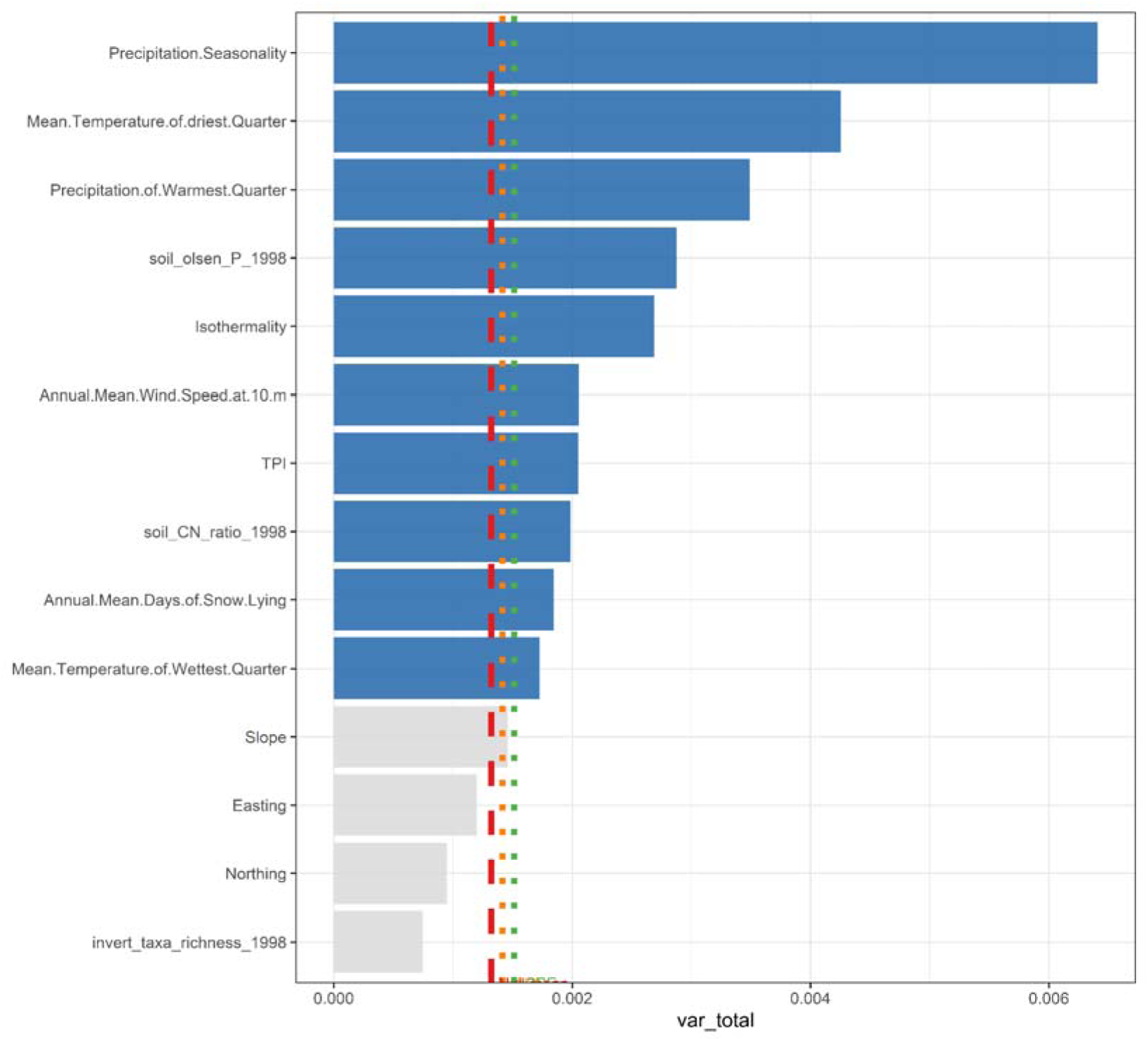
Variance in SNP effect sizes explained by each environmental variable in GEAs compared to null. The x-axis represents the total variance in SNP effect sizes extracted from lfmm2() models including 14 environmental variables (y-axis). The red dashed line represents the mean variance explained by 10 permuted dummy variables, and the orange and green dotted lines represent one and two standard errors from this mean, respectively. Variables explaining more variance than 2 SE from the null mean are coloured in blue and were considered significant.

**Supplementary Figure 19:**
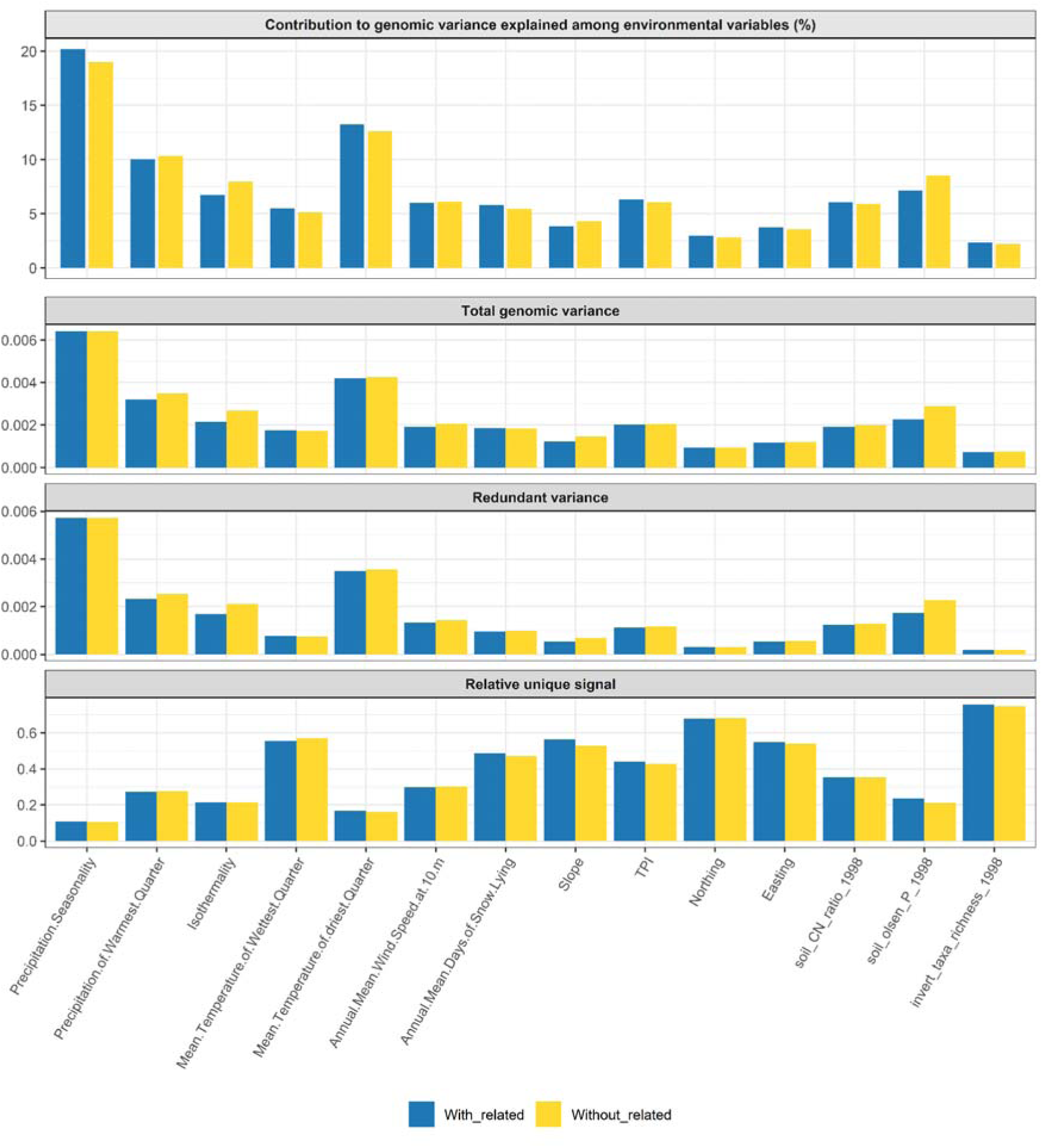
Assessment of the unique and shared contribution of each environmental variable to genetic variation with and without related individuals. Values were calculated from the effect size matrix (β) computed by lfmm2(), where **a,** shows the proportional contribution of each environmental variable to the total genomic variation explained across all other variables, **b,** shows the raw total genomic variance of each variable, **c,** shows the redundant variance as that which can be explained by other variables, and d, shows the relative unique signal representing the truly independent genomic signal of each variable. Analysis was repeated with related individuals (blue) and after the removal of ∼800 individuals related to second degree or higher (yellow) for interpretation of the possible confounding caused by within- and between-provenance relations.

**Supplementary Figure 20:**
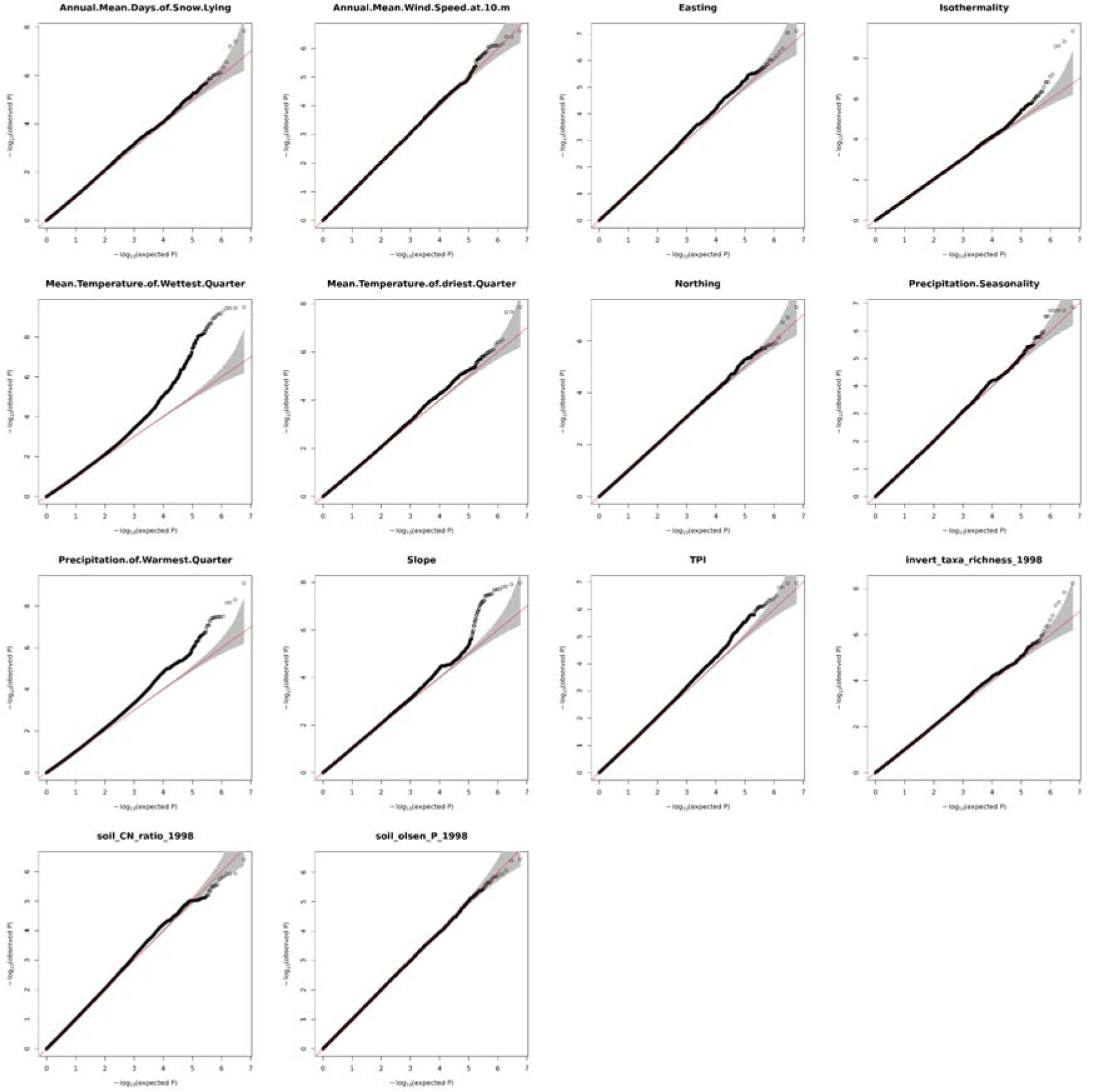
QQ-plots for 14 genome-environment associations in silver birch (*Betula pendula*). Corresponding Manhattan plots are shown in Extended Figure 5. GEAs were run with 1,020 individuals at K=11 using LFMM2().

**Supplementary Figure 21:**
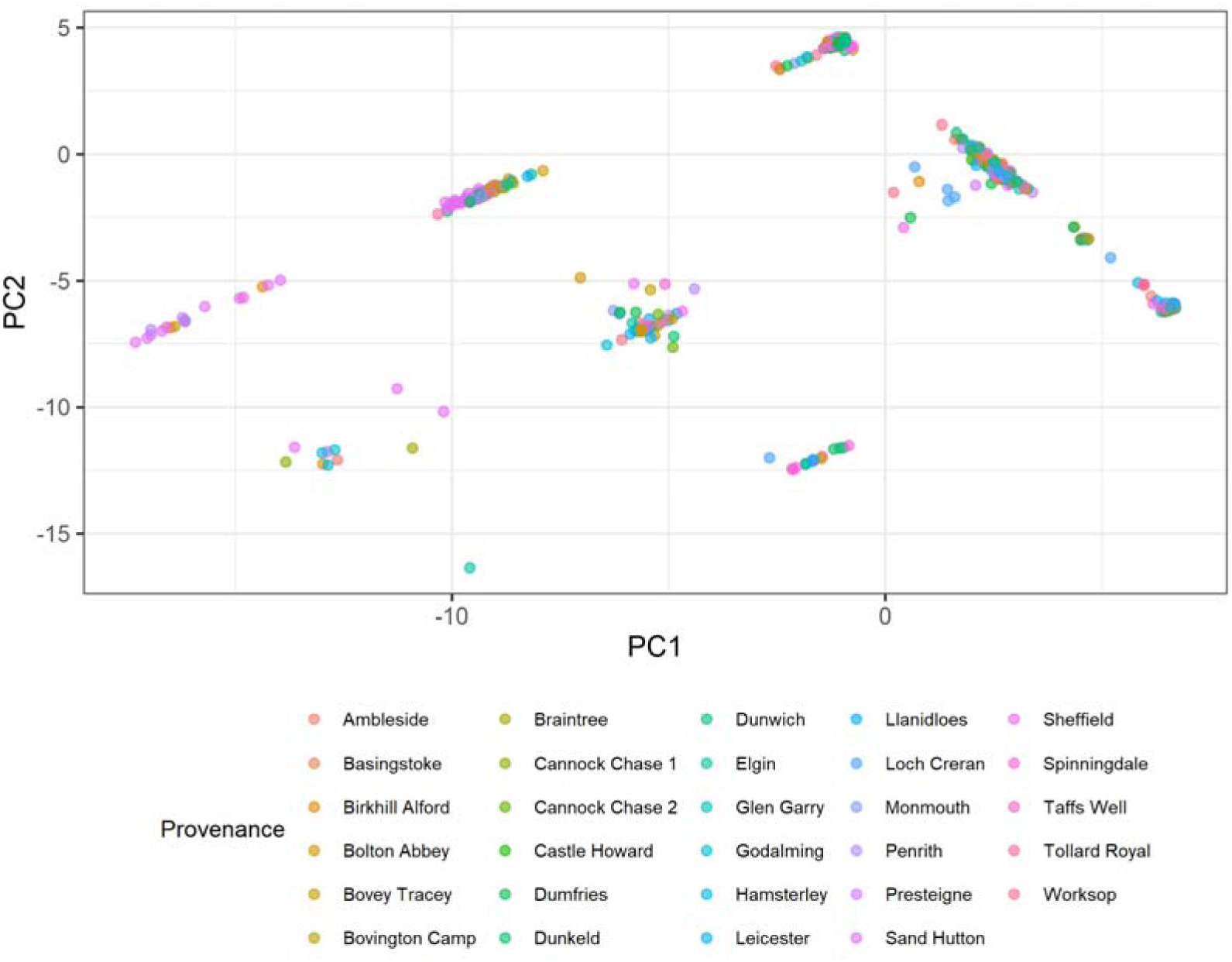
PCA of SNPs associated with environmental variables in *Betula pendula*. LFMMs identified 117 SNPs associated with 14 uncorrelated environmental variables. PCA reveals SNPs clustered in discrete blocks consistent with structural variation.

**Supplementary Figure 22:**
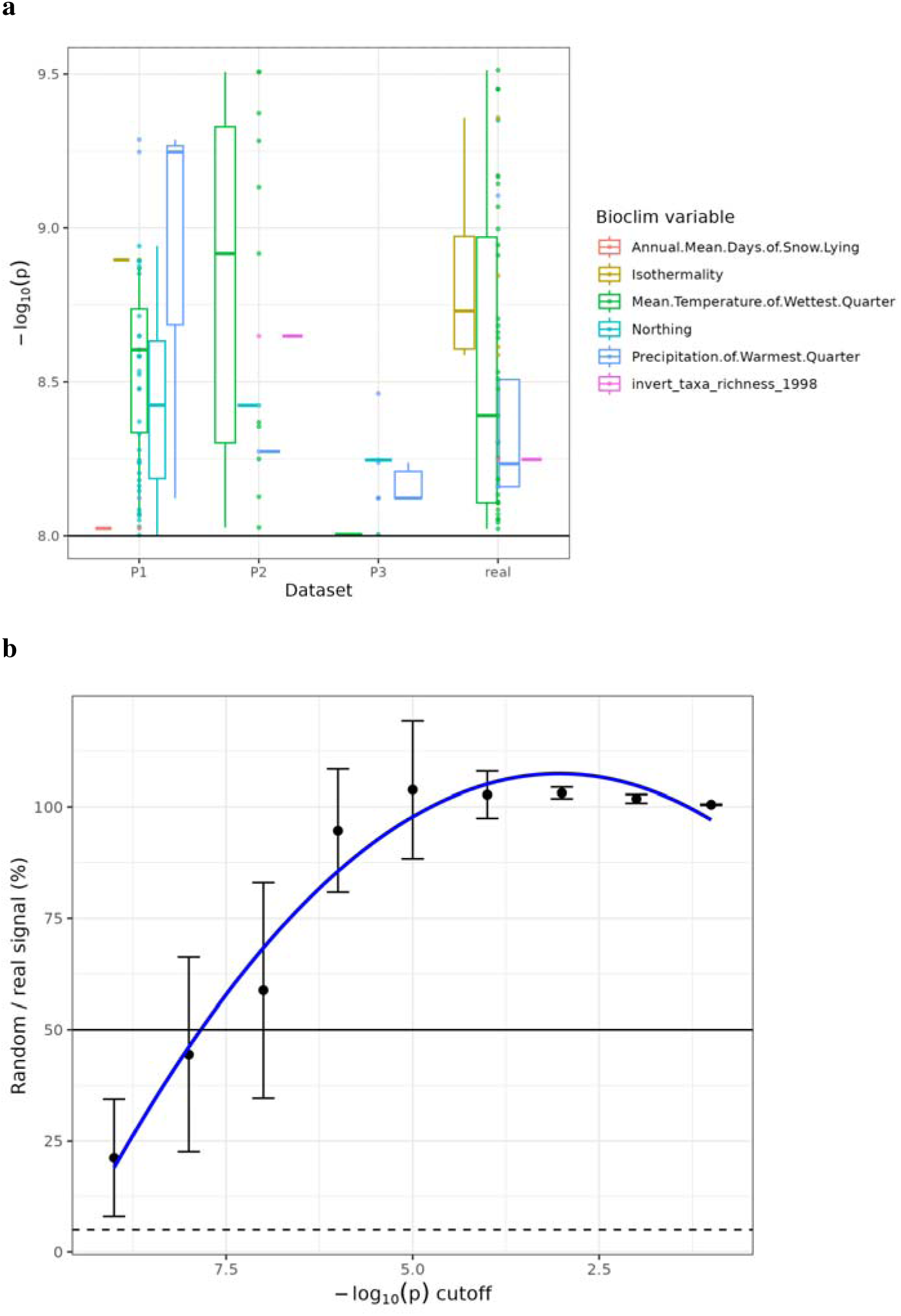
Genome-environment association signals compared with noise. **a,** box plots of - log10 p-values per environmental variable for which there were significant signals in the real data. The x-axis refers to each permutation run (P1, P2 and P3) and the real dataset (real). Only -log10 p-values above 8 are shown for simplicity; **b,** ratio of SNPs in the real and permuted dataset at certain bins of increasing -log10 p-values (see ref). Dots and bars represent the mean and standard errors for the 3 permutations, and the blue line shows a polynomial regression.

**Supplementary Figure 23:**
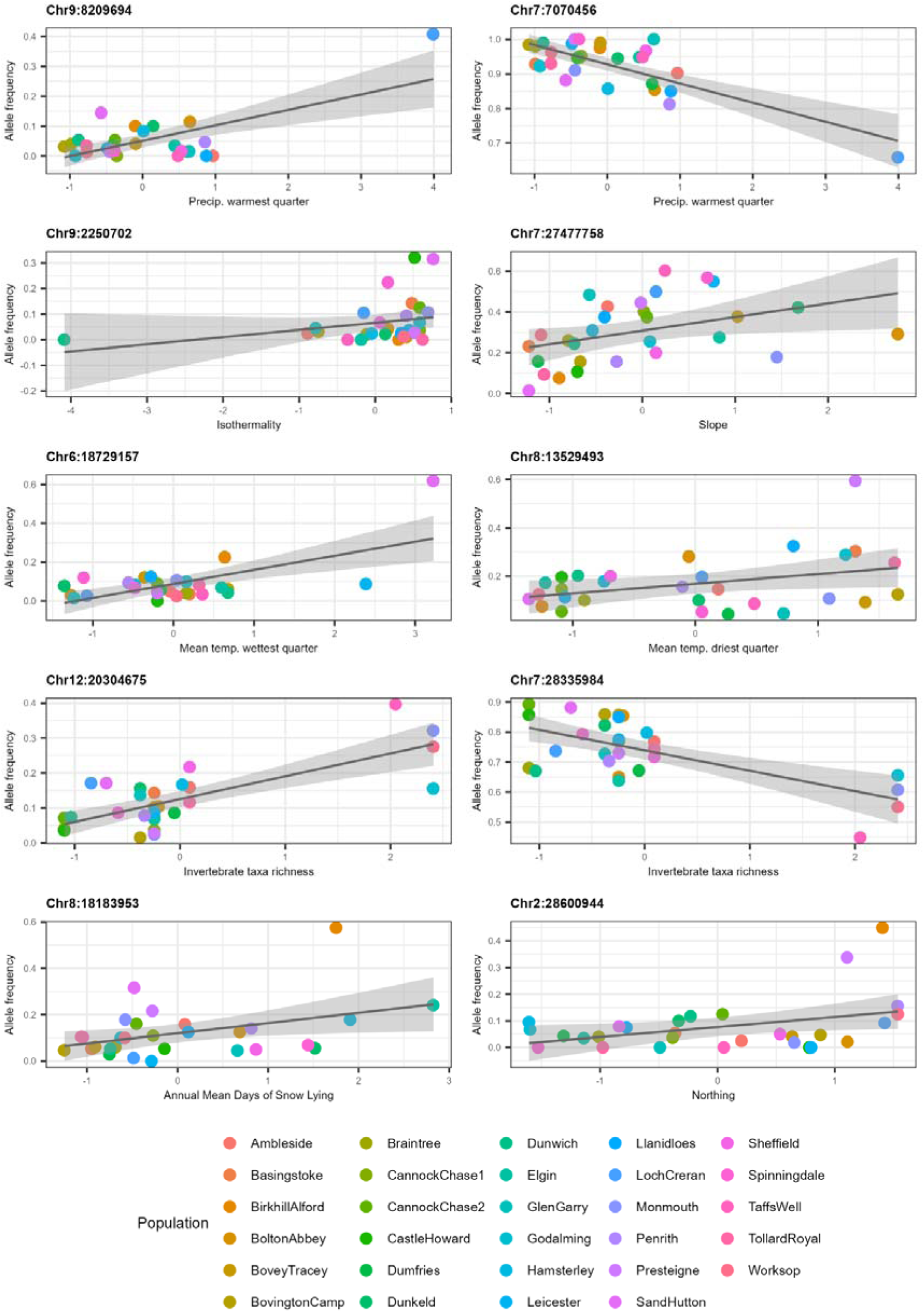
Population-level allele frequencies of the most significantly associated SNP for each environmental variable according to the modified Bonferroni significance threshold against the environmental conditions of the associated variable at source location. Allele frequencies for significant SNPs with the smallest p-value for each environmental variable were calculated using plink. SNPs identified within discrete genomic regions / chromosomes / peaks on Manhattan plots for one environmental variable are included as separate plots for individual interpretation.

**Supplementary Figure 24:**
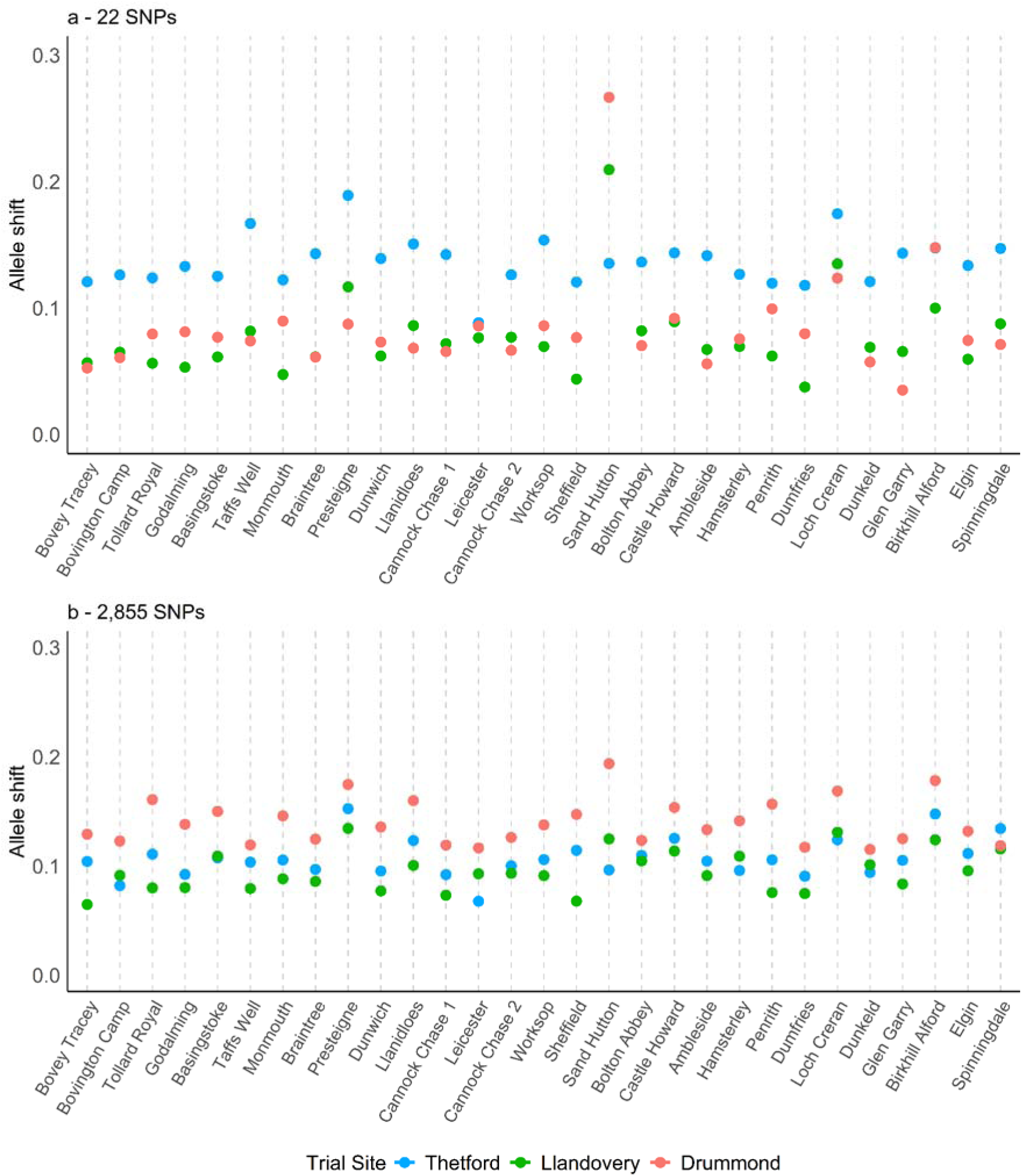
Predicted mean shift in allele frequencies necessary for local adaptation of each provenance to each trial site. Provenances are ordered by latitude. LFMMs were run on 1020 individuals across 29 provenances to 14 environmental, terrain and soil variables resulting in 117 modified Bonferroni-significant loci. Alleleshift analysis was performed on **a,** a set of 22 unlinked climate-associated loci after reducing the 117significant SNPs based on linkage, and **b,** a set of 2,855 unlinked SNPs after filtering the top 10,000 ranked by -log10pvalue based on linkage. Allele shifts were calculated by subtracting the predicted allele frequency at each trial site from the allele frequency at each original provenance location.

**Supplementary Figure 25:**
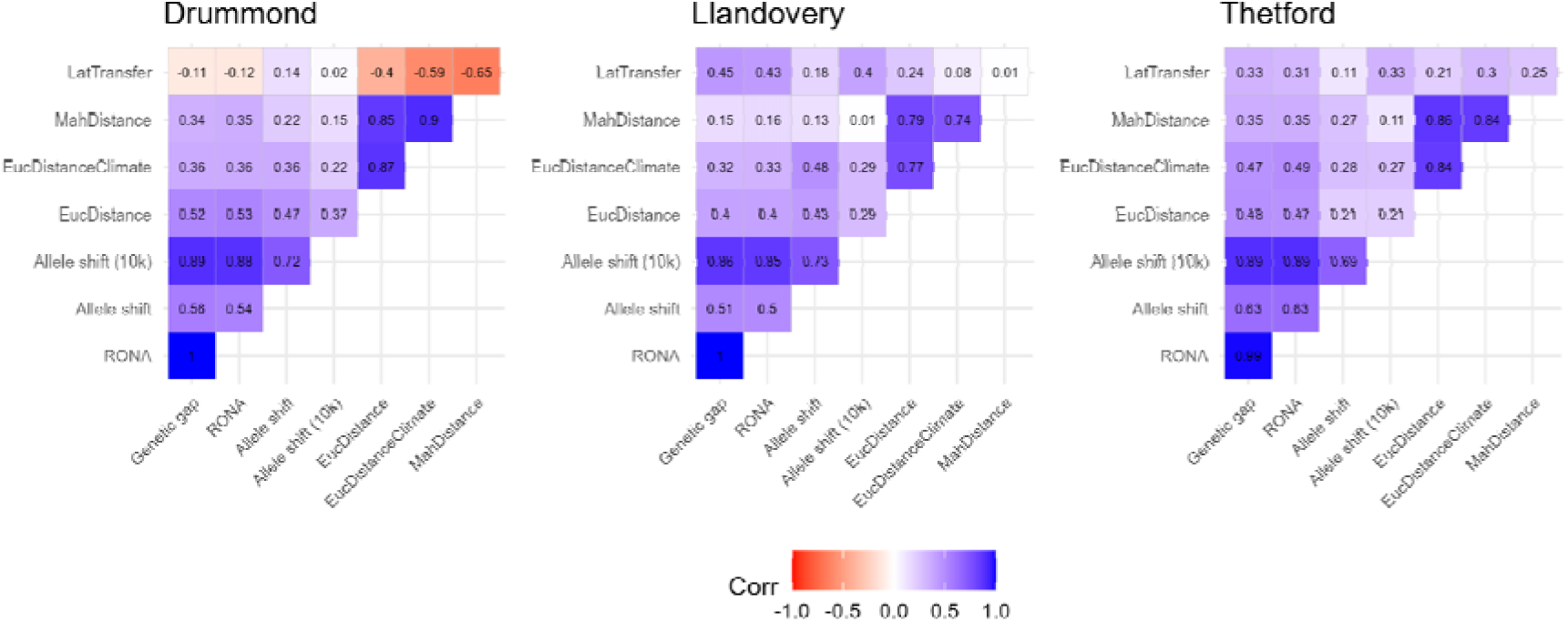
Correlation between estimates of genetic offset, environmental dissimilarity, and latitudinal transfer between provenance and trial site local conditions. Correlations were performed independently by site (left panel = Drummond; center panel = Llandovery; Right panel = Thetford). Note: EucDistance = euclidean distance; EucDistanceClimate = euclidean distances restricted to climatic variables; MahDistance = Mahalanobis distance; Allele Shift 10k = allele shifts calculated with the top 2.8K SNPs by p-value identified in a GEA following filtering the top 10K SNPs for LD; Allele shift = allele shifts calculated with SNPs above the significant threshold in GEAs; RONA = modified risk of non-adaptedness. For details of how these estimates were calculated, please see Online Methods.

**Supplementary Figure 26:**
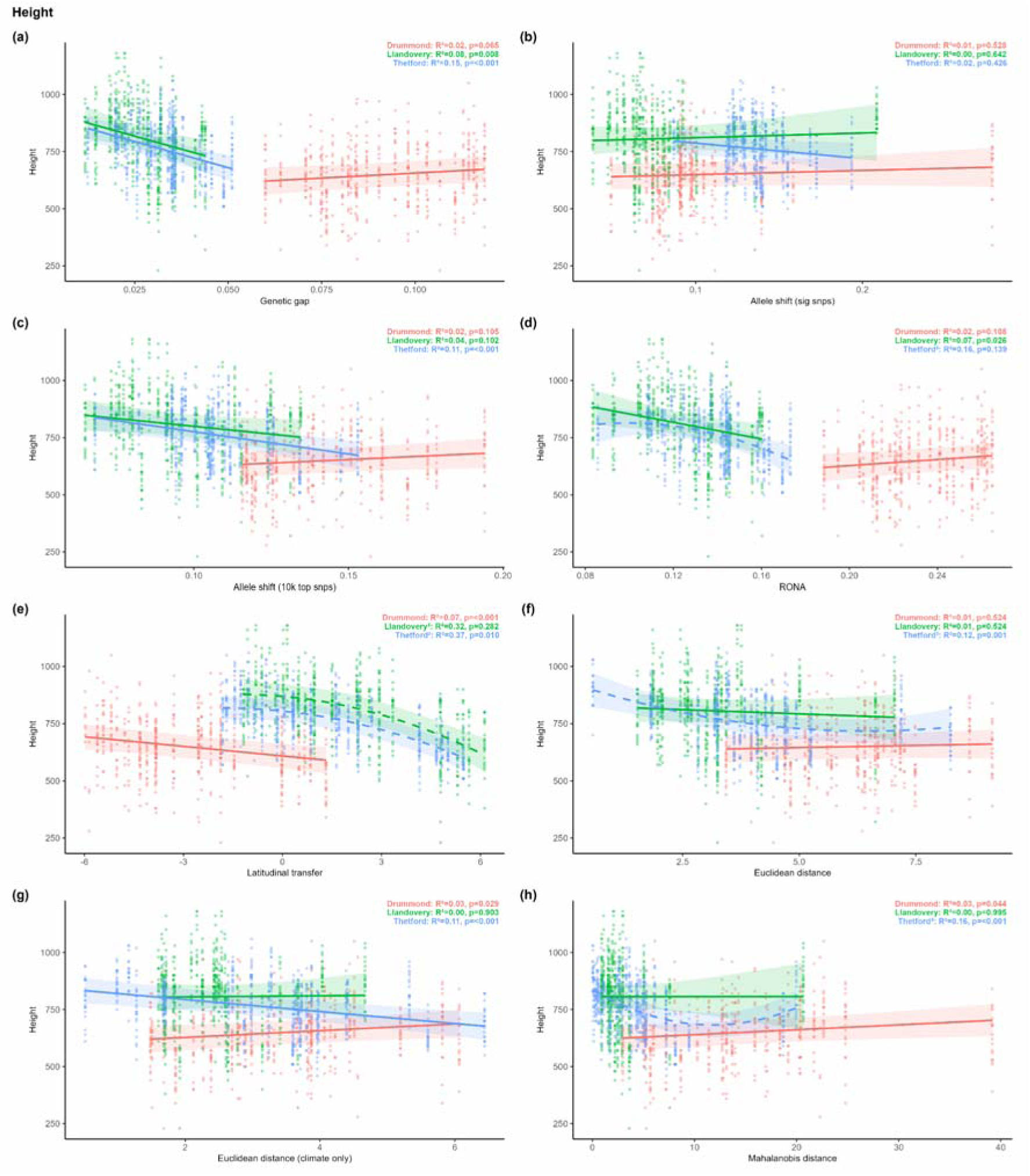
Relationship between height after 8 years’ growth and (a-d) measures of genetic offset (e-f) measures of environmental distance. **a,** Genetic gap calculated with LEA using ∼739K SNPs, **b,** allele shift calculated with 22 SNPs, **c,** allele shift calculated with 10,000 SNPs (reduced to ∼2.8K following LD filtering), **d,** RONA calculated with LEA using ∼739K SNPs **e,** latitudinal transfer, **f,** euclidean distance with all environmental variables **g,** euclidean distance without soil variables, **h,** mahalanobis distance.

**Supplementary Figure 27:**
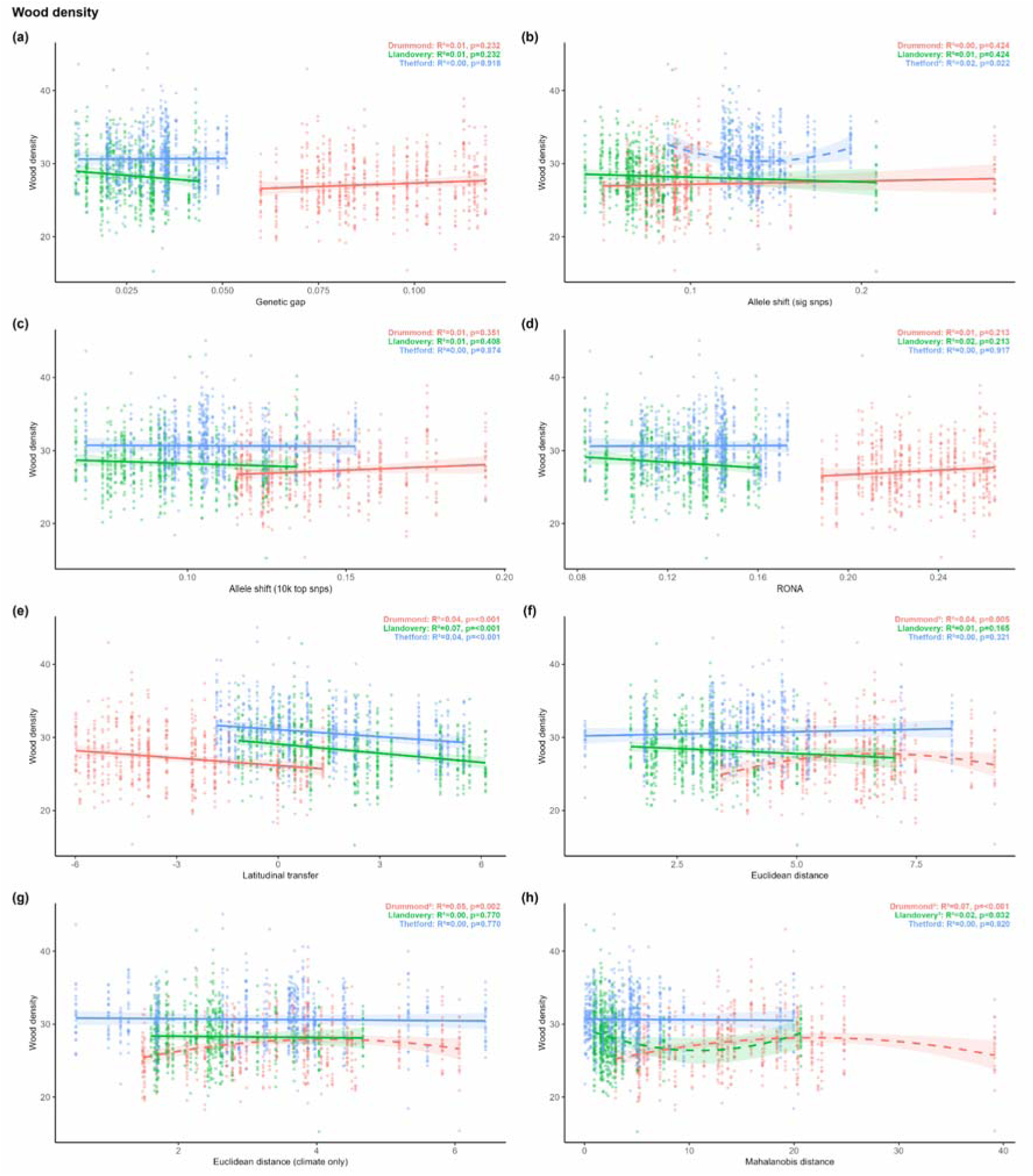
Relationship between wood density after 20 years’ growth and (a-d) measures of genetic offset (e-f) measures of environmental distance. **a,** Genetic gap calculated with LEA using ∼739K SNPs, **b,** allele shift calculated with 22 SNPs, **c,** allele shift calculated with 10,000 SNPs (reduced to ∼2.8K following LD filtering), **d,** RONA calculated with LEA using ∼739K SNPs **e,** latitudinal transfer, **f,** euclidean distance with all environmental variables **g,** euclidean distance without soil variables, **h,** mahalanobis distance.

**Supplementary Figure 28:**
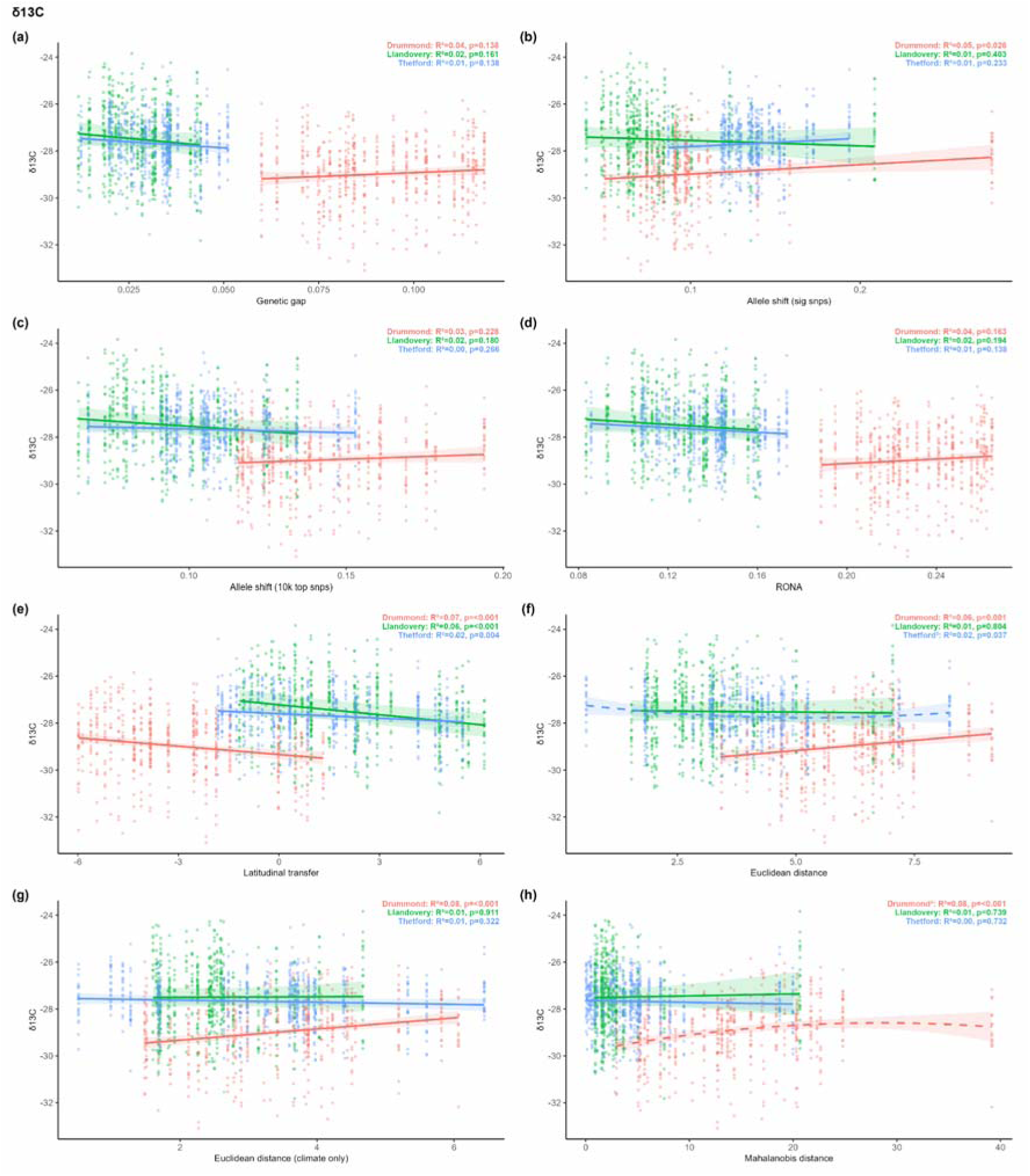
Relationship between carbon isotope ratio after 20 years’ growth and (a-d) measures of genetic offset (e-f) measures of environmental distance. **a,** Genetic gap calculated with LEA using ∼739K SNPs, **b,** allele shift calculated with 22 SNPs, **c,** allele shift calculated with 10,000 SNPs (reduced to ∼2.8K following LD filtering), **d,** RONA calculated with LEA using ∼739K SNPs **e,** latitudinal transfer, **f,** euclidean distance with all environmental variables **g,** euclidean distance without soil variables, **h,** mahalanobis distance.

**Supplementary Figure 29:**
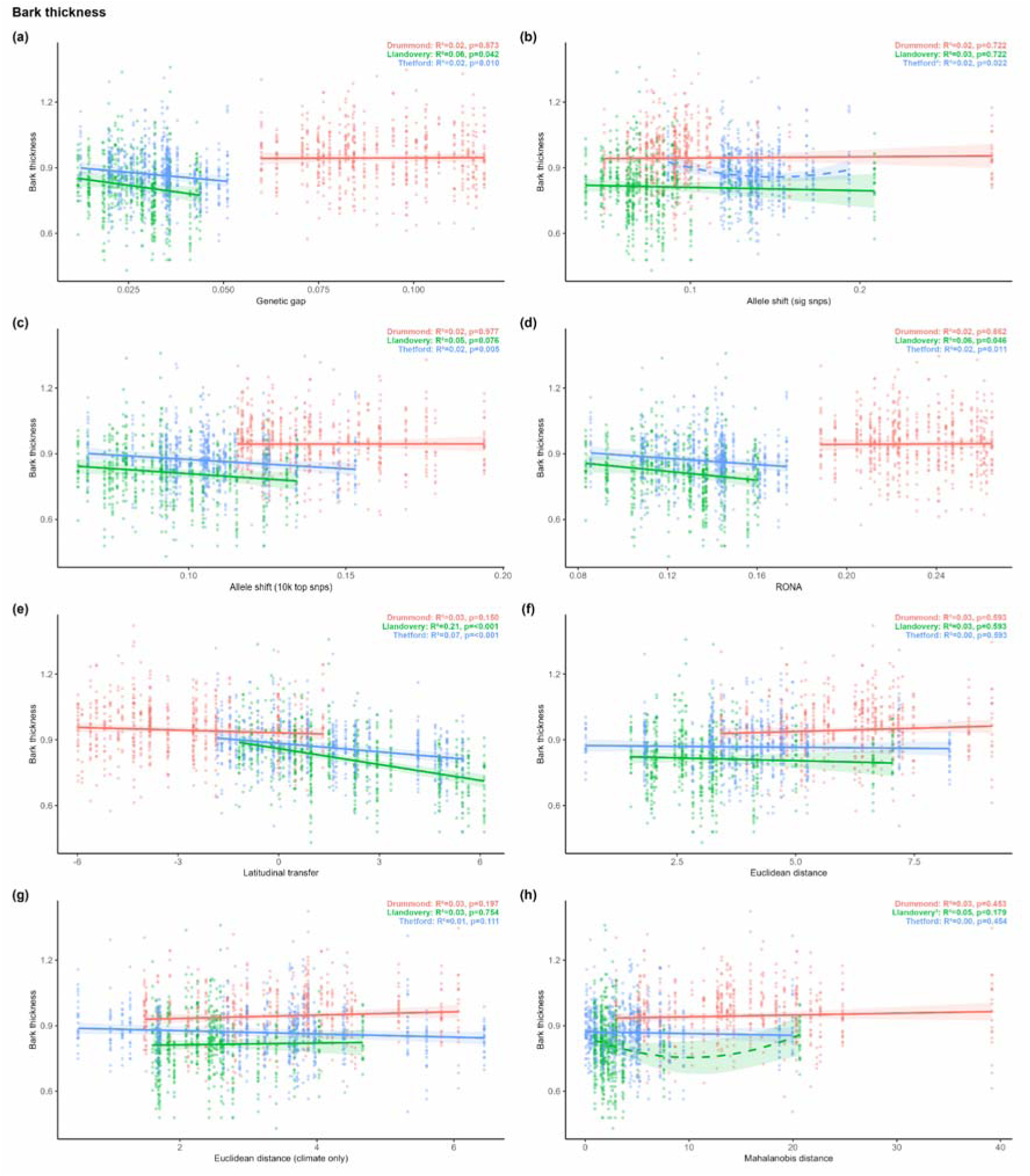
Relationship between bark thickness after 20 years’ growth and (a-d) measures of genetic offset (e-f) measures of environmental distance. **a,** Genetic gap calculated with LEA using ∼739K SNPs, **b,** allele shift calculated with 22 SNPs, **c,** allele shift calculated with 10,000 SNPs (reduced to ∼2.8K following LD filtering), **d,** RONA calculated with LEA using ∼739K SNPs **e,** latitudinal transfer, **f,** euclidean distance with all environmental variables **g,** euclidean distance without soil variables, **h,** mahalanobis distance.

**Supplementary Figure 30:**
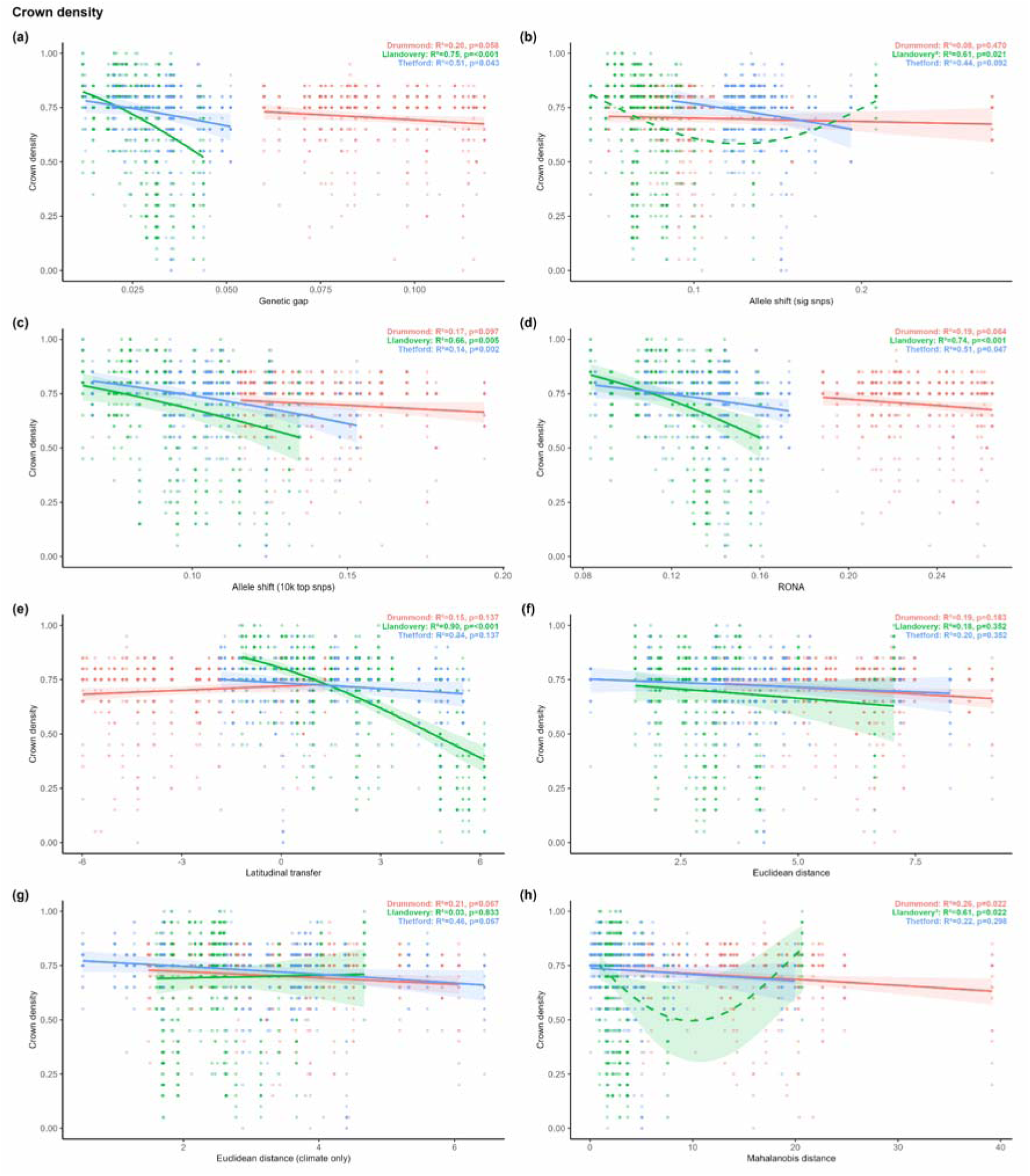
Relationship between crown density after 20 years’ growth and (a-d) measures of genetic offset (e-f) measures of environmental distance. **a,** Genetic gap calculated with LEA using ∼739K SNPs, **b,** allele shift calculated with 22 SNPs, **c,** allele shift calculated with 10,000 SNPs (reduced to ∼2.8K following LD filtering), **d,** RONA calculated with LEA using ∼739K SNPs **e,** latitudinal transfer, **f,** euclidean distance with all environmental variables **g,** euclidean distance without soil variables, **h,** mahalanobis distance.

**Supplementary Figure 31:**
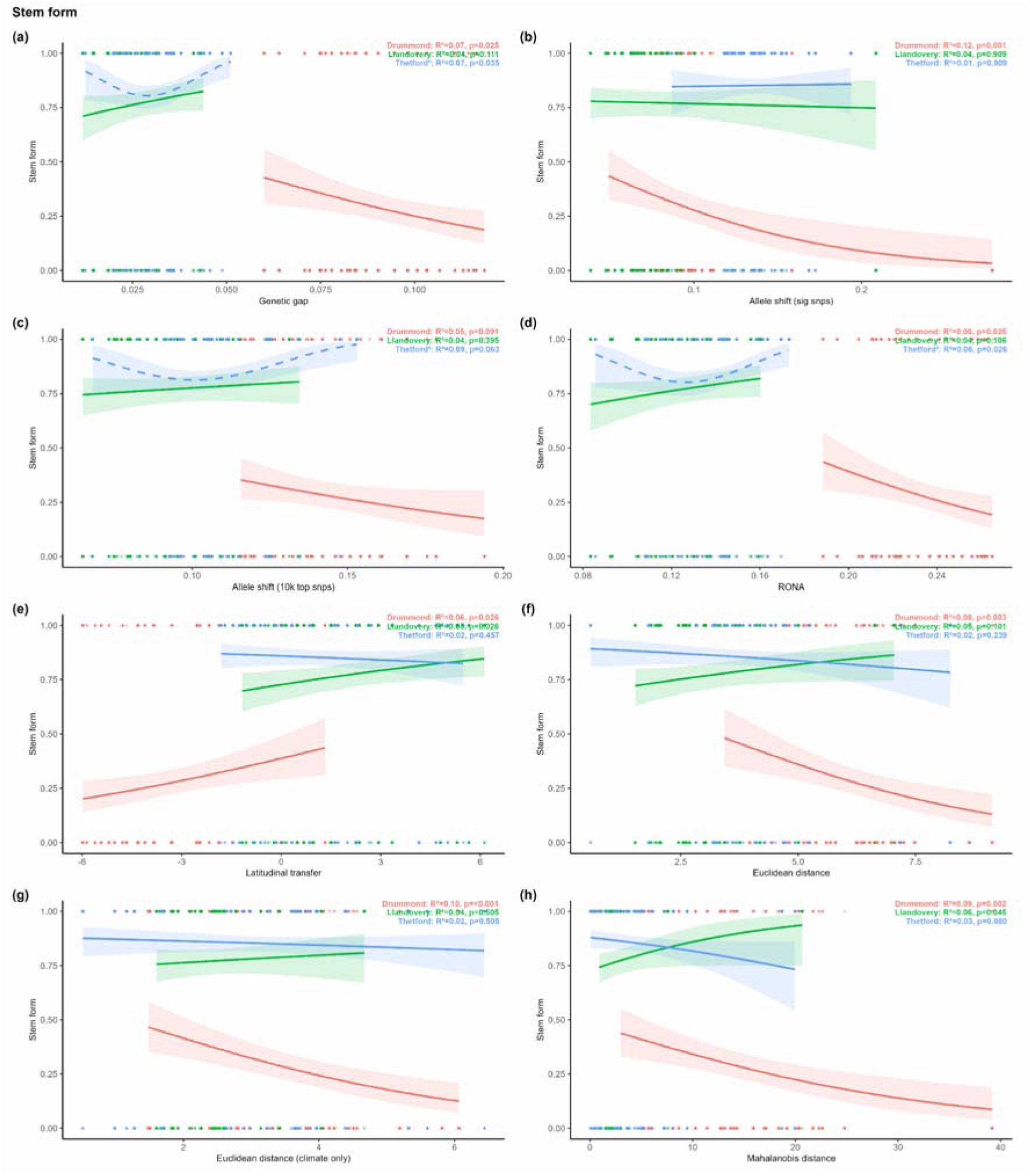
Relationship between stem form after 20 years’ growth and (a-d) measures of genetic offset (e-f) measures of environmental distance. **a,** Genetic gap calculated with LEA using ∼739K SNPs, **b,** allele shift calculated with 22 SNPs, **c,** allele shift calculated with 10,000 SNPs (reduced to ∼2.8K following LD filtering), **d,** RONA calculated with LEA using ∼739K SNPs **e,** latitudinal transfer, **f,** euclidean distance with all environmental variables **g,** euclidean distance without soil variables, **h,** mahalanobis distance.

**Supplementary Figure 32:**
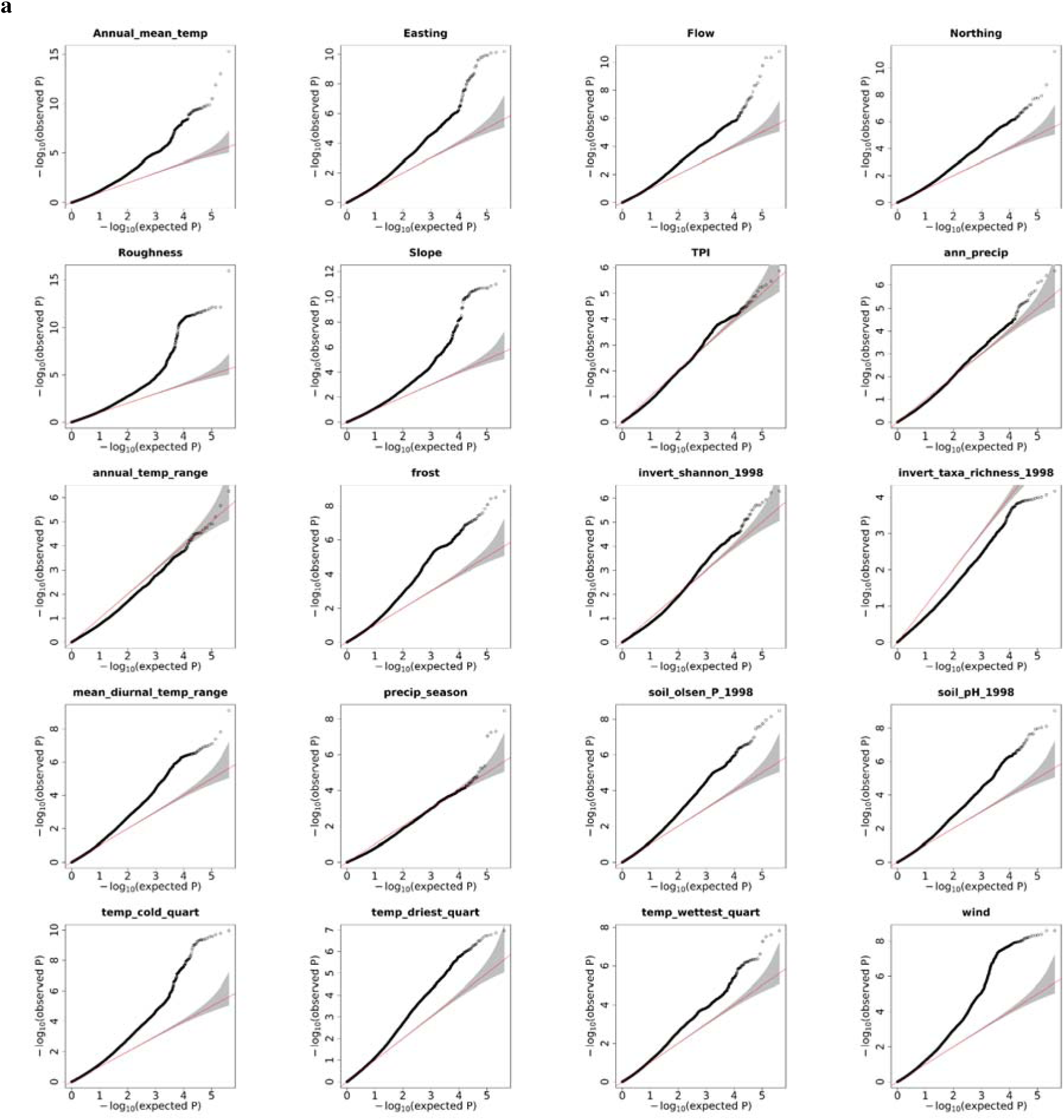
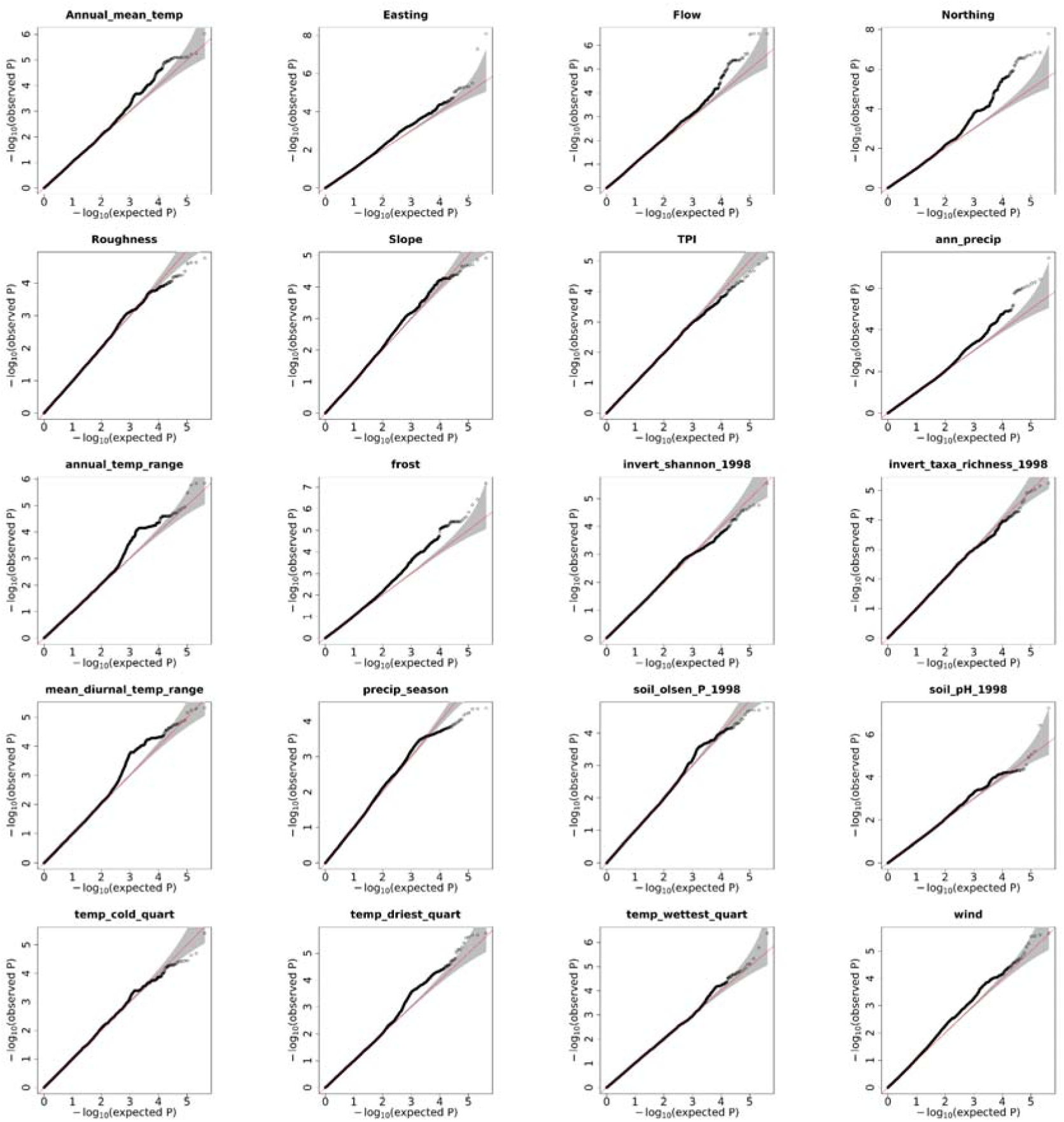
QQ-plots comparing a, lfmm() and b, lfmm2() algorithms. Analysis was performed on 1020 individuals at K=11 with genomic control using 420,165 SNPs present in chromosome nine to identify associations with 20 environmental, terrain and soil variables.

**Supplementary Figure 33:**
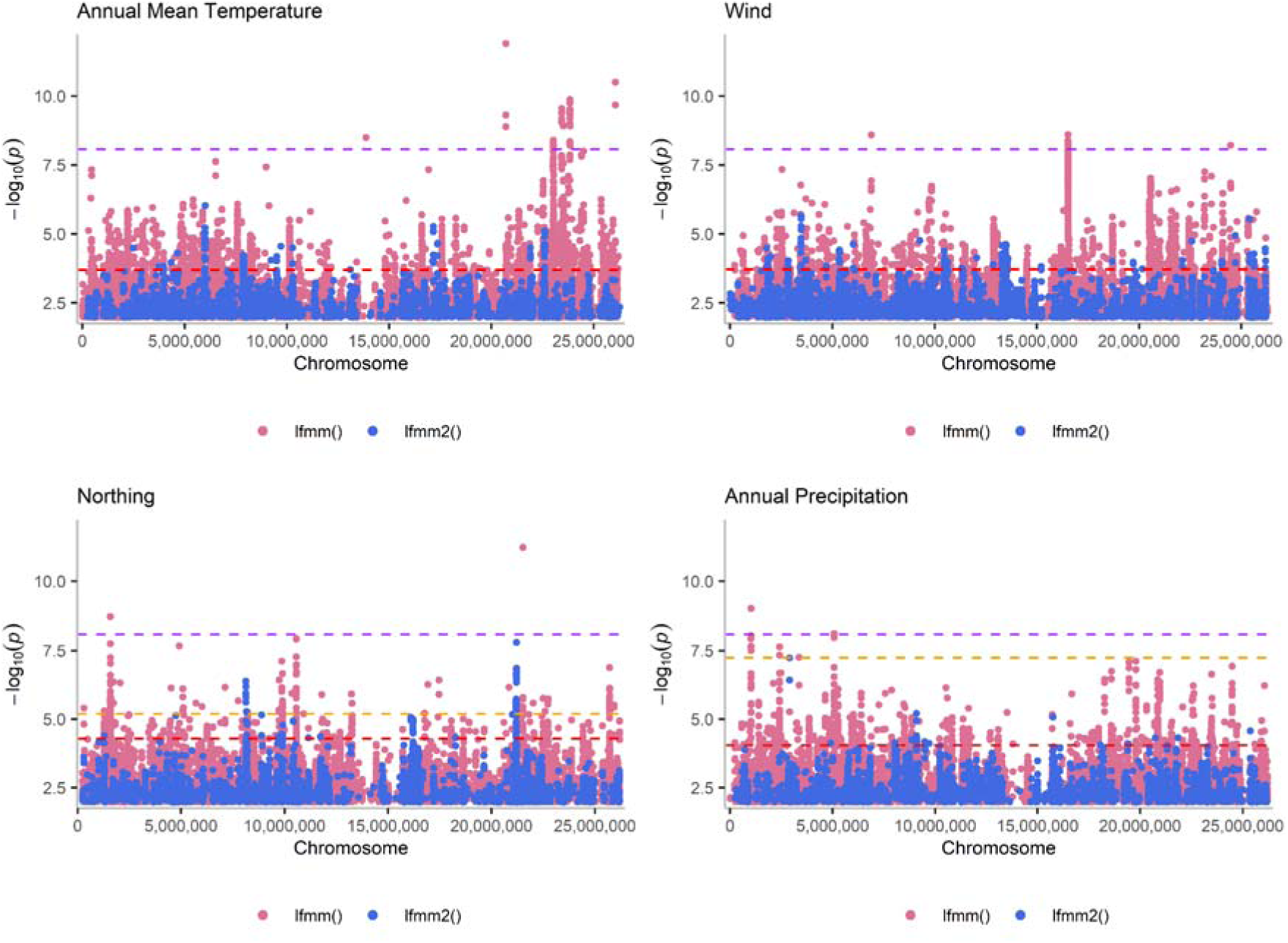
Manhattan plots comparing lfmm() and lfmm2() algorithms for a subset of variables. Analysis was performed on 1020 individuals at K=11 with genomic control using 420,165 SNPs present in chromosome nine to identify associations with 20 environmental, terrain and soil variables. Points are coloured according to the method, with pink for lfmm() and blue for lfmm2(). The dashed purple line represents the <u>genome</u>-wide Bonferroni significance threshold, while the red and orange lines indicate the chromosome level false discovery rate (FDR) of 5% following lfmm() and lfmm2() analyses, respectively.

