## Supplementary text for "Genetic basis of traits and performance in UK silver birch"

*Supplementary Text 1 - Effect of including related individuals on GEA*

Including ~800 individuals related to second-degree or higher had a reasonably small impact on the distribution of genomic variance (Supplementary Figure 19), with precipitation seasonality and mean temperature of the driest quarter still contributing the greatest proportions of total genomic variance (20.2% and 13.3% respectively, marginally greater than the dataset without related individuals). This suggests relatedness does not confound the partitioning of variance within the LFMM.

The largest changes were observed for soil phosphorus availability, isothermality, and precipitation of the warmest quarter, which showed slight reductions in both total genomic signal and redundancy when related individuals were included (Supplementary Figure 19). This suggests minor redistribution of signal across environmental variables rather than novel gain or loss of explanatory structure.

*Supplementary Text 2 - Annotation of FDR-significant SNPs following LFMM2 GEA*

Of the 117 SNPs identified in the GEA, nine fell within the coding regions of seven genes. These genes and their putative function are shown in Extended Table 2. This included two for mean temperature of the wettest quarter such as a pentatricopeptide repeat-containing protein. This same gene annotation was identified for a SNP associated with precipitation of the warmest quarter within a different chromosome hinting at a role in responding to temperature and precipitation. Pentatricopeptide repeat proteins function in post-transcriptional processing, and have a diverse array of functions including responding to abiotic and biotic stress, with links to reducing the generation of reactive oxygen species during stress (Wang and Tan 2025; Zsigmond et al. 2012).

Additional annotations for precipitation of the warmest quarter included a plant protein of unknown function with a pleckstrin-homology-like region. Such regions are linked with intra-cellular communication, cytoskeleton re-arrangements and lipid binding, with mutants showing differential response to disease stress (Aksoy et al. 2025; Yamaguchi and Kawasaki 2012; Tang et al. 2005; Vorwerk et al. 2007).

Finally, for northing, a SNP was identified within a NAD(P)H-ubiquinone oxidoreductase B1 protein. This functions in the electron transport chain of respiration, with mutants showing greater resilience against oxidative stress such as ammonium toxicity (Geisler et al. 2007; Podgórska et al. 2018). A further 25 gene annotations were identified in proximity to climate-associated SNPs (Supplementary Table 5).
